# Thermodynamic Stabilization of Human Frataxin

**DOI:** 10.1101/2023.09.08.556816

**Authors:** Reyes Núñez-Franco, Angel Torres-Mozas, Claudio D. Navo, Andreas Schedlbauer, Mikel Azkargorta, Ibon Iloro, Félix Elortza, Gabriel Ortega, Oscar Millet, Francesca Peccati, Gonzalo Jiménez-Osés

## Abstract

Recombinant proteins and antibodies are routinely used as drugs to treat prevalent diseases such as diabetes or cancer, while enzyme replacement and gene therapies are the main therapeutic intervention lines in rare diseases. In protein-based therapeutics, optimized *in vivo* stability is key as intrinsic denaturation and intracellular proteostatic degradation will limit potency, particularly in treatments requiring a sustained action, while clearance mechanisms may limit the amount of circulating protein. *In vivo* stability is ultimately correlated with the intrinsic thermodynamic stability of the biomolecule, but this is difficult to optimize because it often goes at the expense of reducing protein activity. Here, we have used *in silico* engineering approaches to thermodynamically stabilize human frataxin, a small mitochondrial protein that acts as an allosteric activator for the biosynthesis of Fe-S clusters, whose genetically-driven impairment results in a rare disease known as Friedreich ataxia. Specifically, we developed an efficient thermostability engineering computational approach that combines information on amino acid conservation, the Rosetta energy function, and two recent artificial intelligence tools – AlphaFold and ProteinMPNN – to produce thermodynamically stabilized variants of human frataxin. Such protein variants rescued the large destabilization exerted by well-known pathological mutations, with an increase over 20 °C in the melting temperature and a thermodynamic stabilization of more than 3 kcal·mol^-1^ at the physiological temperature. This stability surplus is translated into an enhanced resistance to proteolysis, while maintaining the protein fully functional. This case-study highlights the power of our combined computational approach to generate optimized variants, adequate for protein-based therapeutics.

## Introduction

Protein-based therapeutics offer advantages over small molecule drugs in terms of potency and the versatility of their functions (e.g. catalysis, signaling, transport); also, having evolved to carry out very specific roles, they tend to induce less side-effects.^1^ Since the first FDA approval of a recombinant protein (insulin, 1982), ∼350 protein-based therapeutics have been developed, including antibodies (e.g. Blinatumomab^2^ for the treatment of acute lymphoblastic leukemia), enzymes (e.g. Pegademase bovine^3^ for the treatment of adenosine deaminase deficiency), coagulation factors (e.g. Eptacog Alfa for the treatment of haemophilia^4^), protein hormones (e.g. insulin detemir for the treatment of diabetes mellitus^5^) and cytokines (e.g. interferon beta-1b for the treatment of multiple sclerosis^6^). Protein therapeutics classified as Group I encompass exogenous enzymes and regulatory proteins used to mitigate the consequences of the deficiency of a natural protein, either caused by its absence or lack of function due to pathological mutations.^7^ Classic examples include the use of insulin for the treatment of diabetes, or the use of pancreatic enzymes isolated from pigs for the treatment of cystic fibrosis.^7^

The success of protein-based therapeutics is often constrained by the stability of the biomolecular active principle. First, many examples show that an *in vitro* reduced thermodynamic stability is coupled with an impaired protein homeostasis *in vivo*^8^ due to premature unfolding and subsequent degradation of the active species in the proteasome^9^ or other protein clearance pathways.^10^ In principle, this can be overcome with protein variants that stabilize the folded conformation *at the physiological temperature*. For this protein *thermodynamic stabilization* strategy to work, the optimized variants should resemble its natural analogue as closely as possible, to retain the native function and to avoid triggering an immune response.^11^

Several strategies have been used for optimizing protein-based therapeutics by engineering their physicochemical properties. ^1,11^ These approaches seek to control protein homeostasis, to improve their bioavailability and to increase the time they remain functional in the body. Some examples include strategies can improve bioavailability by increasing the hydrodynamic diameter of the protein to reduce kidney filtration through the chemical modification of the protein surface with polymer grafts such as polyethylene glycol, or their fusion to larger, more soluble proteins.

In turn, thermodynamic stabilization and solubility can be achieved by engineering the amino acid sequence.^1^ This approach is particularly attractive because it does not require chemical modifications, thus improving reproducibility and yield. It has been successfully used to modulate, for example, the pharmacokinetics of insulin. Engineering insulin’s amino acid sequence to modulate its isoelectric point has provided the long-acting variant (24 h) glargine,^12^ where the slower absorption is the consequence of an increased precipitation resulting from an isoelectric point closer to physiological pH, as well as the faster acting variant glulisine, where shifting the isoelectric point in the opposite direction reduces the formation of oligomers and facilitates absorption.^13^ The mutagenesis approach has also been used on interferon β-1B, interleukin-2, and the human growth hormone to cite a few.^14–16^

Friedreich’s ataxia is a disease amenable to treatment with Group I protein-therapeutics. This autosomal recessive disorder with early onset (10–15 years) manifests with progressive ataxia, loss of tendon reflexes and dysarthria, with a prevalence of 1–2 per 50,000.^17^ All Friedreich’s ataxia patients present low levels of functional frataxin, a small iron storage protein that chaperones ferrochelatase and also acts as an allosteric modulator for iron-sulfur clusters biosynthesis by binding to the assembly complex in the mitochondria comprising proteins NFS1, ISD11, ACP, and ISCU (Figure 1). Ultimately, the disease leads to decreased maturation of iron–sulfur cluster proteins, including mitochondrial respiratory complexes, the proteins of the Krebs cycle, and proteins involved in DNA repair and replication.^17^ Human frataxin undergoes maturation from a 210 amino acid precursor protein, and after proteolytic processing into mitochondria is converted to the mature, functional form which comprises amino acids 81-210.^18^ From a structural point of view, it is composed of a flexible N-terminus, a conserved antiparallel β-sheet, involved in protein-protein recognition, two α-helices, and an unstructured C-terminus.^19^ Each frataxin unit can bind up to 7-10 Fe^2+^ ions through a conserved anionic surface (i.e., the acidic ridge, Figure 1).^20,21^ Fe^2+^-bound frataxin specifically binds protoporphyrin IX ─ which features the same binding epitope as ferrochelatase ─ with micromolar affinity.^22^

**Figure 1.**
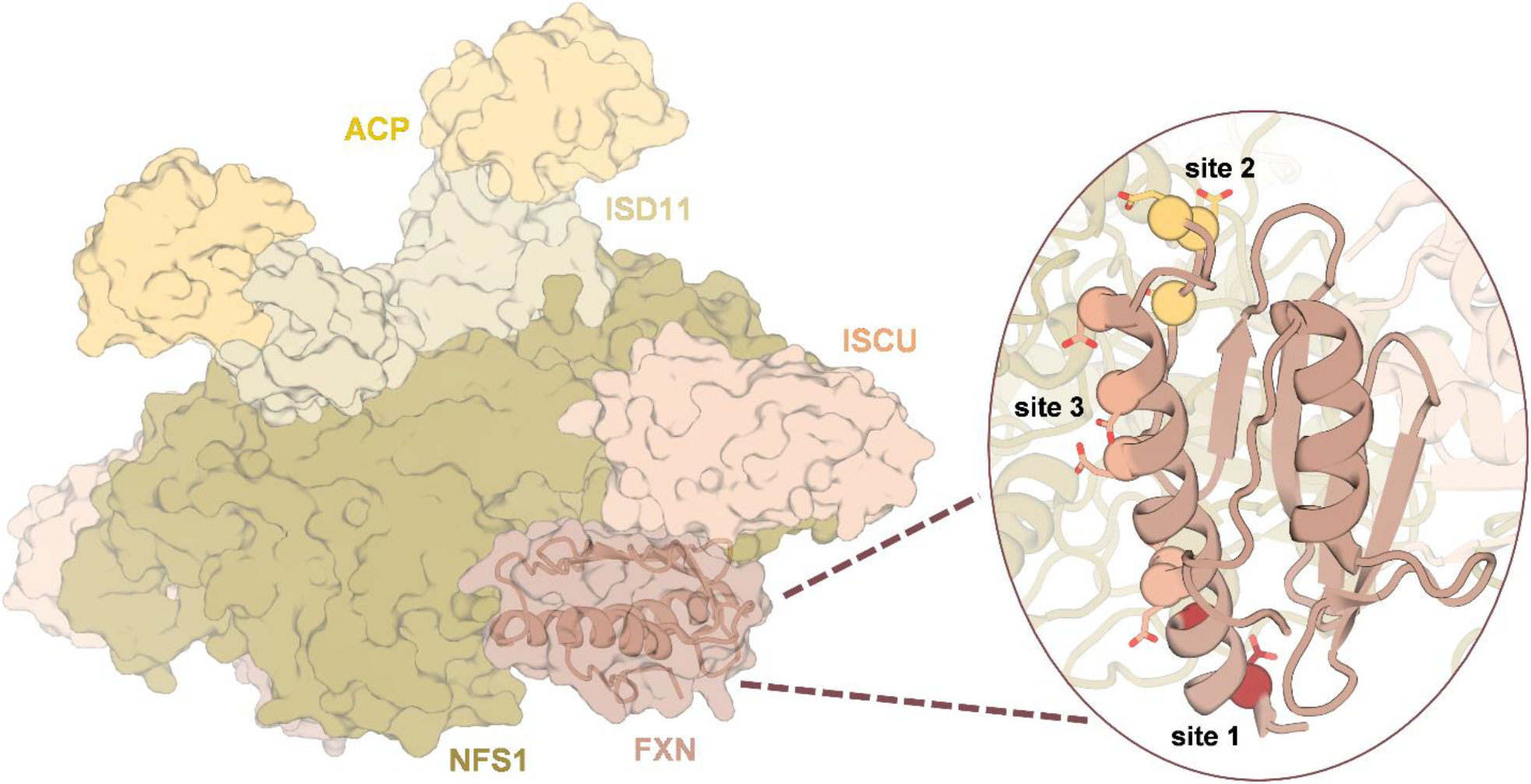
Cryo-EM structure of the frataxin-bound active human complex (PDB ID: 6NZU).^30^ The complex contains two copies of the NFS1-ISD11-ACP-ISCU-FXN hetero-pentamer. NFS1: cysteine desulfurase, mitochondrial (colored in green); ISD11: LYR motif-containing protein 4 (colored in light brown); ACP: acyl carrier protein (colored in orange); ISCU: iron-sulfur cluster assembly enzyme (colored in orange); FXN: frataxin (colored in dark red). A close-up view of frataxin is shown in the circle. The spheres represent the residues involved in iron binding, with each color representing one potential iron binding site, capable of binding more than one ion^31^ (site 1: E92, E96, site2: E121, D122, D124, site3: E100, E101, D104, E108, E111, D112, D115).

In over 96% of the patients, homozygous expansions of GAA trinucleotide repeat in intron 1 of the frataxin (FXN) gene are observed, triggering local chromatin changes leading to transcriptional repression of the gene (FXN mRNA levels reduced by 70–95%).^17^ In few other cases, Friedreich’s ataxia patients present a GAA expansion on one allele and a *missense mutation* on the other allele of the FXN gene; over 10 missense mutations have been identified, among them I154F and I198R, which decrease frataxin levels compromising its stability, solubility, and function.^23,24^ While no cure for this disease has been approved yet, there are indications that raising frataxin levels can inhibit and rescue the associated condition, opening opportunities for replacement approaches based on gene therapy.^25–27^ Alternatively, protein replacement therapy is a promising option provided that frataxin can be delivered to the mitochondria.^17^ In this regard, a Phase 2 study in humans is currently ongoing to explore the use as therapeutic of a fusion protein combining FXN into CTI-1601 (Larimar Therapeutics, Inc. ClinicalTrials.gov Identifier: NCT05579691), a protein that relies on the cell-penetrating ability of the trans-activator of transcription (TAT) peptide for targeted delivery.^28,29^ In the context of improving protein replacement therapy alternatives, engineering the homeostasis of frataxin by increasing its thermodynamic stability, and solubility may improve its resistance to intracellular degradation and, eventually, its bioavailability, thus optimizing its therapeutic effect.

In this contribution, we focus on the structured section of mature human frataxin comprising residues 91–210, and use computational design to propose engineered thermostable variants of both the wild-type frataxin and the pathological single mutants I154F and L198R as candidates for protein replacement therapy. Advancements in Artificial Intelligence (AI) have recently provided programs capable of predicting highly accurate three-dimensional protein models from sequence (e.g. AlphaFold,^32^ RoseTTAFold,^33^ ESMFold^34^), which are progressively closing the sequence-structure gap^35^ and provide exciting opportunities for the development of novel *in silico* protein design and engineering approaches.^36–38^ Our group has recently combined AlphaFold structure ensembles and Rosetta energy predictions to develop a computational approach to predict protein thermostability changes upon mutation which has been validated on several families of enzymes engineered through directed evolution.^39^ In this work, we combine i) mutational hotspot selection from evolutionary information, ii) sequence sampling on the selected hotspots with ProteinMPNN^38^ ─ a deep learning approach for sequence optimization of a given fold ─ and iii) our AlphaFold/Rosetta-based approach for thermostability evaluation to design stabilized, biologically competent frataxin variants. These variants have been expressed in *Escherichia coli* and their melting temperatures (T_m_) characterized. All 26 tested designs show equal or higher thermostability and unfolding reversibility than the corresponding natural sequence. Variant FXN-10 achieves a large thermostabilization of ΔT_m_ = +23 °C over the wild-type. Remarkably, the same variant also shows enhanced thermodynamic stabilization at physiological temperature, leading to increased resistance to proteolysis while maintaining the same ability to bind divalent cations and the FeS assembly complex as wild-type frataxin, representing a promising candidate for protein replacement therapy.

## RESULTS AND DISCUSSION

### Designs based on the consensus approach

Thermostability is an important readout of protein thermodynamic stability, and engineering this property aims at broadening the temperature range of stability and function of proteins and is crucial for the development of industrial and biomedical applications.^40^ The consensus approach has proven successful for the thermostabilization of enzyme families such as phytase-1 achieving stabilizations over 20 °C arising from the sum of small individual contributions.^41,42^ Hence, and in order to set a baseline for the thermostabilization that can be achieved through evolutionary information while preserving biological function, we first set out to design a consensus variant (FXN-01, Table 1) using the 7 mutational hotspots derived from a conservation analysis of the frataxin-enriched multiple sequence alignment MSA80 (alignment of sequences with at least 80% identity to human frataxin; positions with conservation below 60% selected as mutational hotspots; see Supporting Information for details), and selecting the most abundant amino acid at each position. Consensus variant FXN-01 was predicted by our AlphaFold/Rosetta-based protocol to be more stable than the wild-type and indeed presents a ΔT_m_ of +6.1 °C. The F120P single mutation – located in a loop region – carries the most thermostabilizing effect, as the remaining six mutations are nearly thermoneutral (variant FXN-02, Table 1; ΔT_m_ = +0.4 °C) and the single mutation alone (variant FXN-03, Table 1) induces a +5.1 °C change in T_m_. Regarding the beneficial F120P substitution, loop mutations to proline locally increase the rigidity of the protein in both the native and denatured states,^43^ and are a classic thermostabilizing strategy observed in thermophilic-mesophilic protein pairs.^44^

Inspection of the available of X-ray and NMR structures of wild-type frataxin and Friedreich’s Ataxia single mutants^19,30,45–47^ shows a flexible region encompassing residues 136–141 that is in direct contact with Phe120 (Figure 2a). Accordingly, AlphaFold ensembles of wild-type frataxin carry information on this flexibility showing significant population of two different backbone conformations for residues 138 and 139 (Figure 2a), confirming emerging reports on the ability of AlphaFold to predict not only static but also dynamic protein features.^39,48^ In the ensembles obtained for FTX-03, one of those backbone conformations is almost exclusively selected, suggesting that the F120P mutation stabilizes not only of the loop region where it is inserted, but also of the neighboring 137–140 region. Of note, this selected conformation is more stable in terms of Rosetta energy, particularly in FXN-03 (Figure 2b).

**Figure 2.**
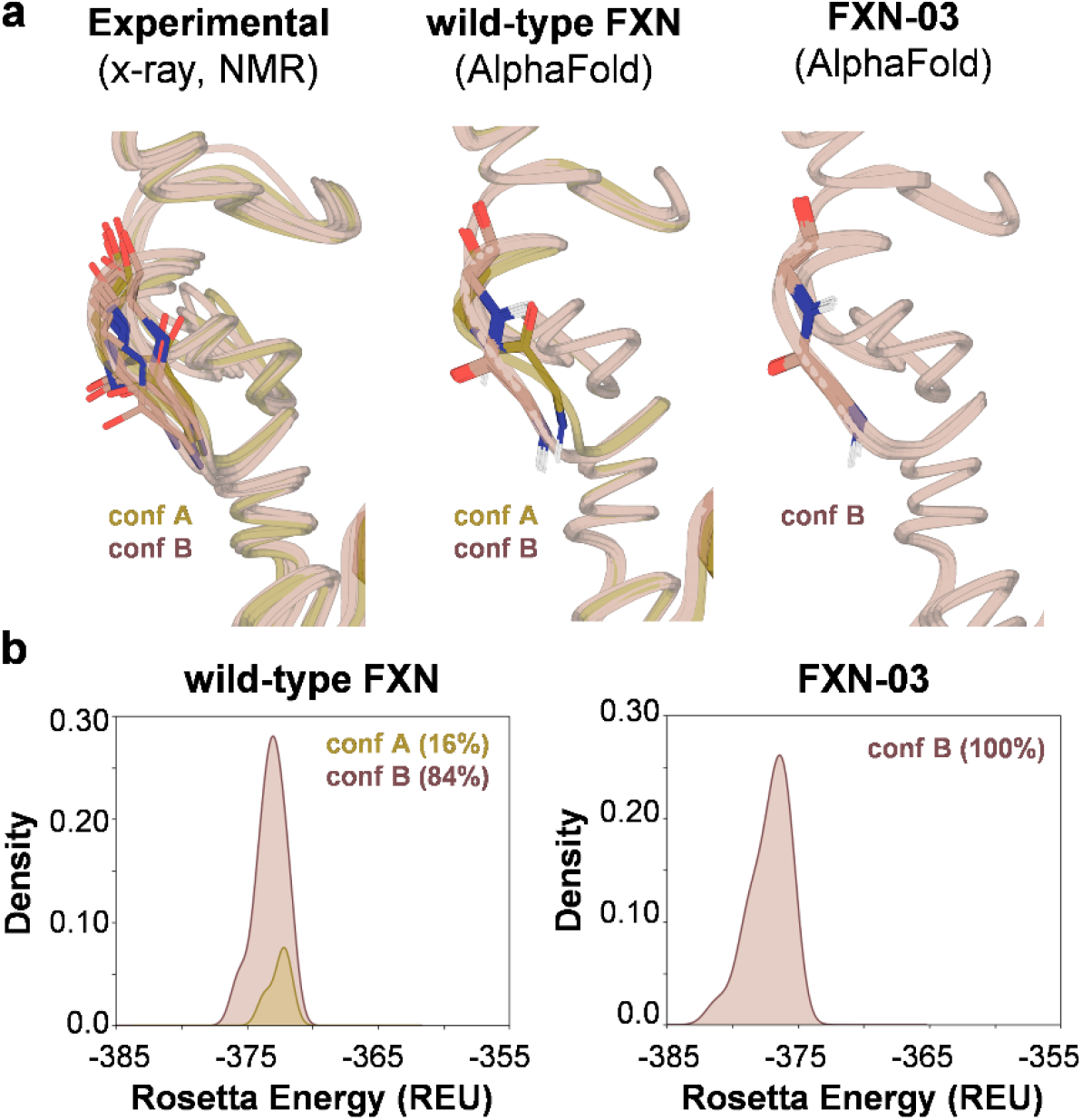
a) Flexible region between residues 137 to 140 in a) an overlay of NMR and X-ray structures (PDB ID: 1EKG, 3S4M, 3S5D, 3S5E, 3S5F, 3T3J, 3T3L, 3T3T, 3T3X, 1LY7, 6NZU) of different frataxin variants, and AlphaFold ensembles (25 structures) of wild-type frataxin and F120P single mutant FXN-03. While two backbone conformations are observed for residues 138 and 139 in both the experimental and AlphaFold predicted structures for wild-type frataxin, only one conformation is observed for FXN-03. b) Normalized kernel density estimates of the Rosetta energy distributions calculated for each variant in the two conformations. REU: Rosetta energy units.

Intrigued by the evolutionary significance of the highly conserved F120P mutation, we performed a phylogenetic analysis of the frataxin-enriched MSA80 alignment which clearly identified this substitution, together with two additional mutations present in consensus variant FXN-01 (S160T and T191S), as branch points between different orders of mammals and amphibians/reptilians/birds (see Supporting Information). Of note, by combining these mutations into the triple mutant FXN-04, predicted by our protocol to be at least as stabilizing as FXN-01, a further increase in T_m_ of +8.6 °C over the wild-type was achieved.

### ProteinMPNN sequence design

Recently, a very fast and robust deep learning–based protein sequence approach (ProteinMPNN) has been developed by the Baker lab, which achieves very high protein sequence recovery on native backbones without carrying out large scale sidechain packing calculations.^38^ By sampling with ProteinMPNN the same 7 mutational hotspots found in the MSA80 dataset (see Supporting Information), three more variants were produced: FXN-05 and FXN-06 (5 mutations each) predicted to be the two most stable sequences, and FXN-07 as the most frequent solution (5 mutations) (Table 2). All three designs showed increased T_m_ over wild-type FXN between +2.5 and +7.3 °C, matching or even improving the thermostabilization achieved by the consensus approach.

**Table 2.** Summary of the thermostabilized designs for wild-type frataxin, with indication of the method used for hotspot selection, the number of mutational hotspots (N. hotspots), the approach followed to generate the mutations, the name of the variant, the number of mutations (N. mutations), the difference in melting temperature (∆T_m_), and the predicted unfolding free energy change (∆∆*G*^*calc*^) (REU: Rosetta energy units). All quantities are referred to wild-type frataxin. The identity of the mutations for each variant are presented in Table S10. See Supporting Information for the definition of multiple sequence alignments MSA80, MSA70 and MSA60. All measured T_m_ values are summarized in Table S11.

| Hotspot detection | N. hotspots | Mutation approach | Variant | N. mutations | $\Delta T_m$ (°C) | $\Delta\Delta G^{calc}$ (REU) |
| --- | --- | --- | --- | --- | --- | --- |
| Per-residue conservation $\leq 60\%$ on MSA80 | 7 | Consensus | FXN-01 | 7 | 6.1<sup>a</sup> | 8.2 |
|  |  |  | FXN-02 | 6 | 0.4<sup>a</sup> | 3.1 |
|  |  |  | FXN-03 | 1 | 5.1 | 4.1 |
|  |  |  | FXN-04 | 3 | 8.6 | 7.7 |
|  |  | ProteinMPNN | FXN-05 | 5 | 7.3 | 11.9 |
|  |  |  | FXN-06 | 5 | 4.8 | 10.6 |
|  |  |  | FXN-07 | 5 | 2.5 | 9.0 |
| Per-residue conservation $\leq 60\%$ on MSA70 | 11 | ProteinMPNN | FXN-08 | 9 | 11.9 | 12.2 |
| Per-residue conservation $\leq 60\%$ on MSA60 | 26 | ProteinMPNN | FXN-09 | 16 | 12.7 | 14.3 |
| Shannon entropy (SE > 0.65) on MSA80 | 20 | ProteinMPNN | FXN-10 | 13 | 23.3 | 12.7 |

<sup>f</sup>The difference in melting temperature ( $\Delta T_m$ ) for these variants is calculated with respect to non-6xHis-tagged wild-type frataxin.

To ascertain whether a more extensive sequence sampling can lead to better thermostabilization, we sampled with ProteinMPNN the larger pools of hotspots generated from the MSA70 and MSA60 sequence alignments (sequences with a minimum 70% and 60% sequence identity to human frataxin, where 11 and 26 positions show conservation below 60%, respectively, and are set as mutable; see Supporting Information), obtaining FXN-08 (9 mutations) and FXN-09 (16 mutations) as the variants predicted by our AlphaFold/Rosetta protocol to be the most stable from each pool, and more stable than all previous designs. With these variants, we achieved an even larger increase in T_m_ over the wild-type (+11.9 °C and +12.7 °C, respectively) (Table 2). These results strongly suggest that improved thermostabilization can indeed be achieved by more extensive sequence design. Of note, ProteinMPNN only mutates a fraction of the available hotspots to achieve a significant improvement of the target properties.^38^

An alternative way of exploring evolutionary information without imposing an arbitrary per-residue conservation threshold, is to compute the per-position Shannon entropy^49^ (SE) on a given MSA. SE not only accounts for the degree of conservation but also for the global distribution of amino acid identities across different sequences. Hence, those positions which vary more frequently and to more diverse amino acids across the MSA are characterized by the highest SE values, and can be selected as mutational hotspots (see Supporting Information). Sampling with ProteinMPNN the 20 mutational hotspots identified through Shannon entropy on MSA80 yielded FXN-10 (13 mutations) – predicted to be the most stable of this pool – which shows a sizeable +23.3 °C increase in T_m_ over the wild-type. (Table 2). Of note, our computational protocol produced a superstable frataxin variant by efficiently selecting only 13 mutations out of a relatively small mutational space (20 positions); importantly, this significant thermostability increase was achieved in a single design step, unlike the multiple rounds of extensive mutation required by iterative saturation mutagenesis or directed evolution,^50,51^ and providing thermostabilization comparable to the most successful computational approaches^52–54^ at a fraction of the computational cost (less than 1 hour per variant on a mid-range workstation, see Supporting Information).

### Rescue of pathological mutants I154F and L198R

Having succeeded in thermostabilizing wild-type FXN, we turned to thermostabilize two pathological single mutants, FXN-I154F and FXN-L198R, which induce large decreases in T_m_ over 10 °C. The role of pathological mutations in Friedreich’s ataxia has been widely discussed, as different mutations lead to specific phenotypes of the disease tuning to different extents the protein levels, localization and function.^55^ Full-length frataxin has an unstructured region at the N-terminus encompassing the first 41 amino-acids, and its processing to the mature, functional form (residues 81-210) takes place through at least one intermediate (residues 42–210).^55^ Pathological mutation I154F (ΔT_m_ = –12.9 °C, Figure S4) has been shown to decrease both mature frataxin levels and accumulation of the intermediate without impairing its association with mitochondria,^23^ as well as promoting precipitation upon iron binding due to an increased flexibility.^56^ On the other hand, pathological mutation L198R (ΔT_m_ = –20.5 °C, Figure S4), located in the C-terminal region, places a positive charge in an apolar environment and destabilizes the native state inducing a more dynamic behavior in the microsecond/millisecond timescale around the mutation.^21,55,57^

For each of these two pathological variants, eight designs were generated by i) transplanting either one or the three stabilizing mutations identified in the phylogenetic analysis (F120P, S160T, T191S), ii) resampling with ProteinMPNN the same 26 mutational hotspots sampled in the wild-type, and iii) transplanting the mutations of the most stable engineered variants of the wild-type (FXN-09 and FXN-10; the L198I mutation of FXN-09 was not transplanted to FXN-L198R in order to preserve the original pathogenicity). Designs FXN-11 to FXN-18 (derived from FXN-I154F) and FXN-19 to FXN-26 (derived from FXN-L198R), were therefore produced (Tables 3 and S8). Regarding the I154F mutation, five variants achieve at least +20 °C increase in T_m_, which not only fully compensates the destabilizing effect of the pathological mutation, but also rises the stability over that of the wild-type. Similar results were obtained for the pathological mutation L198R, where four variants show ΔT_m_ ≥ +24 °C. In both cases, the largest stabilizations were achieved by transferring the mutations from FXN-10 (variants FXN-18 and FXN-26), although resampling with ProteinMPNN (variants FXN-15/16 and FXN-23/24) achieved very similar results in terms of both thermostability (ΔT_m_ +20 to +24 ºC) and introduced mutations (96–98% sequence identity), as expected. Importantly, almost all our variants increase stability without decreasing folding reversibility; this is particularly remarkable for variants designed with ProteinMPNN, which in many cases even enhanced reversibility (Table S11, Figures S7 and S9). This feature is important for improving protein homeostasis *in vivo*, as irreversibly unfolded proteins are more prone to aggregation and degradation.

**Table 3.** Summary of the thermostabilized designs for FXN-I154F and FXN-L198R, with indication of the method used for hotspot selection, the number of mutational hotspots (N. hotspots), the method used to generate the mutations, the name of the variant, the number of mutations (N. mutations), the difference in melting temperature (∆T_m_), and the predicted unfolding free energy change (∆∆*G*^*calc*^) (REU: Rosetta energy units). All quantities are referred to the parent pathological mutant of each variant (FXN-I154F for designs FXN-11 to FXN-18 and FXN-L198R for designs FXN-19 to FXN-26). The identity of the mutations for each variant are presented in Table S10. See Supporting Information for the definition of multiple sequence alignments MSA80, MSA70 and MSA60. All measured T_m_ values are summarized in Table S11.

| Hotspot detection | N. hotspots | Mutation approach | Variant | N. mutations | $\Delta T_m$ (°C) | $\Delta\Delta G^{calc}$ (REU) |
| --- | --- | --- | --- | --- | --- | --- |
| I154F + mutations from FXN-04 |  |  | FXN-11 | 3 | 9.9 | 9.5 |
| I154F + mutations from FXN-03 |  |  | FXN-12 | 1 | 6.7 | 5.7 |
| Per-residue conservation $\leq 60\%$ on MSA60 | 26 | ProteinMPNN | FXN-13 | 16 | 20.3 | 8.8 |
|  |  |  | FXN-14 | 15 | 21.1 | 9.0 |
| Shannon entropy (SE > 0.65) on MSA80 | 20 | ProteinMPNN | FXN-15 | 11 | 21.1 | 12.5 |
|  |  |  | FXN-16 | 11 | 20.5 | 13.3 |
| I154F+ mutations from FXN-09 |  |  | FXN-17 | 16 | 16.4 | 14.0 |
| I154F+ mutations from FXN-10 |  |  | FXN-18 | 13 | 23.4 | 15.2 |
| L198R + mutations from FXN-04 |  |  | FXN-19 | 3 | 9.3 | 2.6 |
| L198R + mutations from FXN-03 |  |  | FXN-20 | 1 | 4.1 | 1.7 |
| Per-residue conservation $\leq 60\%$ on MSA60 | 26 | ProteinMPNN | FXN-21 | 12 | 19.8 | 10.5 |
|  |  |  | FXN-22 | 14 | 22.9 | 8.1 |
| Shannon entropy (SE > 0.65) on MSA80 | 20 | ProteinMPNN | FXN-23 | 13 | 23.7 | 11.7 |
|  |  |  | FXN-24 | 12 | 23.9 | 9.6 |
| L198R+ mutations from FXN-09 |  |  | FXN-25 | 15 | 16.6 | 15.1 |
| L198R + mutations from FXN-10 |  |  | FXN-26 | 13 | 24.1 | 12.3 |

Overall, these results show that our approach is able to allosterically rescue the pathological single mutations by extensive substitutions in different regions of the protein.

### Physical origin of improved thermostability

Figure 3a shows the CD spectra of wild-type frataxin and superstable variant FXN-10 at different temperatures. While the former is unfolded at 75 ºC, the latter maintains a secondary structure that closely matches the wild-type one even at such high temperature, showing that the mutations introduced in FXN-10 do not significantly alter the folded structure of the protein. The most stable variants share many common mutations (8 between FXN-08 and FXN-10; 6 between FXN-09 and FXN-10) including hydrophobic-to-charged (A114K, A188E/K, A204K), hydrophobic-to-polar (A187S), charged-to-polar (K171T/S), substitution to closely related amino acids (R97K, Y118F, S160T, T191S), interconversion of polar amino acids (S202H), and substitution to proline (F120P, discussed above) (Table S10). These types of superficial mutations to either charged amino-acids or proline are commonly observed in the thermophilic counterparts of mesophilic proteins.^44^ Accordingly, ProteinMPNN is known to favor the introduction of charged amino acids at low sampling temperatures, which may contribute to thermostabilization of the designed sequences.^38^ In fact, the four charged residues introduced in the most stable variant FXN-10 (Lys97, Lys114, Lys180, Glu188 and Lys204) create a dense network of salt bridges not present in wild-type frataxin (Figure 3b), in line with previous observations that optimization of charge-charge interactions at the protein surface is a mechanism for stabilization.^58–60^ Still, the theoretical isoelectric point (IP = 5.4) and total charge at pH=7.4 (*Z* = –8.1) of FXN-10 are nearly identical to those of the wild-type (IP = 5.2; *Z* = –8.1) (Table S10).

**Figure 3.**
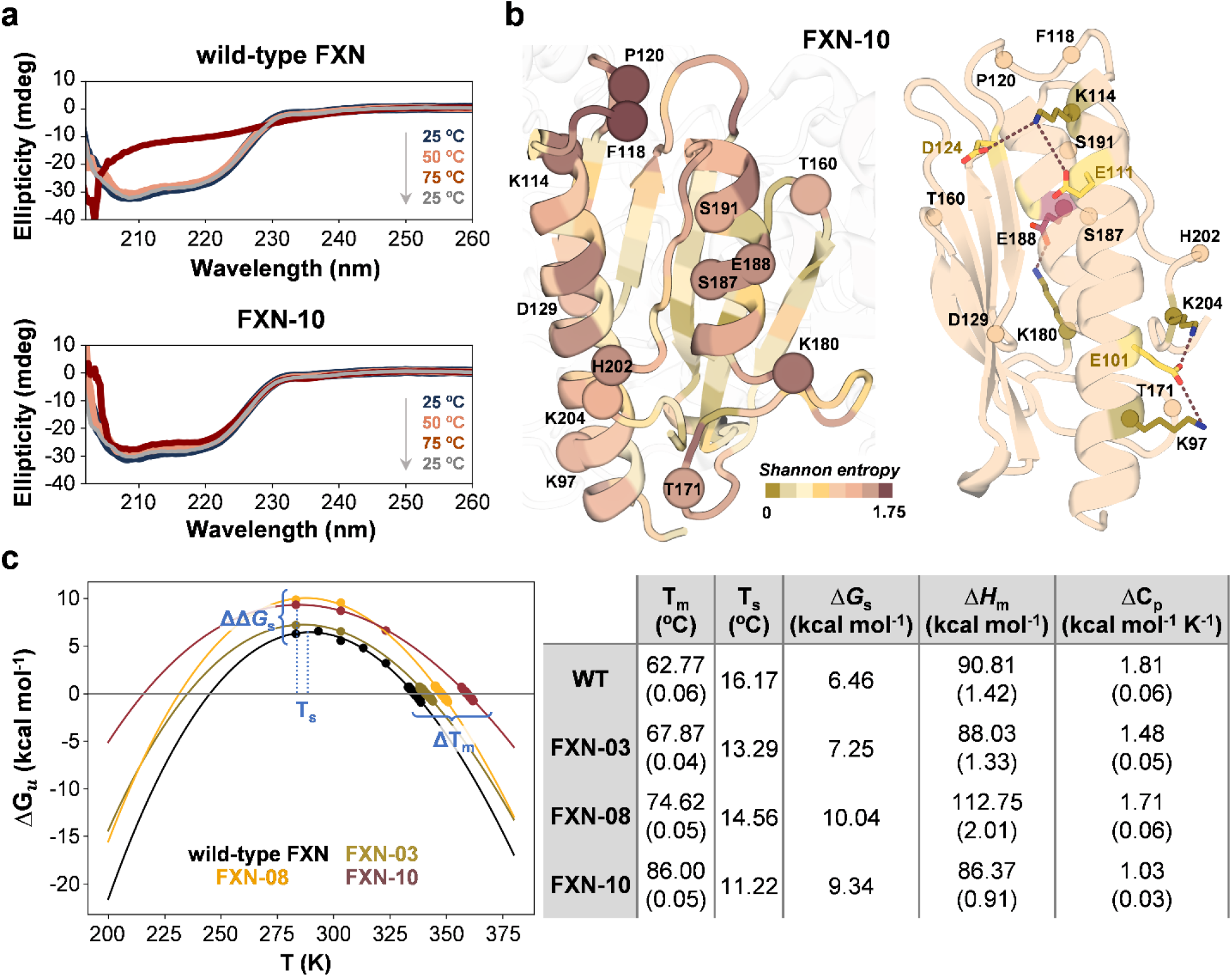
a) CD spectra at 25, 50, 75 ºC (heating phase) and 25 ºC (cooling phase) for wild type frataxin and design FXN-10. b) AlphaFold model for and FXN-10 colored according to Shannon entropy with indication of the mutated positions as spheres (left). Salt bridges introduced in thermostable design FXN-10 and not present in wild-type frataxin (right). The mutated charged residues engaged in new salt bridges are shown as green (K97, K,114, K180, K204) and magenta (E188) sticks. Non-mutated charged residues are shown as yellow sticks. c) Stability curves measured for wild-type frataxin, FXN-03, FXN-08 and FXN-10, together with the melting temperature (T_m_), temperature of maximum thermodynamic stability (T_s_), maximum thermodynamic stability (∆*G*_*s*_), enthalpy of unfolding at T_m_ (∆*H*_*m*_), and heat capacity of unfolding (∆*C*_*p*_) derived from these curves. Numbers in parenthesis indicate the standard errors arising from the fitting. Note that FXN-08 is the most *thermodynamically stable* variant (i.e., largest ∆∆*G*_*s*_), while FXN-10 is the most *thermostable* one (i.e., largest ΔT_m_).

In an attempt to gain further insights into the thermostabilization mechanisms operating in our designs, we measured the protein stability curves as a function of temperature^61^ of selected frataxin variants, characterizing the temperature dependence of the unfolding free energy ∆*G*_*u*_(*T*) of wild-type, FXN-03, FXN-08 and FXN-10 (Figure 3c and Supporting Information). The four variants show similar temperatures of maximal stability (T_s_ = 11–16 ºC) and a raised stability curve at all temperatures – as observed for the vast majority of mesophile-thermophile pairs^44,62–65^ – with variants FXN-08 and FXN-10 achieving a thermodynamic stabilization of ∆∆*G*_*s*_ ∼ 3 kcal mol^-1^. While the FXN-08 stability curve has essentially the same curvature as the wild-type one – implying similar heat capacities of unfolding (Δ*C*_p_) – and is shifted upwards due to a larger enthalpy of unfolding at T_m_ (∆*H*_*m*_), FXN-03 and FXN-10 show comparable ∆*H*_*m*_ and lower Δ*C*_p_ values than wild-type frataxin. For the variant showing the largest increase in T_m_, FXN-10, Δ*C*_p_ drops by 0.8 kcal mol^-1^ K^-1^ resulting in a substantial flattening of the curve, a stabilization mechanism observed in heat adaptation.^62,64,65^ Importantly, the fact that our variants feature higher stability at physiological temperatures supports the value of our approach to improve the homeostasis of the protein *in vivo*.

### Evaluation of *in silico* prediction accuracy

To evaluate the global accuracy and predictivity of our method, we compared the calculated (∆∆*G*^*calc*^) and experimentally measured changes in thermostability for the set of variants designed in this work. As described above, the curvature of the stability curve might not be the same for all variants, so the common approximation that changes in melting temperatures and unfolding free energies are lineally related^61^ (see Supporting Information), cannot be adopted in our case; on the other hand, a rigorous determination of the thermodynamic stabilization (∆∆*G*_*s*_) of the nearly 30 assayed variants from their stability curves is beyond the scope of this work. Therefore, a direct comparison between Rosetta and experimental energies is not possible, and hence we directly correlated ∆∆*G*^*calc*^ with ∆T_m_ values with respect to wild-type frataxin instead. Overall, we obtained a strong correlation between computed and experimental stabilities within a T_m_ range spanning over 40 °C (Figure 4), achieving better predictivity than popular force-field and machine-learning-based approaches.^66^

**Figure 4.**
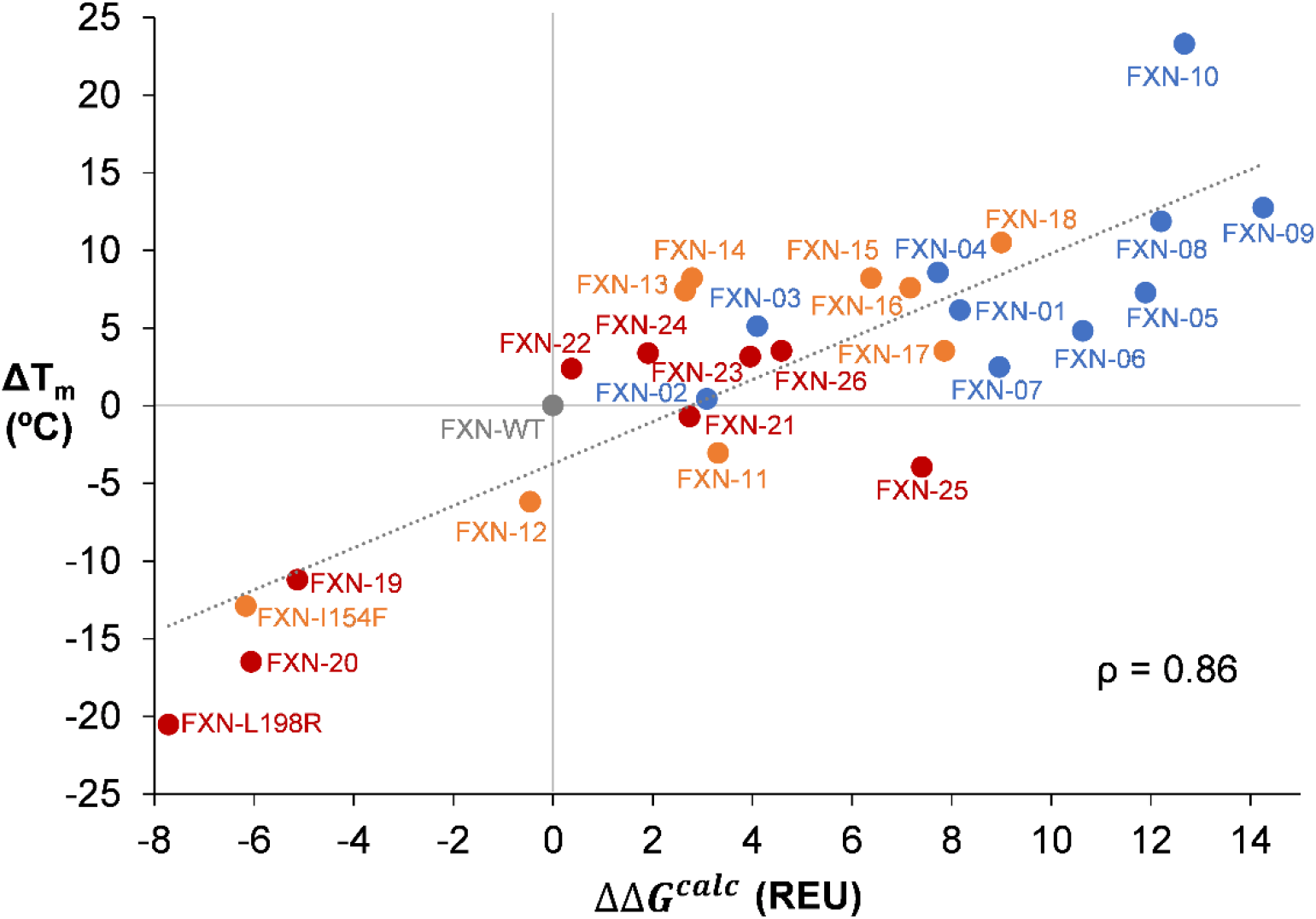
Correlation between theoretical ∆∆*G*^*calc*^ and experimental ΔT_m_ values for the 26 variants designed to stabilize wild-type frataxin (in blue) and the two pathological mutants I154F (in orange) and L198R (in red), using wild-type frataxin (in grey) as a reference. Positive and negative values indicate stabilization and destabilization, respectively. Each data point corresponds to a different protein variant. ρ: Pearson correlation coefficient. The dashed line indicates a linear regression with slope 1.35 and intercept –3.74.

Outliers from the generally good agreement are designs FXN-10 and FXN-25, whose ∆*T*_*m*_ are under- and overestimated, respectively. For FXN-10, the thermodynamic parameters derived from the stability curves suggest that the underestimation may arise from the flattening of the curve due to lower Δ*C*_p_, which differentially increases ∆*T*_*m*_ over ∆∆*G*_*s*_. In fact, for the thermodynamically characterized variants (Figure 3c), Rosetta energies correlate much better with the latter (Figure S11), suggesting that our approach might be best suited for predicting changes in *thermodynamic stability* (i.e., ∆∆*G*_*s*_) rather than *thermostability* (∆T_m_).

Interestingly, a decomposition of Rosetta energies shows that FXN-10 is also an outlier in terms of the contributions to the computed values. In particular, we detected notable differences in the relative importance of Rosetta’s *ref* term in the computed energy of each variant with respect to the wild-type. This term accounts for the contribution of the unfolded state to the unfolding free energy;^67^ since explicit calculations on the inherently complex and unknown unfolded state are prohibitively expensive and error prone, the *ref* term is structure-independent and assigns to each amino acid of the sequence a weight empirically determined to maximize native sequence recovery.^67^ In FXN-10, this contribution is much larger than in the other variants and dominates ∆∆*G*^*calc*^ (Figure S5), suggesting an important contribution of the unfolded state to the thermodynamic stabilization. Since Δ*C*_p_ values have been shown to correlate with changes in the solvent accessible surface area between unfolded and folded states (ΔSASA),^68^ we tentatively propose that the observed decrease in Δ*C*_p_ for FXN-10 might be due to a reduced SASA in its unfolded state compared to the wild-type. This hypothesis is supported by the small SASA difference (∼2%) computed for the *folded* states of the two proteins (Table S9). These results suggest that our AlphaFold/Rosetta-based protocol is not only is capable of correctly predicting the sign and relative intensity of thermostability changes upon mutation, but also provides qualitative insights into the underlying biophysical thermostabilization mechanisms.

### Stabilizing mutations confer improved resistance to proteolytic degradation

To evaluate the extent to which our variants would improve protein homeostasis in a biological environment, we measured their propensity to enzymatic degradation. Early studies on extremely thermophilic bacteria demonstrated that enzymes derived from these organisms seemed to be more resistant to proteolysis than the corresponding enzymes from mesophilic organisms,^69^ suggesting that this resistance may be a general feature of thermostable proteins.^70^ In fact, resistance to proteolytic degradation is used nowadays to assess the thermodynamic folding stability of large libraries of proteins.^71^ Accordingly, the most thermostable frataxin variant designed in this work, FXN-10 (∆T_m_ = 23.3 ºC), exhibited a notable resilience to proteolysis by trypsin when compared to the wild-type (Figure 5a and Supporting Information). This improved resistance to proteolytic degradation will increase the lifetime of the protein in biological environments, further supporting the use of our engineered frataxin variants as therapeutics.

**Figure 5.**
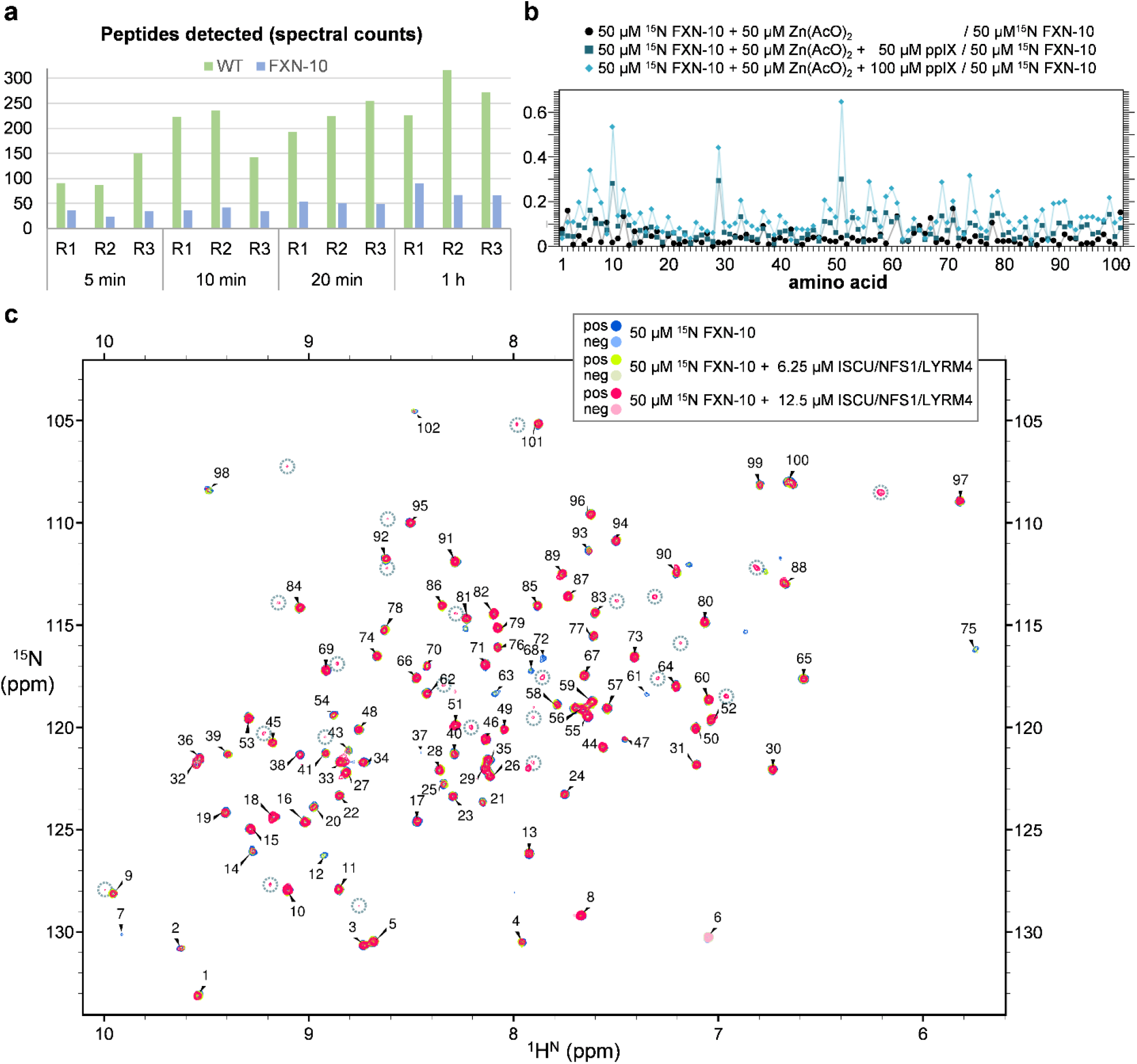
a) Protein stability assay by trypsin degradation and mass spectrometry analysis shows decreased proteolysis for engineered variant FXN-10 compared to wild-type frataxin. Spectral counts refer to peptide-spectrum match (PSM), i.e., peptides identified after protein digestion. b) CSP observed for u-^15^N,^13^C-labelled FXN-10 in the presence of Zn^2+^ and protoporphyrin IX (ppIX). c) 800 MHz 2D ^15^N-sf-HMQC spectra of u-^15^N,^13^C-labelled FXN-10 in the presence of increasing amount of FeS assembly complex. Residue numbers are arbitrary due to lack of complete assignment of the protein.

### Stabilizing mutations preserve biological function

Besides being thermally and enzymatically stable, an engineered protein must retain the biological activity of interest in order to be used as a potential therapeutic. By performing a series of NMR experiments developed in our laboratory,^22^ we verified that superstable FXN-10 is able to bind Zn^2+^ ions (as a proxy for Fe^2+^) and protoporphyrin IX (ppIX) – which, when bound to metal-loaded frataxin shares the binding epitope with ferrochelatase – as well as to engage in protein-protein interactions with the iron-sulfur assembly cluster (Figure 5b and Supporting Information). Measurement of chemical shift perturbation (CSP) upon addition of Zn^2+^ and ppIX show changes upon binding consistent with those observed for wild type frataxin.^22^ Also, interaction of frataxin with the iron-sulfur assembly complex shows the appearance of a new set of signals (Figure 5c and Supporting Information), in line with the slow exchange regime of the interaction. This is again in agreement with our prior observations for wild-type frataxin, proving the biological competency of the designed, stable variant.

## Conclusions

Highly stabilized human frataxin variants have been designed *in silico* by combining conservation-based mutation hotspot selection from evolutionary information with deep-learning-based sequence sampling using ProteinMPNN, and stability evaluation using AlphaFold ensembles and the Rosetta energy function. These predictions have been made in one single step, with minimal computational cost, and with a 100% success ratio (i.e., all designed variants exhibit improved thermostability compared to the wild-type, with varying levels of enhancement). The most stable design FXN-10, whose stabilization is largely associated to a reduced ∆*C*_*p*_ of unfolding resembling observed heat adaptation mechanisms, is highly resistant to proteolytic degradation and retains the wild-type’s ability to bind metal ions and interact with the FeS assembly complex. All these beneficial properties confer this variant a great potential for being used in protein replacement therapy for Friedreich’s ataxia. We believe these studies might establish a generalizable, rational strategy for the design of proteins with improved properties for its use as therapeutics, as well as in other biotechnological applications.

## ASSOCIATED CONTENT

### Supporting Information

Detailed methods, characterization and computational data, and additional figures (PDF)

## ACKNOWLEDGMENTS

This research was supported by the Agencia Estatal Investigacion of Spain (AEI; grants PID2021-125946OB-I00, PID2021-124171OB-I00, PDC2022-133725-C22, PDC2022-133677-I00, CEX2021-001136-S and a predoctoral fellowship to R.N.F.) and the Partnership for Advanced Computing in Europe (PRACE-ICEI, reference icei-prace-2021-0001). F.P. thanks the Ministerio de Economía y Competitividad for a Juan de la Cierva Incorporación (IJC2020-045506-I) research contract. G.O.Q. acknowledges support from Ikerbasque, the Basque Foundation for Science, and from the Ramón y Cajal Program (AEI; grant RYC2021-031074-I).

