## Supporting Information for "Thermodynamic Stabilization of Human Frataxin"

[†]Basque Research & Technology Alliance (BRTA), CIC bioGUNE, Bizkaia Technology Park, Building 800, 48160 Derio, Spain.

[‡]Ikerbasque, Basque Foundation for Science, 48013 Bilbao, Spain.

[⁋]ATLAS Molecular Pharma, Bizkaia Science and Technology Park, 48160 Derio, Spain.

[‖]Biomedical Research Network on Hepatic and Digestive Diseases (CIBEREHD), Instituto de Salud Carlos III, 28029 Madrid, Spain.

**CONTENTS**

1. Mutational hotspots selection ........ S2–S7

2. Phylogenetic analysis ........ S8–S10

3. Pathological mutations ........ S11

4. Sequence sampling with ProteinMPNN ........ S11

5. Prediction of relative thermostability of designed variants ........ S11–S13

6. Protein expression and purification ........ S14

7. Amino acid sequences of expressed frataxin variants ........ S15–S17

8. Circular dichroism (CD) spectroscopy ........ S18–S21

9. Melting temperature (Tm) measurement ........ S22–S28

10. Stability curve determination ........ S29–S31

11. Proteolytic resistance assay ........ S32

12. Mass spectrometry data ........ S33–S56

13. Binding of frataxin variants to $Zn^{2+}$/ppIX and FeS assembly complex ........ S57–S66

14. References ........ S67–S68

### 1. Mutational hotspots selection

Different sets of mutable amino acids were chosen from various conservation analyses performed on the same multiple sequence alignment (MSA). This MSA was generated by the *jackhmmer*[1] search of the wild-type target frataxin sequence (residues 91-210) against the UniRef90 database[2,3] at the initial stage of the AlphaFold version 2.3.0 structure prediction (database date: 2022-01; database size: 140,403,594; flags: --F1 0.0005 --F2 5e-05 --F3 5e-07 --incE 0.0001 -E 0.0001 -N 1), and contains 1999 sequences. The distribution of sequence identities, computed after removing the gaps from the target frataxin sequence (Figure S1) has a maximum at around 40% identity. At a higher identity (over 60%), we identified a set of clusters mostly represented by frataxin homologs, with the occasional appearance of ferroxidase and phosphatidylinositol 4-phosphate 5-kinase (type-1 beta isoform) homologs (Table S1). Of note, yeast frataxin has shown ferroxidase activity.[4,5]

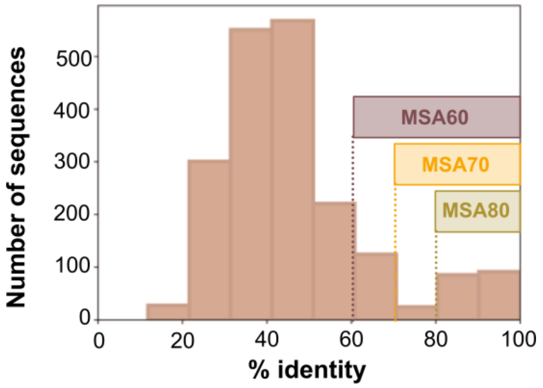


**Figure S1.** Distribution of sequences in the original Uniref90 multiple sequence alignment for human frataxin (1999 sequences), and derived subsets at 60% (MSA60), 70% (MSA70) and 80% (MSA80) identity, used for mutational hotspots identification.

Aiming to obtain mutational pools of different sizes, this MSA was then filtered three times based on sequence identity with the target human frataxin sequence after removing gaps (60%, 70%, and 80%), yielding MSA60, MSA70 and MSA80, respectively (Table S1). For each reduced alignment, we selected as mutational hotspots positions with conservation below 60%, with the rationale that least conserved positions are the most amenable to mutation without compromising structural integrity and associated function.[6–9] In this way, none of the selected mutational hotspots was located on the β-sheet plane responsible for binding to ISCU (iron-sulfur cluster assembly enzyme), where the pathological mutation W155G has been shown to delete the desulfurase activity of the supercomplex for iron–sulfur cluster assembly.[10]

**Table S1.** Composition of the multiple sequence alignments used for identification of mutational hotspots. The different protein categories are extracted from their definition in the FASTA files downloaded from UniProt (https://www.uniprot.org).

| | Uniref90 | | MSA60 | | MSA70 | | MSA80 | |
|---|---|---|---|---|---|---|---|---|
| **Category** | **Number of sequences** | **Percentage of sequences** | **Number of sequences** | **Percentage of sequences** | **Number of sequences** | **Percentage of sequences** | **Number of sequences** | **Percentage of sequences** |
| Frataxin | 576 | 29% | 316 | 94% | 196 | 93% | 170 | 94% |
| Frataxin-like | 12 | 1% | 0 | 0% | 0 | 0% | 0 | 0% |
| Ferroxidase | 1041 | 52% | 2 | 1% | 0 | 0% | 0 | 0% |
| Iron-sulfur_cluster | 75 | 4% | 0 | 0% | 0 | 0% | 0 | 0% |
| Iron_donor_protein | 82 | 4% | 0 | 0% | 0 | 0% | 0 | 0% |
| Other_iron-related | 9 | 0% | 0 | 0% | 0 | 0% | 0 | 0% |
| Uncharacterized | 103 | 5% | 7 | 2% | 7 | 3% | 5 | 3% |
| Other | 101 | 5% | 10 | 3% | 7 | 3% | 5 | 3% |
| **TOTAL** | **1999** | **100%** | **335** | **100%** | **210** | **100%** | **180** | **100%** |

Additionally, a fourth set of mutational hotspots was obtained from frataxin-enriched MSA80 dataset by scoring each position of frataxin orthologs using Shannon entropy[11] (SE) and excluding gaps:

$$SE(position) = -\sum_i f_i \ln f_i \qquad \text{eq. 1}$$

where $f_i$ is the frequency of each amino acid at the given position. Variable positions return positive SE values, while fully conserved positions return an SE value of zero. In this case, we arbitrarily selected the 20 highest entropy positions (SE > 0.65) to allow at most 15% of the sequence to mutate. Fourteen of these positions are in common with the hotspots identified from the MSA60 dataset using a per-residue conservation ≤ 60% threshold (see Table S2 for further comparison between the two approaches).

**Table S2.** Number of sequences and mutational hotspots obtained after applying different consecutive filters to the initial UniRef90 MSA obtained for the target human frataxin sequence (1999 sequences) during AlphaFold prediction of wild-type frataxin: 1) sequence identity and 2) individual amino acid conservation or Shannon entropy. Common hotspots across the different datasets are shown in red (found in the four mutation pools), orange (found in at least three of the mutation pools) and purple (found through either per-residue conservation or Shannon entropy approaches).

| **Multiple sequence alignment** | **Sequence identity threshold** | **Number of sequences** | **Per-residue conservation threshold** | **Number of mutational hotspots** | **Mutational hotspots** |
|---|---|---|---|---|---|
| MSA60 | 60% | 335 | 60 % | 26 | 93, 94, 97, 105, 108, 114, 116, 118, 120, 121, 140, 160, 171, 172, 184, 187, 188, 190, 192, 193, 194, 197, 198, 202, 204, 208 |
| MSA70 | 70% | 210 | 60 % | 11 | 97, 118, 120, 160, 171, 187, 188, 191, 192, 202, 204 |
| MSA80 | 80% | 180 | 60 % | 7 | 120, 160, 171, 187, 188, 191, 192 |
| **Multiple sequence alignment** | **Identity threshold** | **Number of sequences** | **Shannon entropy threshold** | **Number of mutational hotspots** | **Mutational hotspots** |
| MSA80 | 80% | 180 | 0.65 | 20 | 97, 108, 114, 118, 120, 129, 134, 140, 152,160, 171, 179, 180, 187, 188, 190, 191, 192, 202, 204 |

**Table S3.** Per-residue conservation (*cons*) of the mutational hotspots derived from the MSA60 dataset (335 sequences, conservation threshold 60%), sorted by amino acid type. For clarity, only values 5 ≤ *cons* ≤ 60% are shown. Cells are colored according to their conservation value (0%: white; 100%: red). A sequence logo representation of these mutational hotspots is shown below.

| | **93** | **94** | **97** | **105** | **108** | **114** | **116** | **118** | **120** | **121** | **140** | **160** | **171** | **172** | **184** | **187** | **188** | **190** | **192** | **193** | **194** | **197** | **198** | **202** | **204** | **208** |
|---|---|---|---|---|---|---|---|---|---|---|---|---|---|---|---|---|---|---|---|---|---|---|---|---|---|---|
| **G** | | | | | | | | | 24 | | | | | | | | | | | | | | | | | |
| **A** | 27 | 27 | | 36 | | 54 | | | | 14 | | | | | | 38 | 13 | | 13 | 58 | | | | | 38 | |
| **P** | | | | | | | | | 25 | | | | | | | | | | | | | | | | 18 | |
| **V** | | | | | | | | | | | | | | | | | | | 7 | | | | 5 | | 8 | |
| **L** | | | | | | | | | 13 | | 44 | | | | | | | 53 | | | 59 | | 59 | | 5 | |
| **I** | | | | | | | | | | | | | | | | | | | 14 | 30 | | | 17 | | 20 | |
| **M** | | | | | | 7 | | | | | 21 | | | | | | | | | | | | 15 | | | |
| **F** | | | | | | | | 52 | 22 | | | | | | | | | 40 | | | 36 | | | | | |
| **Y** | | | | | | | | 37 | | | | | | | | | | | | | | | | | | |
| **W** | | | | | | | | | | | | | | | | | | | | | | | | | | |
| **H** | | | | | | | | | | | 27 | | | 6 | | | | | | | | | | 10 | | |
| **S** | | | | 58 | | | | | 7 | | | 43 | | 6 | | 47 | | | | | | | | 41 | | |
| **T** | 59 | 55 | | | | 31 | | | | | | 48 | | | | | 19 | | 16 | | | | | 5 | | |
| **C** | | | | | | | | | | | | | | | | | | | | | | | | 13 | | |
| **N** | | | | | | | | | | | | | | 55 | | | | | | | | 22 | | | | |
| **Q** | | | | | | | 7 | | | | | | | | 25 | | | | | | | | | | | |
| **R** | | | 42 | | | | | | | | | | 24 | 22 | | | 8 | | | | | | | | | |
| **K** | | | 53 | | | | 58 | | | 8 | | | 38 | | | | 41 | | 21 | | | 57 | | | | 54 |
| **E** | | | | | 59 | | 28 | | | 54 | | | 25 | | 56 | | | | 10 | | | 5 | | | | |
| **D** | | | | | 36 | | | | | | | | | | 8 | | | | | | | | | 10 | | |

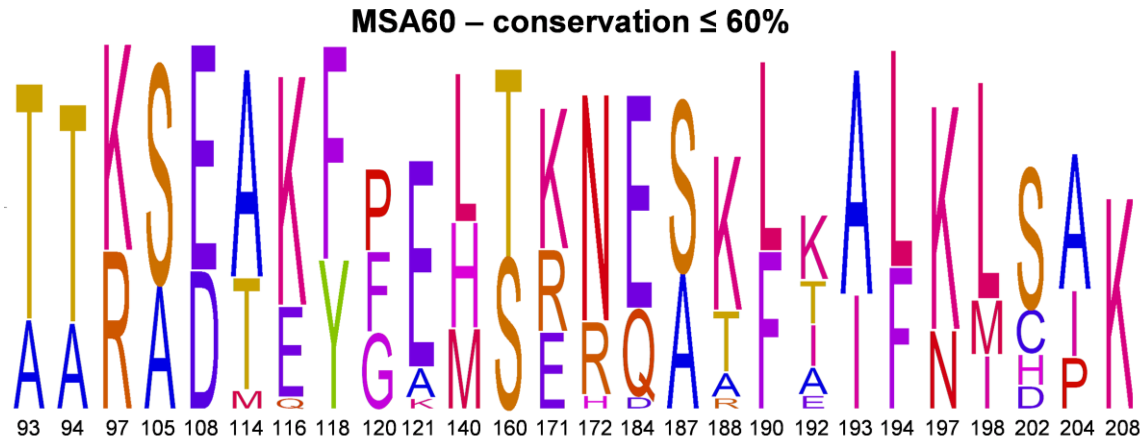

**Table S4.** Per-residue conservation (*cons*) of the mutational hotspots derived from the MSA70 dataset (210 sequences, conservation threshold 60%), sorted by amino acid type. For clarity, only values $5 \leq cons \leq 60\%$ are shown. Cells are colored according to their conservation value (0%: white; 100%: red). A sequence logo representation of these mutational hotspots is shown below.

| | **97** | **118** | **120** | **160** | **171** | **187** | **188** | **191** | **192** | **202** | **204** |
|---|---|---|---|---|---|---|---|---|---|---|---|
| **G** | | | | | | | | | | | |
| **A** | | | | | | 56 | 17 | | 11 | | 56 |
| **P** | | | 35 | | | | | | | 7 | |
| **V** | | | | | | | | | | | 10 |
| **L** | | | 19 | | | | | | | | |
| **I** | | | | | | | | | | | 22 |
| **M** | | | | | | | 7 | | | | |
| **F** | | 40 | 32 | | | | | | | | |
| **Y** | | 55 | | | | | | | | | |
| **W** | | | | | | | | | | | |
| **H** | | | | | | | | | | | |
| **S** | | | 9 | 56 | | 28 | | 47 | | 60 | |
| **T** | | | | 37 | | 5 | 29 | 52 | 21 | 7 | |
| **C** | | | | | | | | | | 18 | |
| **N** | | | | | | | | | | | |
| **Q** | | | | | | | | | 7 | | |
| **R** | 57 | | | | 36 | | 9 | | | | |
| **K** | 40 | | | | 53 | | 31 | | 30 | | |
| **E** | | | | | 5 | 6 | | | 16 | | |
| **D** | | | | | | | | | | | |

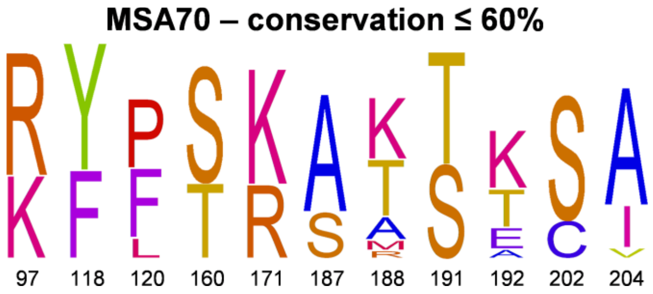

**Table S5.** Per-residue conservation (*cons*) of the mutational hotspots derived from the MSA80 dataset (180 sequences, conservation threshold 60%), sorted by amino acid type. For clarity, only values 5 ≤ *cons* ≤ 60% are shown. Cells are colored according to their conservation value (0%: white; 100%: red). A sequence logo representation of these mutational hotspots is shown below.

| | **120** | **160** | **171** | **187** | **188** | **191** | **192** |
|---|---|---|---|---|---|---|---|
| **G** | | | | | | | |
| **A** | | | | 59 | 17 | | 12 |
| **P** | 34 | | | | | | |
| **V** | | | | | | | |
| **L** | 19 | | | | | | |
| **I** | | | | | | | |
| **M** | | | | | 7 | | |
| **F** | 36 | | | | | | |
| **Y** | | | | | | | |
| **W** | | | | | | | |
| **H** | | | | | | | |
| **S** | 7 | 60 | | 25 | | 43 | |
| **T** | | 36 | | 5 | 31 | 57 | 23 |
| **C** | | | | | | | |
| **N** | | | | | | | |
| **Q** | | | | | | | 8 |
| **R** | | | 39 | | 10 | | |
| **K** | | | 53 | | 28 | | 34 |
| **E** | | | 5 | 5 | | | 16 |
| **D** | | | | | | | |

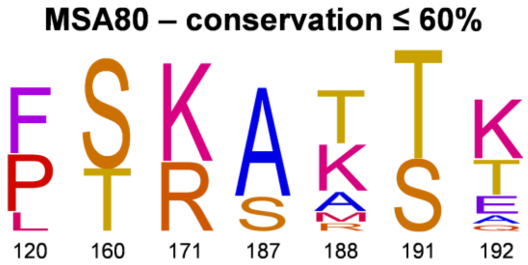

**Table S6.** Per-residue conservation (*cons*) of the mutational hotspots derived from the MSA80 dataset (180 sequences, Shannon entropy, SE > 0.65), sorted by amino acid type. For clarity, only values *cons* ≥ 5% are shown. Cells are colored according to their conservation value (0%: white; 100%: red). A sequence logo representation of these mutational hotspots is shown below.

| | **97** | **108** | **114** | **118** | **120** | **129** | **134** | **140** | **152** | **160** | **171** | **179** | **180** | **187** | **188** | **190** | **191** | **192** | **202** | **204** |
|---|---|---|---|---|---|---|---|---|---|---|---|---|---|---|---|---|---|---|---|---|
| **SE** | 0.69 | 0.81 | 0.79 | 0.81 | 1.38 | 0.81 | 0.83 | 0.72 | 0.67 | 0.80 | 0.99 | 0.73 | 0.75 | 1.17 | 1.71 | 0.81 | 0.68 | 1.75 | 1.10 | 1.08 |
| **G** | | | | | | 6 | | | | | | 74 | | | | | | | | |
| **A** | | | 71 | | | | | | | | | | | 59 | 17 | | | 12 | | 62 |
| **P** | | | | | 34 | | | | | | | | | | | | | | 7 | |
| **V** | | | | | | | 63 | | | | | | 78 | | | | | | | 11 |
| **L** | | | | | 19 | | | 67 | | | | | | | | 72 | | | | |
| **I** | | | | | | | 33 | | | | | | | | | | | | | 22 |
| **M** | | | | | | | | 32 | | | | | 13 | | 7 | 7 | | | | |
| **F** | | | | 34 | 36 | | | | | | | | | | | 19 | | | | |
| **Y** | | | | 62 | | | | | | | | | | | | | | | | |
| **W** | | | | | | | | | | | | | | | | | | | | |
| **H** | | | | | | | | | | | | | | | | | | | | |
| **S** | | | | | 7 | 78 | | | | 60 | | | | 25 | | | 43 | | 66 | |
| **T** | | | 24 | | | | | | | 36 | | | | 5 | 31 | | 57 | 23 | | |
| **C** | | | | | | | | | | | | | | | | | | | 19 | |
| **N** | | | | | | 10 | | | | | | | | | | | | | | |
| **Q** | | | | | | | | | | | | | | | | | | 8 | | |
| **R** | 62 | | | | | | | | 26 | | 39 | 21 | | | 10 | | | | | |
| **K** | 38 | | | | | | | | 71 | | 53 | | | | 28 | | | 34 | | |
| **E** | | 65 | | | | | | | | | 5 | | | 5 | | | | 16 | | |
| **D** | | 31 | | | | | | | | | | | | | | | | | | |

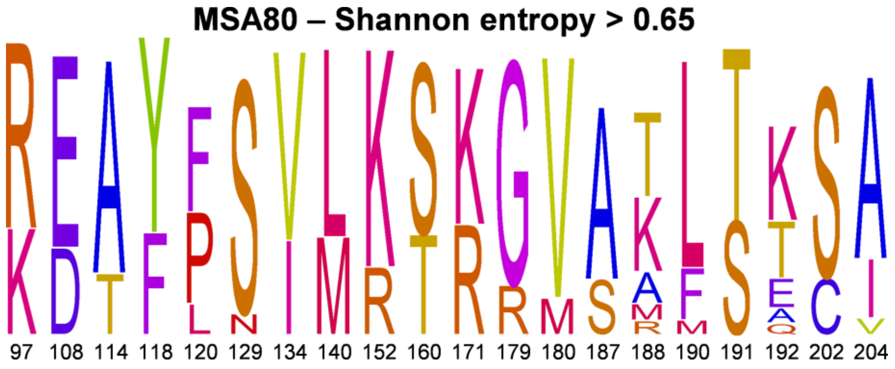

### 2. Phylogenetic analysis

First, a protein sequence search was performed using wild-type frataxin (residues 91-210) as a query and the Basic Local Alignment Search Tool (BLAST; https://blast.ncbi.nlm.nih.gov) on the non-redundant protein sequences (nr) database, excluding models (XM,XP), non-redundant RefSeq proteins (WP) and uncultured/environmental sample sequences with default parameters (*blastp* algorithm expect threshold 0.05, word size 5, max matches in a query range 0, matrix scoring BLOSUM62, gap costs of existence: 11 and extension 1, conditional compositional score matrix adjustment and no filters or masks). 994 sequences were found, which were filtered to ≥ 80% identity resulting in 389 sequences. A distance tree representation of these results (Blast Tree View; tree method: fast minimum evolution; max seq difference: 0.9; distance: Grishin protein; sequence label: Blast name) clearly shows evolutionary differences between amphibians/reptilians/birds and mammals characterized by mutations at positions 120 and 160.

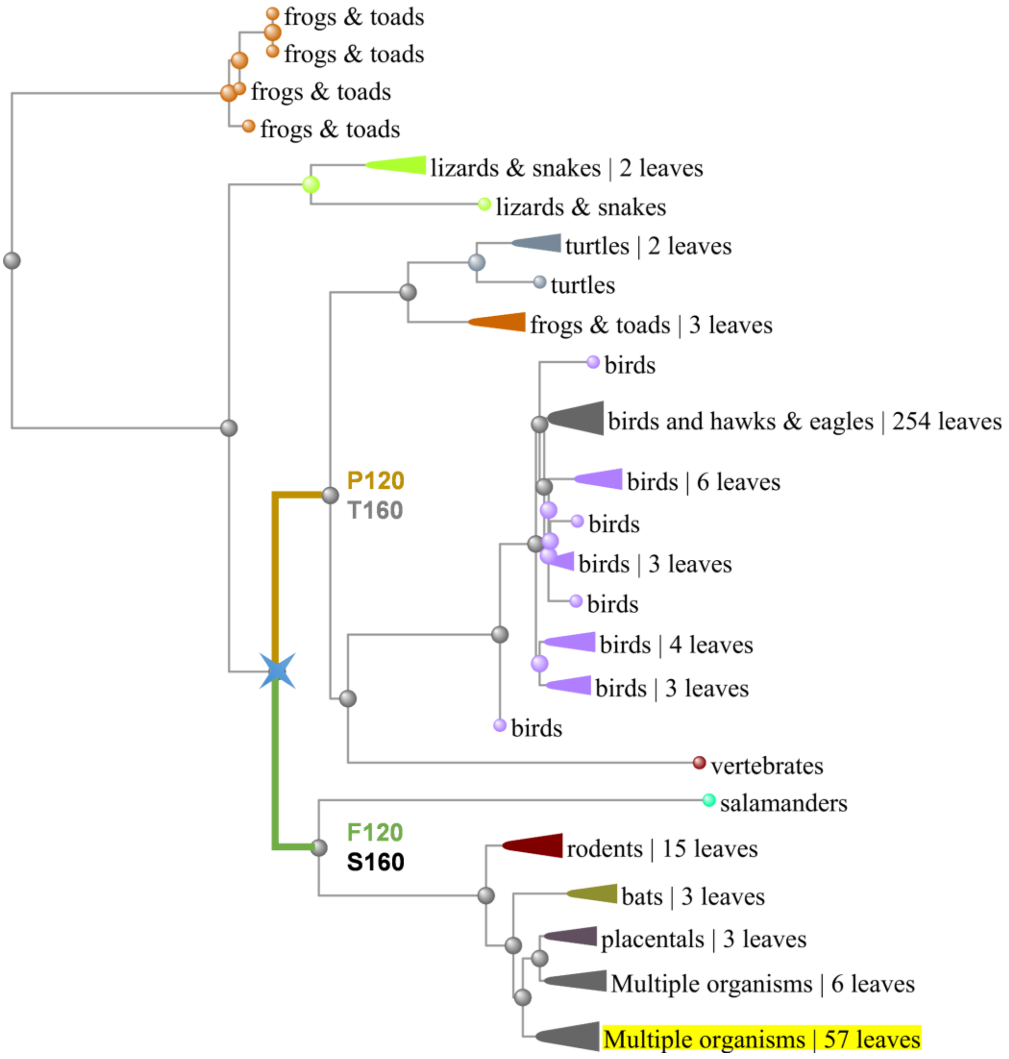


**Figure S2.** Blast tree view of sequences producing significant alignments to wild-type frataxin (contained in the cluster highlighted in yellow). The node characterized with mutations at positions 120 and 160 is highlighted with a blue star.

Second, the MSA80 dataset (180 sequences) was curated removing redundancies and sequences of unknown origin to yield 141 sequences, and their taxonomy annotated (kingdom, phylum, class, order, family, genus, species) using the Global Biodiversity Information Facility (GBIF; https://www.gbif.org/tools/species-lookup). Sequences were clustered at positions 120, 160 and 191 and an approximately-maximum-likelihood phylogenetic tree was constructed using FastTree,[12,13] and represented using Interactive Tree Of Life (iTOL; https://itol.embl.de). A clear partitioning is observed between different orders of mammals and amphibians/reptilians/birds (characterized by the F120P and S160T mutations with respect to human frataxin).

**Table S7.** Number of sequences and population of the clusters found within the curated MSA80 dataset (141 sequences) as a function of the amino acid identities at positions 120, 160 and 191, together with the taxonomic classes and orders of the species characterizing these clusters.

| Position | | | Num. of. sequences | Population | Taxonomic class | Taxonomic order |
|---|---|---|---|---|---|---|
| 120 | 160 | 191 | | | | |
| Phe | Ser | Thr | 49 | 34.8% | Mammalia | Primates, carnivora, chiroptera, cetacea |
| Phe | Ser | Ser | 6 | 4.3% | Mammalia | Afrosoricida, macroscelidea, proboscidea |
| Leu | Ser | Thr | 27 | 19.1% | Mammalia | Rodentia, carnivora |
| Leu | Cys | Thr | 2 | 1.4% | Mammalia | Rodentia |
| Ser | Ser | Ser | 4 | 2.8% | Mammalia | Monotremata, diprotodontia |
| Ser | Ser | Thr | 2 | 1.4% | Mammalia | Rodentia |
| Ser | Thr | Ser | 3 | 2.1% | Mammalia | Didelphimorphia, diprotodontia |
| Ser | Thr | Thr | 1 | 0.7% | Mammalia | Soricomorpha |
| Cys | Ser | Thr | 5 | 3.5% | Mammalia | Chiroptera, cetacea |
| Arg | Ser | Thr | 1 | 0.7% | Mammalia | Erinaceomorpha |
| Pro | Thr | Ser | 40 | 28.4% | Aves<br>Reptilia<br>Amphibia | Passeriformes, galliformes, crocodylia, testudines |
| Pro | Ile | Ser | 1 | 0.7% | Reptilia | Squamata |

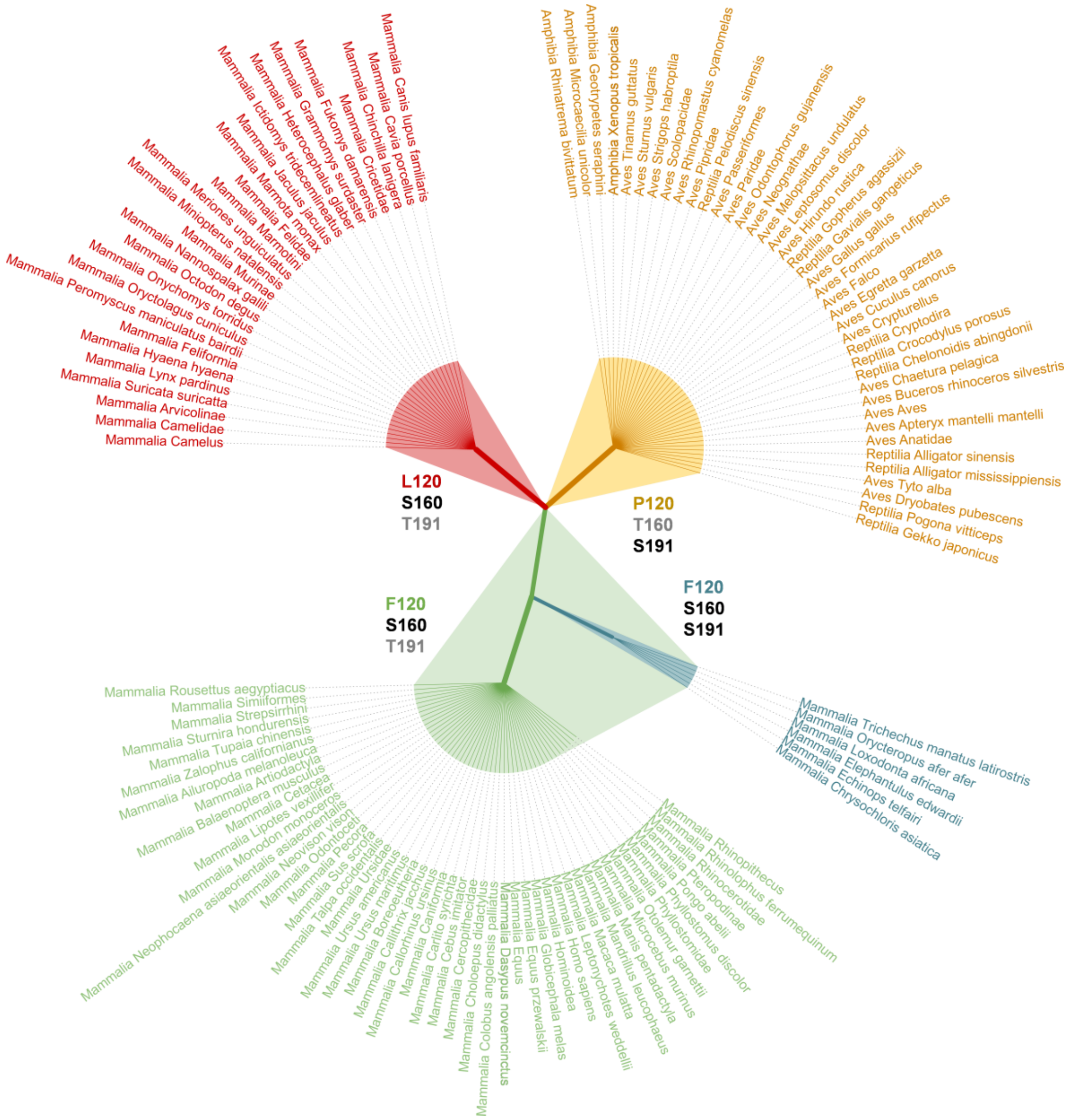


**Figure S3.** Unrooted phylogenetic tree of the sequences in a curated version of the MSA80 dataset, focusing on positions 120, 160 and 191. Branch lengths are ignored for clarity. Only branches with population >4% (highlighted with the corresponding colors in Table S7) are shown for clarity.

### 3. Pathological mutations

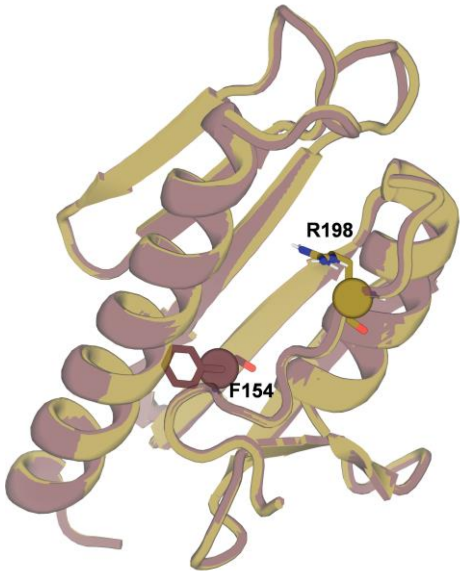


**Figure S4.** Overlay of AlphaFold models for FTX-I154F and FTX-L198R frataxin variants showing the location of the pathological mutations.

### 4. Sequence sampling with ProteinMPNN

Reference sequences were mutated simultaneously at all the selected hotspots for each mutation pool with ProteinMPNN[14] at three so-called temperatures (0.1, 0.2, and 0.3) excluding mutations to cysteine. In ProteinMPNN, the temperature is an hyperparameter that influences the amino acid compositional bias; lower temperatures have been shown to increase the frequency of charged over polar amino acids in the designs, leading to increased thermostability.[14] For each set of mutational hotspots, 30 sequences were generated (10 sequences per temperature) and filtered selecting either the most frequently occurring ones or, more often, the sequences corresponding to the highest thermostabilization over the reference frataxin variant predicted with our AlphaFold/Rosetta-based protocol (see below).

### 5. Prediction of relative thermostability of designed variants

For each design, we calculated its relative thermostability ($\Delta\Delta G^{calc}$) defined as the difference in free energy of *unfolding* between each variant and the reference (either wild-type frataxin or the pathological single mutants I154F or L198R) using the approach described recently by our group.[15] Briefly, each sequence is subjected to 30 independent AlphaFold structure predictions (models 3-5, which do not use PDB templates, see Table S9 for a characterization of structure similarity) and each of the 90 generated models is scored using Rosetta's *minimize* application.[16]

The average Rosetta energies of the 25 top scoring decoys of each protein represents its *folding* free energy ($\Delta G_f$), in such a way that relative stability values are computed as $\Delta\Delta G^{calc} = -(\Delta G^{calc}_{design,f} - \Delta G^{calc}_{reference,f})$. Therefore, positive and negative $\Delta\Delta G^{calc}$ values indicate stabilization and destabilization, respectively.

For consistency, predictions on all variants were made using sequences excluding the N-terminal residues derived from the inserted affinity tags (i.e., starting at position 91) and the two last very flexible, negatively charged (Asp-Glu) C-terminal residues (i.e., finishing at position 208).

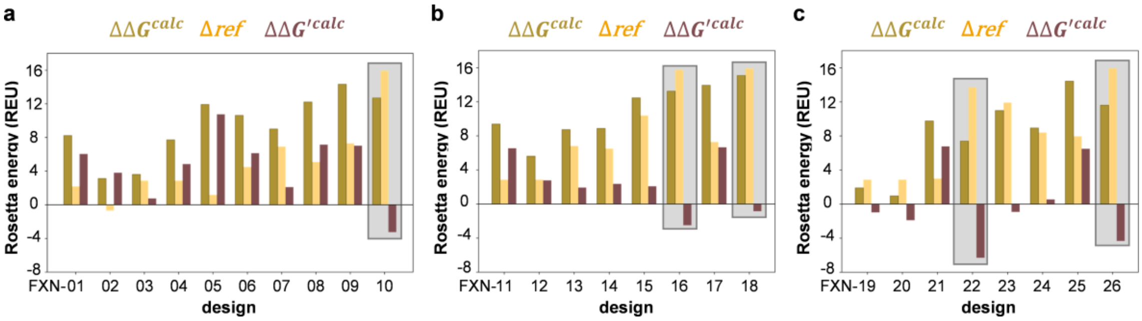


**Figure S5.** Calculated energies decomposition ($\Delta\Delta G'^{calc} = \Delta\Delta G^{calc} - \Delta ref$) for variants designed to stabilize a) wild-type frataxin, b) pathological mutant FXN-I154F, and c) pathological mutant FXN-L198R. In each case the corresponding frataxin variant is used as a reference. Variants whose calculated stabilization is clearly dominated by the $\Delta ref$ term are highlighted in grey.

The most computationally demanding part of our ProteinMPNN/AlphaFold/Rosetta protein design protocol is the generation of the AlphaFold ensembles (models 3, 4, 5; 30 replicas; 90 predicted structures in total). Overall, designing and characterizing a mutant takes under an hour (around 40 minutes if precomputed MSA are reused), making this protocol attractive for protein engineering in the scale of hundreds of variants.

**Table S8.** Timings for computational $\Delta\Delta G^{calc}$ prediction of a frataxin variant. Hardware: Intel Xeon Gold 6240R CPU 2.40GHz, Nvidia Geforce RTX 3090 GPU.

| Type of calculation | Software | Hardware | Time |
|---|---|---|---|
| ProteinMPNN sampling | ProteinMPNN 1.0.1 | CPU (1 core, 4 GB RAM memory)<br>GPU (24 GB) | 13 s |
| MSA generation | AlphaFold 2.3.0 | CPU | 20 m |
| Structure prediction | AlphaFold 2.3.0<br>models 3, 4, 5<br>30 replicas | CPU (1 core, 80 GB RAM memory)<br>GPU (24 GB) | 36 m |
| Minimization | Rosetta 3.13 | CPU (1 core, 2 GB RAM memory) | 1 m |

**Table S9.** Minimum average root-mean-square deviation of the atomic positions of the alpha carbons (RMSD-Cα), solvent-accessible surface area (SASA; probe radius 1.4 Å) and average Rosetta stability ($\Delta G_f^{calc}$) in Rosetta energy units (REU) within the AlphaFold ensemble (top 25 decoys) for wild-type, pathological mutants and designed frataxin variants. Relative stability values ($\Delta\Delta G^{calc}$) are computed with respect to wild-type frataxin and pathological variants FXN-I154F or FXN-L198R.

| **Variant** | **RMSD-Cα (Å)** | **SASA (Å²)** | $\Delta G_f^{calc}$ **(REU)** | $\Delta\Delta G^{calc}$ **(REU)**[a] | $\Delta\Delta G^{calc}$ **(REU)**[b] | $\Delta\Delta G^{calc}$ **(REU)**[c] |
|---|---|---|---|---|---|---|
| wild-type FXN | 0.10 | 6442 ± 29 | –373.1 ± 1.1 | 0.0 | | |
| FXN-01 | 0.12 | 6480 ± 36 | –381.3 ± 2.0 | 8.2 | | |
| FXN-02 | 0.14 | 6482 ± 28 | –376.2 ± 1.2 | 3.1 | | |
| FXN-03 | 0.09 | 6454 ± 36 | –377.2 ± 1.5 | 4.1 | | |
| FXN-04 | 0.09 | 6453 ± 34 | –380.8 ± 1.9 | 7.7 | | |
| FXN-05 | 0.08 | 6456 ± 33 | –385.0 ± 1.9 | 11.9 | | |
| FXN-06 | 0.08 | 6497 ± 32 | –383.8 ± 1.7 | 10.6 | | |
| FXN-07 | 0.12 | 6537 ± 28 | –382.1 ± 1.2 | 9.0 | | |
| FXN-08 | 0.08 | 6608 ± 23 | –385.3 ± 1.5 | 12.2 | | |
| FXN-09 | 0.10 | 6213 ± 22 | –387.4 ± 1.4 | 14.3 | | |
| FXN-10 | 0.11 | 6606 ± 23 | –385.8 ± 1.5 | 12.7 | | |
| FXN-I154F | 0.12 | 6495 ± 16 | –367.0 ± 1.1 | –6.2 | 0.0 | |
| FXN-11 | 0.11 | 6514 ± 25 | –376.4 ± 1.4 | 3.3 | 9.5 | |
| FXN-12 | 0.12 | 6506 ± 16 | –372.7 ± 1.8 | –0.5 | 5.7 | |
| FXN-13 | 0.13 | 6547 ± 19 | –375.8 ± 2.6 | 2.7 | 8.8 | |
| FXN–14 | 0.15 | 6460 ± 21 | –375.9 ± 1.3 | 2.8 | 9.0 | |
| FXN–15 | 0.08 | 6706 ± 21 | –379.5 ± 1.2 | 6.4 | 12.5 | |
| FXN–16 | 0.13 | 6652 ± 19 | –380.3 ± 1.3 | 7.2 | 13.3 | |
| FXN–17 | 0.16 | 6253 ± 19 | –381.0 ± 0.7 | 7.9 | 14.0 | |
| FXN–18 | 0.09 | 6648 ± 15 | –382.1 ± 1.2 | 9.0 | 15.2 | |
| FXN-L198R | 0.11 | 6438 ± 34 | –365.4 ± 1.5 | –7.7 | | 0.0 |
| FXN–19 | 0.15 | 6421 ± 38 | –368.0 ± 1.6 | –5.1 | | 2.6 |
| FXN–20 | 0.15 | 6415 ± 37 | –367.1 ± 2.1 | –6.1 | | 1.7 |
| FXN–21 | 0.12 | 6277 ± 16 | –375.9 ± 2.3 | 2.7 | | 10.5 |
| FXN–22 | 0.17 | 6601 ± 26 | –373.5 ± 3.0 | 0.4 | | 8.1 |
| FXN–23 | 0.13 | 6496 ± 45 | –377.1 ± 1.4 | 4.0 | | 11.7 |
| FXN–24 | 0.13 | 6623 ± 37 | –375.0 ± 1.7 | 1.9 | | 9.6 |
| FXN–25 | 0.11 | 6236 ± 41 | –380.5 ± 2.1 | 7.4 | | 15.1 |
| FXN–26 | 0.11 | 6561 ± 33 | –377.7 ± 1.5 | 4.6 | | 12.3 |

[a] Relative stability calculated with respect to wild-type FXN. [b] Relative stability calculated with respect to FXN-I154F. [c] Relative stability calculated with respect to FXN-L198R.

### 6. Protein expression and purification

The recombinant plasmid pG-S21a (purchased from GenScript Biotech) encoding between restriction sites *NdeI* and *XhoI* for residues 91-210 of either the wild-type frataxin, the pathologic single mutants I154F and L198R or any of the 26 designed proteins bearing a N-terminal 6xHis tag, was transformed into BL21 (D3) *E. coli* competent cells, plated on Luria-Bertani (LB) broth-ampicillin agar plates and incubated overnight at 37 ºC. A single colony from each plate was picked and then resuspended in an aqueous solution of 10 mL of LB broth (Lennox) and 10 μL of 50 mg/mL ampicillin followed by incubation at 37 ºC until the optical density at 600 nm (OD600) reached a value between 0.4-0.6. Induction was carried out by adding 10 μL of 0.5 M isopropyl β-D-1-thiogalactopyranoside (IPTG) and growth continued overnight at 25 ºC. Initial designs FXN-01 and FXN-02 (and a non-6xHis-tagged version of wild-type frataxin) were expressed following the same protocol, but the encoded genes were equipped with a N-terminal 6xHis-GST tag. Cells were harvested by centrifugation at 5000 rpm for 20 min. The cell pellets were resuspended in 40 mL of lysis buffer (120 mM NaCl, 20 mM Tris pH 8.0, 2 mM imidazole, 1 mM protease inhibition cocktail PIC). The suspended cells were lysed by sonication (60% amplitude, 36 x 10 s bursts, with 20 s between each burst), and then clarified by centrifugation at 25000 rpm for 30 minutes at 4 ºC. The soluble fraction was loaded onto 2 mL of Ni-NTA resin (Merck), cleaned with 10 mL of washing buffer (120 mM NaCl, 20 mM Tris pH 8.0) and eluted with 3.5 mL of high-imidazole buffer (120 mM NaCl, 20 mM Tris pH 8.0, 300 mM imidazole). The 6xHis-GST tag of designs FXN-01 and FXN-02 was cleaved by incubation with 2 IU of thrombin per mg of target protein at 4 ºC overnight. The cleaved protein was separated from 6xHis-GST and the uncleaved protein by a second affinity purification step using the Ni-NTA resin. For subsequent imidazole removal the buffer of the pooled fractions was exchanged to 20 mM Tris pH 8.0 and 120 mM NaCl using PD-10 desalting columns packed with Sephadex G-25 resin (GE Healthcare). The elution volume was 3.5 mL. Protein purity (size: ~13.8 to 14.4  kDa) was monitored using SDS-PAGE (5–20% gradient gel) and the concentrations were determined spectrophotometrically using an extinction coefficient ε in the 25440–26930 $cm^{-1}$ $M^{-1}$ range as determined by the ProtParam tool (https://web.expasy.org/protparam).

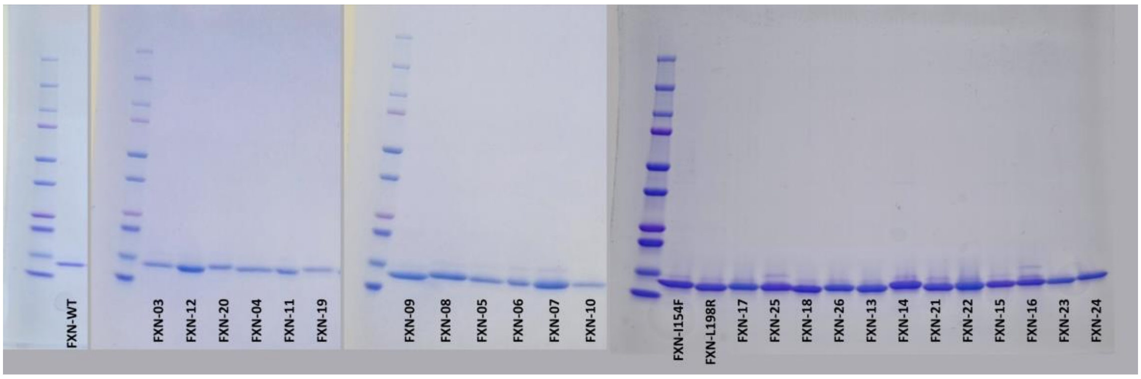


**Figure S6.** SDS-PAGE gels of expressed frataxin variants.

**7. Amino acid sequences of expressed frataxin variants.** Mutations with respect to wild-type human frataxin are highlighted in cyan. Pathological mutations are highlighted in magenta. N-terminal residues highlighted in grey correspond to 6xHis or 6xHis-GST tags (after thrombin cleavage).

>**wild-type frataxin** (non-6xHis-tagged)
GSHMDETTYERLAEETLDSLAEFFEDLADKPYTFEDYDVSFGSGVLTVKLGGDLGTYVINKQTPNKQIWL
SSPSSGPKRYDWTGKNWVYSHDGVSLHELLAAELTKALKTKLDLSSLAYSGKDA

>**wild-type frataxin** (6xHis-tagged)
HHHHHHGDETTYERLAEETLDSLAEFFEDLADKPYTFEDYDVSFGSGVLTVKLGGDLGTYVINKQTPNKQ
IWLSSPSSGPKRYDWTGKNWVYSHDGVSLHELLAAELTKALKTKLDLSSLAYSGKDA

>**FXN-01**
GSHMDETTYERLAEETLDSLAEFFEDLADKPYTPEDYDVSFGSGVLTVKLGGDLGTYVINKQTPNKQIWL
SSPTSGPKRYDWTGRNWVYSHDGVSLHELLSTELSTALKTKLDLSSLAYSGKDA

>**FXN-02**
GSHMDETTYERLAEETLDSLAEFFEDLADKPYTFEDYDVSFGSGVLTVKLGGDLGTYVINKQTPNKQIWL
SSPTSGPKRYDWTGRNWVYSHDGVSLHELLSTELSTALKTKLDLSSLAYSGKDA

>**FXN-03**
HHHHHHGDETTYERLAEETLDSLAEFFEDLADKPYTPEDYDVSFGSGVLTVKLGGDLGTYVINKQTPNKQ
IWLSSPSSGPKRYDWTGKNWVYSHDGVSLHELLAAELTKALKTKLDLSSLAYSGKDA

>**FXN-04**
HHHHHHGDETTYERLAEETLDSLAEFFEDLADKPYTPEDYDVSFGSGVLTVKLGGDLGTYVINKQTPNKQ
IWLSSPTSGPKRYDWTGKNWVYSHDGVSLHELLAAELSKALKTKLDLSSLAYSGKDA

>**FXN-05**
HHHHHHGDETTYERLAEETLDSLAEFFEDLADKPYTPEDYDVSFGSGVLTVKLGGDLGTYVINKQTPNKQ
IWLSSPTSGPKRYDWTGTNWVYSHDGVSLHELLATELSKALKTKLDLSSLAYSGKDA

>**FXN-06**
HHHHHHGDETTYERLAEETLDSLAEFFEDLADKPYTPEDYDVSFGSGVLTVKLGGDLGTYVINKQTPNKQ
IWLSSPTSGPKRYDWTGSNWVYSHDGVSLHELLAKELSKALKTKLDLSSLAYSGKDA

>**FXN-07**
HHHHHHGDETTYERLAEETLDSLAEFFEDLADKPYTPEDYDVSFGSGVLTVKLGGDLGTYVINKQTPNKQ
IWLSSPTSGPKRYDWTGENWVYSHDGVSLHELLAKELSKALKTKLDLSSLAYSGKDA

>**FXN-08**
HHHHHHGDETTYEKLAEETLDSLAEFFEDLADKPFTPEDYDVSFGSGVLTVKLGGDLGTYVINKQTPNKQ
IWLSSPTSGPKRYDWTGTNWVYSHDGVSLHELLAKELSKALKTKLDLSHLKYSGKDA

>**FXN-09**
HHHHHHGDESTYERLAEETLDSLADFFEDLKDQPFTQPDYDVSFGSGVLTVKLGGDLGTYVINKQTPNKQ
IWLSSPTSGPKRYDWTGSSWVYSHDGVSLHELLSEELTKLLKTPIDLSHLAYSGKDA

>**FXN-10**
HHHHHHGDETTYEKLAEETLDSLAEFFEDLKDKPFTPEDYDVSFGDGVLTVKLGGDLGTYVINKQTPNKQ
IWLSSPTSGPKRYDWTGTNWVYSHDGKSLHELLSEELSKALKTKLDLSHLKYSGKDA

>**FXN-I154F**
HHHHHHGDETTYERLAEETLDSLAEFFEDLADKPYTFEDYDVSFGSGVLTVKLGGDLGTYVINKQTPNKQ
FWLSSPSSGPKRYDWTGKNWVYSHDGVSLHELLAAELTKALKTKLDLSSLAYSGKDA

>**FXN-11**
HHHHHHGDETTYERLAEETLDSLAEFFEDLADKPYTPEDYDVSFGSGVLTVKLGGDLGTYVINKQTPNKQ
FWLSSPTSGPKRYDWTGKNWVYSHDGVSLHELLAAELSKALKTKLDLSSLAYSGKDA

**>FXN-12**
HHHHHHGDETTYERLAEETLDSLAEFFEDLADKPYTPEDYDVSFGSGVLTVKLGGDLGTYVINKQTPNKQ
FWLSSPSSGPKRYDWTGKNWVYSHDGVSLHELLAAELTKALKTKLDLSSLAYSGKDA

**>FXN-13**
HHHHHHGDEATYEKLAEETLDSLADFFEDLKDQPFTPEDYDVSFGSGVLTVKLGGDLGTYVINKQTPNKQ
FWLSSPTSGPKRYDWTGEAWVYSHDGVSLHELLAKELTKLLKTPIDLSHLKYSGKDA

**>FXN-14**
HHHHHHGDEATYERLAEETLDSLAEFFEDLKDQPFTPEDYDVSFGSGVLTVKLGGDLGTYVINKQTPNKQ
FWLSSPTSGPKRYDWTGESWVYSHDGVSLHELLAKELTTLLKTPIDLSHLKYSGKDA

**>FXN-15**
HHHHHHGDETTYEKLAEETLDSLAEFFEDLKDKPFTPEDYDVSFGDGVLTVKLGGDLGTYVINKQTPNKQ
FWLSSPTSGPKRYDWTGSNWVYSHDGVSLHELLAKELSKALKTKLDLSHLKYSGKDA

**>FXN-16**
HHHHHHGDETTYEKLAEETLDSLAEFFEDLKDKPFTPEDYDVSFGDGVLTVKLGGDLGTYVINKQTPNKQ
FWLSSPTSGPKRYDWTGSNWVYSHDGKSLHELLAEELSKALKTKLDLSSLKYSGKDA

**>FXN-17**
HHHHHHGDESTYERLAEETLDSLADFFEDLKDQPFTQPDYDVSFGSGVLTVKLGGDLGTYVINKQTPNKQ
FWLSSPTSGPKRYDWTGSSWVYSHDGVSLHELLSEELTKLLKTPIDLSHLAYSGKDA

**>FXN-18**
HHHHHHGDETTYEKLAEETLDSLAEFFEDLKDKPFTPEDYDVSFGDGVLTVKLGGDLGTYVINKQTPNKQ
FWLSSPTSGPKRYDWTGTNWVYSHDGKSLHELLSEELSKALKTKLDLSHLKYSGKDA

**>FXN-L198R**
HHHHHHGDETTYERLAEETLDSLAEFFEDLADKPYTFEDYDVSFGSGVLTVKLGGDLGTYVINKQTPNKQ
IWLSSPSSGPKRYDWTGKNWVYSHDGVSLHELLAAELTKALKTKRDLSSLAYSGKDA

**>FXN-19**
HHHHHHGDETTYERLAEETLDSLAEFFEDLADKPYTPEDYDVSFGSGVLTVKLGGDLGTYVINKQTPNKQ
IWLSSPTSGPKRYDWTGKNWVYSHDGVSLHELLAAELSKALKTKRDLSSLAYSGKDA

**>FXN-20**
HHHHHHGDETTYERLAEETLDSLAEFFEDLADKPYTPEDYDVSFGSGVLTVKLGGDLGTYVINKQTPNKQ
IWLSSPSSGPKRYDWTGKNWVYSHDGVSLHELLAAELTKALKTKRDLSSLAYSGKDA

**>FXN-21**
HHHHHHGDEATYERLAEETLDSLAAFFEDLKDQPETPEDYDVSFGSGVLTVKLGGDLGTYVINKQTPNKQ
IWLSSPTSGPKRYDWTGDTWVYSHDGVSLHELLATELTKLLKTPRDLSSLAYSGKDA

**>FXN-22**
HHHHHHGDEETYEKLAEETLDSLAEFFEDLKDQPFTPEDYDVSFGSGVLTVKLGGDLGTYVINKQTPNKQ
IWLSSPTSGPKRYDWTGESWVYSHDGVSLHELLAKELTKLLKTPRDLSHLKYSGKDA

**>FXN-23**
HHHHHHGDETTYEKLAEETLDSLATFFEDLKDKPFTPEDYDVSFGDGVLTVKLGGDLGTYVINKQTPNKQ
IWLSSPTSGPKRYDWTGSNWVYSHDGKSLHELLAEELSKALKTKRDLSHLKYSGKDA

**>FXN-24**
HHHHHHGDETTYEKLAEETLDSLAKFFEDLKDKPFTPEDYDVSFGDGVLTVKLGGDLGTYVINKQTPNKQ
IWLSSPTSGPKRYDWTGSNWVYSHDGVSLHELLAKELSKALKTKRDLSHLKYSGKDA

**>FXN-25**
HHHHHHGDESTYERLAEETLDSLADFFEDLKDQPFTQPDYDVSFGSGVLTVKLGGDLGTYVINKQTPNKQ
IWLSSPTSGPKRYDWTGSSWVYSHDGVSLHELLSEELTKLLKTPRDLSHLAYSGKDA

**>FXN-26**
HHHHHHGDETTYEKLAEETLDSLAEFFEDLKDKPFTPEDYDVSFGDGVLTVKLGGDLGTYVINKQTPNKQ
IWLSSPTSGPKRYDWTGTNWVYSHDGKSLHELLSEELSKALKTKRDLSHLKYSGKDA

**Table S10.** Number and identity of mutations, molecular weight (MW), theoretical extinction coefficient at λ = 280 nm in water ($E_{280}$), isoelectric point (IP) and total charge at pH 7.4 (*Z*) (calculated with Isoelectric Point Calculator 2.0[17]), and the differences in the number of charged amino acids (ΔN. charged) and prolines (ΔN. Pro) for wild-type, pathological mutants and designed frataxin variants. Pathological mutations are shown in red.

| Variant | N. mutations | Mutations | MW (Da) | $E_{280}$ ($M^{-1}$ $cm^{-1}$) | IP | *Z* | ΔN. charged | ΔN. Pro |
|---|---|---|---|---|---|---|---|---|
| wild-type FXN | 0 | | 14240.70 | 26930 | 5.22 | –8.1 | 0 | 0 |
| FXN-01 | 7 | F120P, S160T, K171R, A187S, A188T, T191S, K192T | 13770.17 | 26930 | 4.55 | –9.1 | -1 | 1 |
| FXN-02 | 6 | S160T, K171R, A187S, A188T, T191S, K192T | 13820.23 | 26930 | 4.55 | –9.1 | -1 | 0 |
| FXN-03 | 1 | F120P | 14190.64 | 26930 | 5.22 | –8.1 | 0 | 1 |
| FXN-04 | 3 | F120P, S160T, T191S | 14190.64 | 26930 | 5.22 | –8.1 | 0 | 1 |
| FXN-05 | 5 | F120P, S160T, K171T, A188T, T191S | 14193.59 | 26930 | 5.04 | –9.1 | -1 | 1 |
| FXN-06 | 5 | F120P, S160T, K171S, A188K, T191S | 14206.63 | 26930 | 5.22 | –8.1 | 0 | 1 |
| FXN-07 | 5 | F120P, S160T, K171E, A188K, T191S | 14248.67 | 26930 | 5.08 | –9.1 | 1 | 1 |
| FXN-08 | 9 | R97K, Y118F, F120P, S160T, K171T, A188K, T191S, S202H, A204K | 14283.81 | 25440 | 5.51 | –7.1 | 1 | 1 |
| FXN-09 | 16 | T93S, E108D, A114K, K116Q, Y118F, F120Q, E121P, S160T, K171S, N172S, A187S, A188E, A193L, K197P, L198I, S202H | 14283.68 | 25440 | 5.02 | –10.0 | -2 | 2 |
| FXN-10 | 13 | R97K, A114K, Y118F, F120P, S129D, S160T, K171T, V180K, A187S, A188E, T191S, S202H, A204K | 14414.90 | 25440 | 5.39 | –8.1 | 4 | 1 |
| FXN-I154F | 1 | I154F | 14274.71 | 26930 | 5.22 | –8.1 | | |
| FXN-11 | 3+1 | F120P, I154F, S160T, T191S | 14224.65 | 26930 | 5.22 | –8.1 | 0 | 1 |
| FXN-12 | 1+1 | F120P, I154F | 14224.65 | 26930 | 5.22 | –8.1 | 0 | 1 |
| FXN-13 | 16+1 | T93A, R97K, E108D, A114K, K116Q, Y118F, F120P, I154F, S160T, K171E, N172A, A188K, A193L, K197P, L198I, S202H, A204K | 14340.86 | 25440 | 5.22 | –9.1 | 1 | 2 |
| FXN-14 | 15+1 | T93A, A114K, K116Q, Y118F, F120P, I154F, S160T, K171E, N172S, A188K, K192T, A193L, K197P, L198I, S202H, A204K | 14371.83 | 25440 | 5.06 | –10.1 | 0 | 2 |
| FXN-15 | 11+1 | R97K, A114K, Y118F, F120P, S129D, I154F, S160T, K171S, A188K, T191S, S202H, A204K | 14388.90 | 25440 | 5.52 | –7.1 | 4 | 1 |
| FXN-16 | 11+1 | R97K, A114K, Y118F, F120P, S129D, I154F, S160T, K171S, V180K, A188E, T191S, A204K | 14368.82 | 25440 | 5.27 | –8.2 | 4 | 1 |
| FXN-17 | 16+1 | T93S, E108D, A114K, K116Q, Y118F, F120Q, E121P, I154F, S160T, K171S, N172S, A187S, A188E, A193L, K197P, L198I, S202H | 14317.69 | 25440 | 5.02 | –10.0 | -2 | 2 |
| FXN-18 | 13+1 | R97K, A114K, Y118F, F120P, S129D, I154F, S160T, K171T, V180K, A187S, A188E, T191S, S202H, A204K | 14448.91 | 25440 | 5.39 | –8.1 | 4 | 1 |
| FXN-L198R | 1 | L198R | 14283.72 | 26930 | 5.39 | –7.1 | | |
| FXN-19 | 3+1 | F120P, S160T, T191S, L198R | 14233.66 | 26930 | 5.39 | –7.1 | 1 | 1 |
| FXN-20 | 1+1 | F120P, L198R | 14233.66 | 26930 | 5.39 | –7.1 | 1 | 1 |
| FXN-21 | 12+1 | T93A, E108A, A114K, K116Q, Y118E, F120P, S160T, K171D, N172T, A188T, A193L, K197P, L198R | 14197.58 | 25440 | 4.93 | –10.0 | 0 | 2 |
| FXN-22 | 14+1 | T93E, R97K, A114K, K116Q, Y118F, F120P, S160T, K171E, N172S, A188K, A193L, K197P, L198R, S202H, A204K | 14437.93 | 25440 | 5.24 | –9.1 | 2 | 2 |
| FXN-23 | 13+1 | R97K, E108T, A114K, Y118F, F120P, S129D, S160T, K171S, V180K, A188E, T191S, L198R, S202H, A204K | 14399.89 | 25440 | 5.65 | –6.1 | 4 | 1 |
| FXN-24 | 12+1 | R97K, E108K, A114K, Y118F, F120P, S129D, S160T, K171S, A188K, T191S, L198R, S202H, A204K | 14396.97 | 25440 | 5.92 | –4.2 | 4 | 1 |
| FXN-25 | 15+1 | T93S, E108D, A114K, K116Q, Y118F, F120Q, E121P, S160T, K171S, N172S, A187S, A188E, A193L, K197P, L198R, S202H | 14326.71 | 25440 | 5.19 | –9.0 | -2 | 2 |
| FXN-26 | 13+1 | R97K, A114K, Y118F, F120P, S129D, S160T, K171T, V180K, A187S, A188E, T191S, L198R, S202H, A204K | 14457.92 | 25440 | 5.53 | –7.1 | 4 | 1 |

### 8. Circular dichroism (CD) spectroscopy

The secondary structure of all frataxin variants was analyzed using CD spectroscopy. Protein samples were prepared at 5 µM concentration in buffer solution (20 mM HEPES pH 7.4, 120 mM NaCl) in a 2 mm quartz cuvette. CD spectra were recorded on a JASCO J-815 spectropolarimeter equipped with a Peltier temperature control unit at 25 ºC. Data were collected from 200 to 300 nm with a 0.2 nm step size and a 2 nm bandwidth.

CD spectra were acquired first at 25 ºC using freshly prepared proteins (*run 1*, in blue in the plots below), and then after heating up to 90 ºC and cooling back to 25 ºC to test *unfolding*/*folding* reversibility (*run 2*, in orange in the plots below).

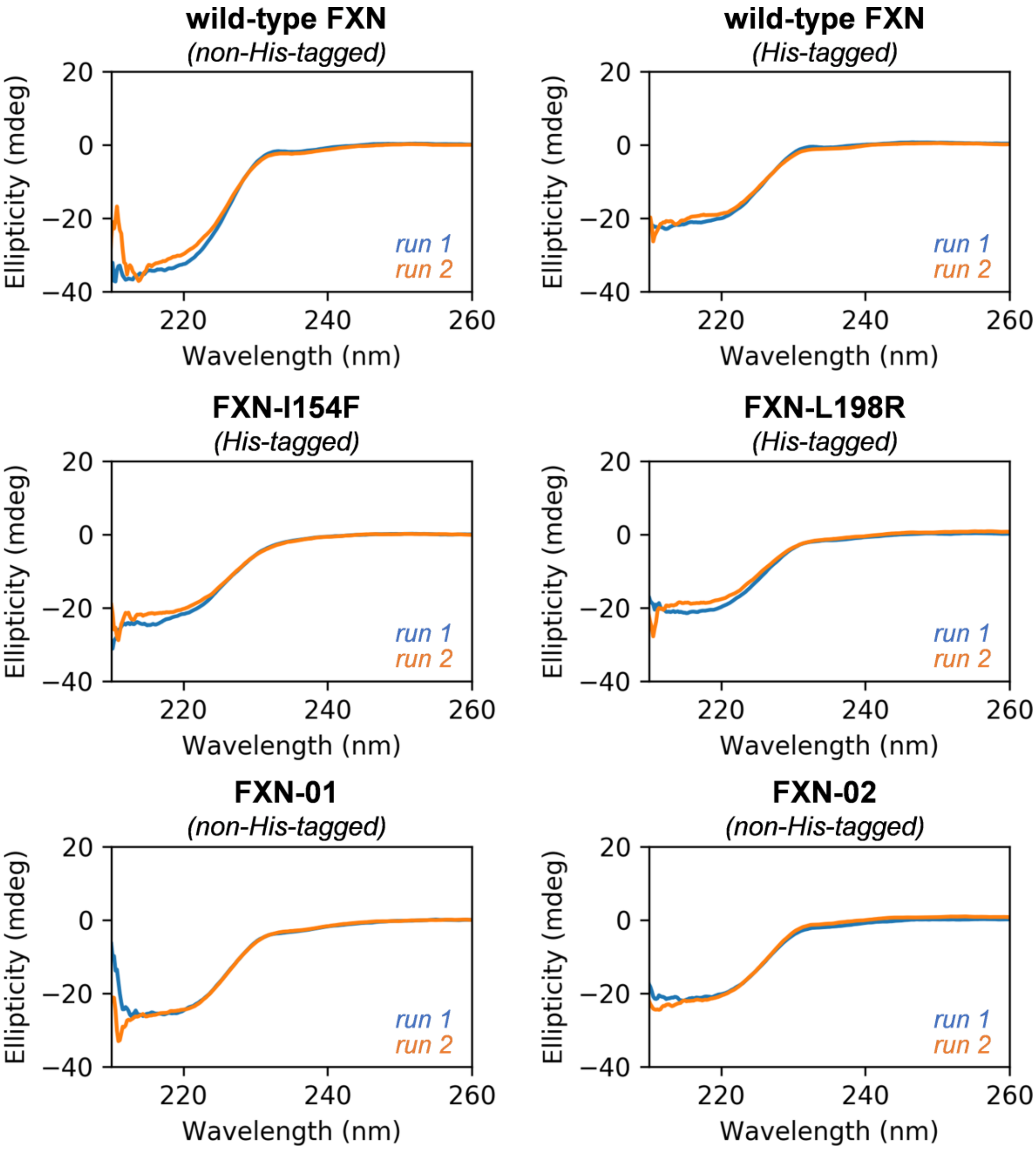

**Figure S7.** CD spectra of frataxin (FXN) variants (210-260 nm).

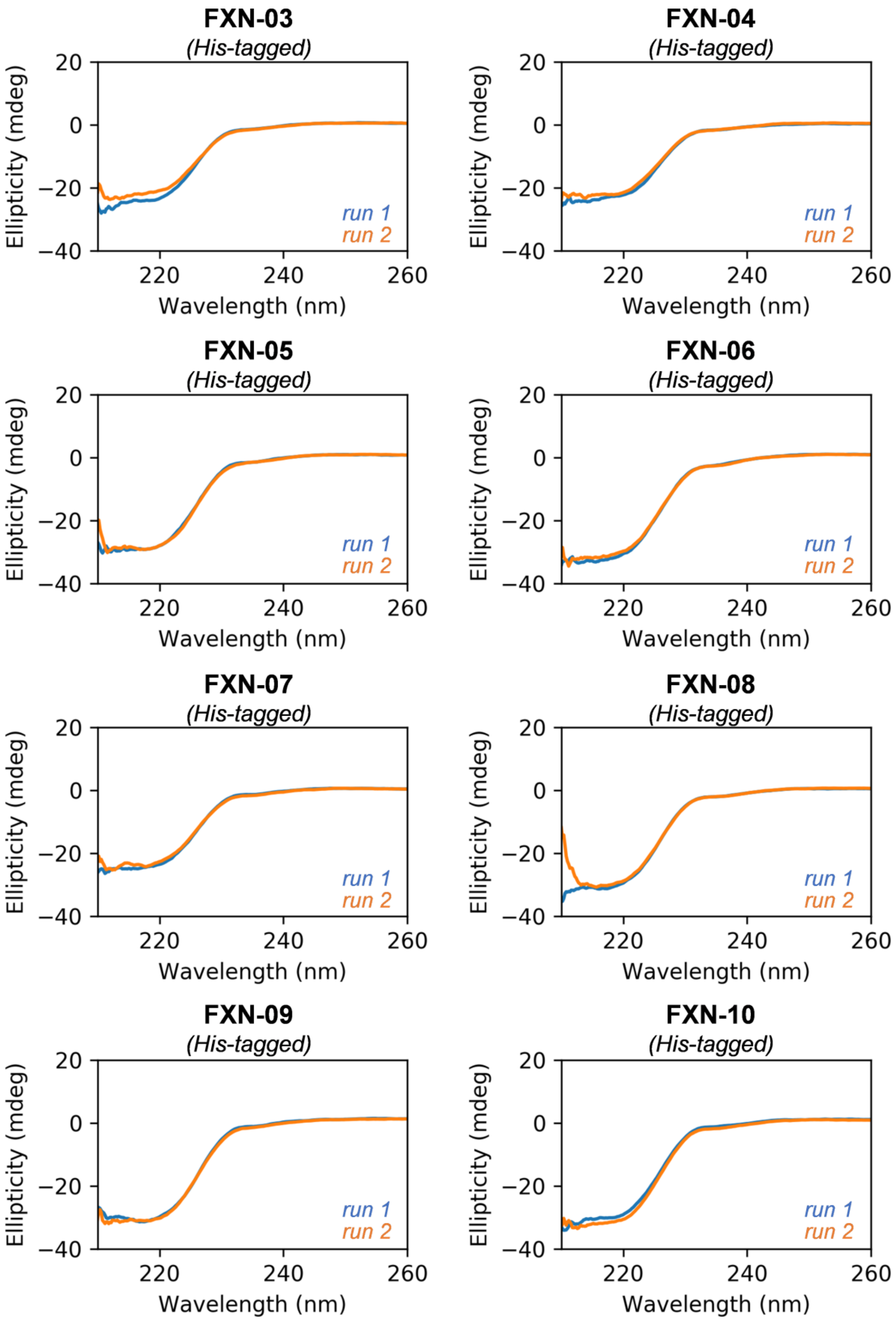


**Figure S7** (cont.). CD spectra of frataxin (FXN) variants (210-260 nm).

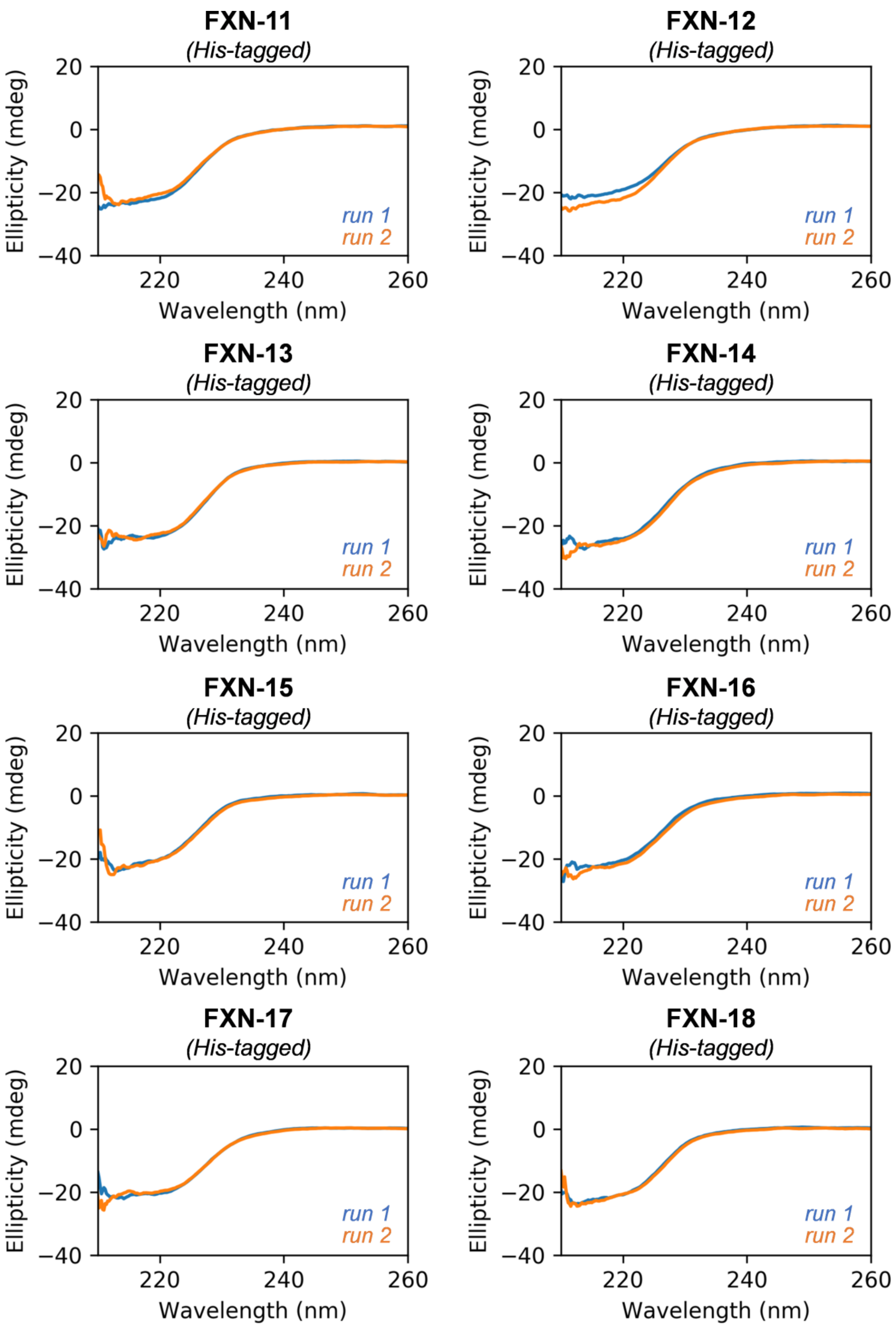


**Figure S7** (cont.). CD spectra of frataxin (FXN) variants (210-260 nm).

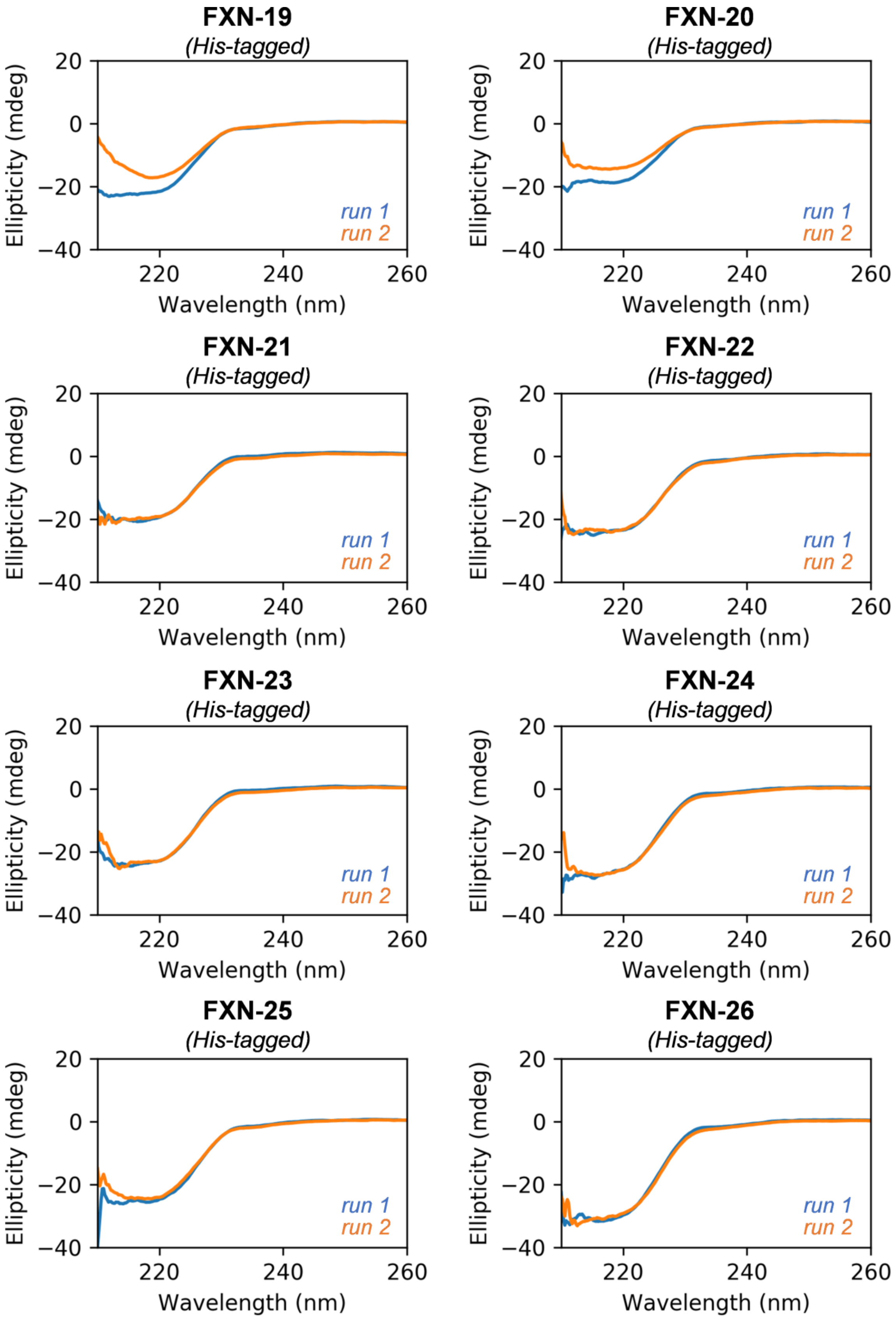


**Figure S7** (cont.). CD spectra of frataxin (FXN) variants (210-260 nm).

### 9. Melting temperature ($T_m$) measurement

Thermal stability was measured by monitoring the change in ellipticity at 222 nm as a function of temperature using a JASCO J-815 spectropolarimeter equipped with a Peltier temperature control unit. Protein samples were prepared at 5 µM concentration in buffer solution (20 mM HEPES pH 7.4, 120 mM NaCl) in a 2 mm quartz cuvette. The CD signal was monitored at a fixed wavelength (222 nm) while the temperature was increased from 35 to 90 ºC at a rate of 1.2 ºC/min. For the pathological FXN mutants I154F and L198R, temperature scans started at 15 ºC.

For each variant, $T_m$ values were computed by fitting ellipticity values ($E$) versus temperature to a two-state (*folded*/*unfolded*) model. The fitting was performed through a least-squares minimization of the error between the measured ellipticity values $E$ and simulated ellipticity values $E_{sim}$ obtained using the following equation:

$$E_{sim} = FF_{sim} \cdot (m_1 T + b_1) + (1 - FF_{sim}) \cdot (m_2 T + b_2) \qquad \text{eq. 2}$$

where $FF_{sim}$ is the fraction of protein in the native (folded) state, T is the temperature, and the parameters $m_1$, $b_1$, $m_2$, and $b_2$ (fitting parameters) are the slopes and intercepts of the (linear) ellipticity in the native and denatured state, respectively (Figure S8). Initial guesses of $m_1$, $b_1$, $m_2$, and $b_2$ were obtained from linear fitting of the first 20 ellipticity points ($m_1$, $b_1$) and the last 20 ellipticity points ($m_2$, $b_2$). $FF_{sim}$ is defined as a parametric function derived from the equilibrium constant of folding ($K_{sim}$).

$$FF_{sim} = \frac{K_{sim}}{1-K_{sim}} \qquad \text{eq. 3}$$

In turn, the equilibrium constant is obtained from the free energy of unfolding ($\Delta G_{sim}$):

$$K_{sim} = e^{\frac{\Delta G_{sim}}{RT}} \qquad \text{eq. 4}$$

where

$$\Delta G_{sim} = \Delta H_{sim} - T\Delta S_{sim} \qquad \text{eq. 5}$$

and the enthalpy ($\Delta H_{sim}$) and entropy ($\Delta S_{sim}$) of unfolding, are calculated from guesses of the enthalpy of unfolding at the melting temperature ($\Delta H_m$), the change in specific heat capacity of unfolding ($\Delta C_p$) and $T_m$, where $\Delta H_m$ and $T_m$ are the parameters to optimize and $\Delta C_p$ is a constant.

$$\Delta H_{sim} = \Delta H_m + \Delta C_p (T - T_m) \qquad \text{eq. 6}$$

$$\Delta S_{sim} = \Delta H_m + \Delta C_p ln\left(\frac{T}{T_m}\right) \qquad \text{eq. 7}$$

The initial guess values for $\Delta H_m$ and $T_m$ were set to 48 kcal mol$^{-1}$ and 333.15 K (60 °C), respectively. The value of $\Delta C_p$ is estimated from the number of residues of the protein.[18]

Assuming a linear relationship between the number of residues ($n$) and the change in solvent accessible surface area of unfolding ($\Delta SASA$), the latter can be computed as:

$$\Delta SASA\ (\text{Å}^2) = -907 + 93 \cdot n \qquad \text{eq. 8}$$

From $\Delta ASA$, $\Delta C_p$ can be estimated as:

$$\Delta C_p (cal\ mol^{-1}\ K^{-1}) = -251 + 0.19 \cdot \Delta SASA \qquad \text{eq. 9}$$

For the 127 residues frataxin and its variants, $\Delta SASA = 10904$ Å$^2$ and $\Delta C_p = 1.82$ kcal mol$^{-1}$ K$^{-1}$.

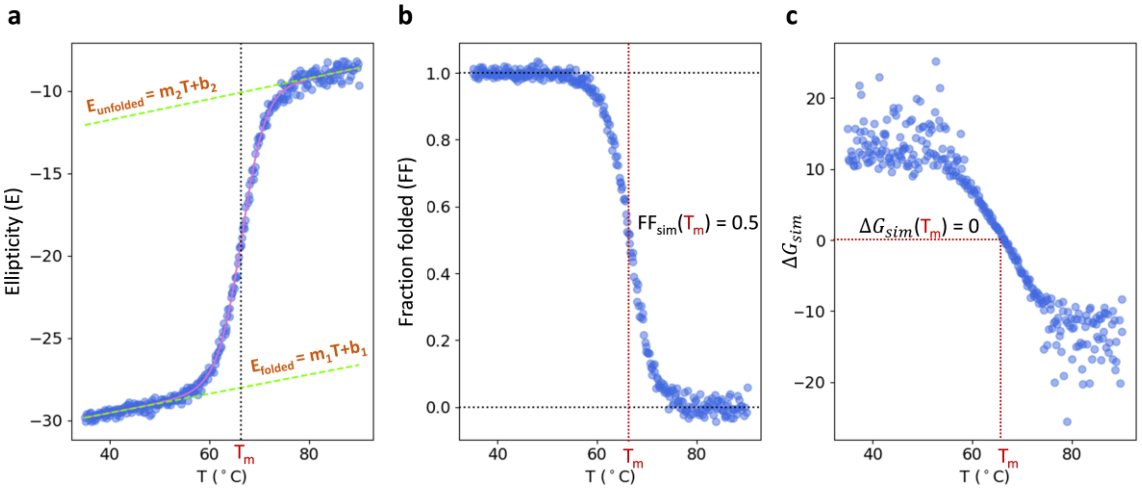


**Figure S8**. a) Evolution of the ellipticity as a function of temperature along thermal unfolding. Blue dots: measured E(T) values; magenta line: fitted $E_{sim}$(T). b) Evolution of the fraction of folded protein as a function of temperature along thermal unfolding. c) Evolution of the free energy of unfolding as a function of temperature along thermal unfolding.

In Figure S9, $T_M$ curves were represented considering for each variant the *heating* and *cooling* ellipticity values and linearly rescaling them to a [0,1] range according to:

$$E_{norm} = \frac{E - E_{min}}{E_{max} - E_{min}} \qquad \text{eq. 10}$$

where $E$ is the measured ellipticity, $E_{min}$ the absolute minimum of $E$ values considering the *heating* and *cooling* curves, and $E_{max}$ the absolute maximum under the same conditions.

Folding reversibility percentages were computed from the fitted ellipticity values (eq. 2) according to the following equation:

$$\text{reversibility (\%)} = \frac{E_{90}^{cool} - E_{35}^{cool}}{E_{90}^{heat} - E_{35}^{heat}} \times 100 \qquad \text{eq. 11}$$

where $E_{90}^{cool}$, $E_{35}^{cool}$, $E_{90}^{heat}$ and $E_{35}^{heat}$ are the ellipticities measured at 90 ºC and 35°C during *heating* (i.e., forward thermal unfolding) and *cooling* (reverse thermal folding).

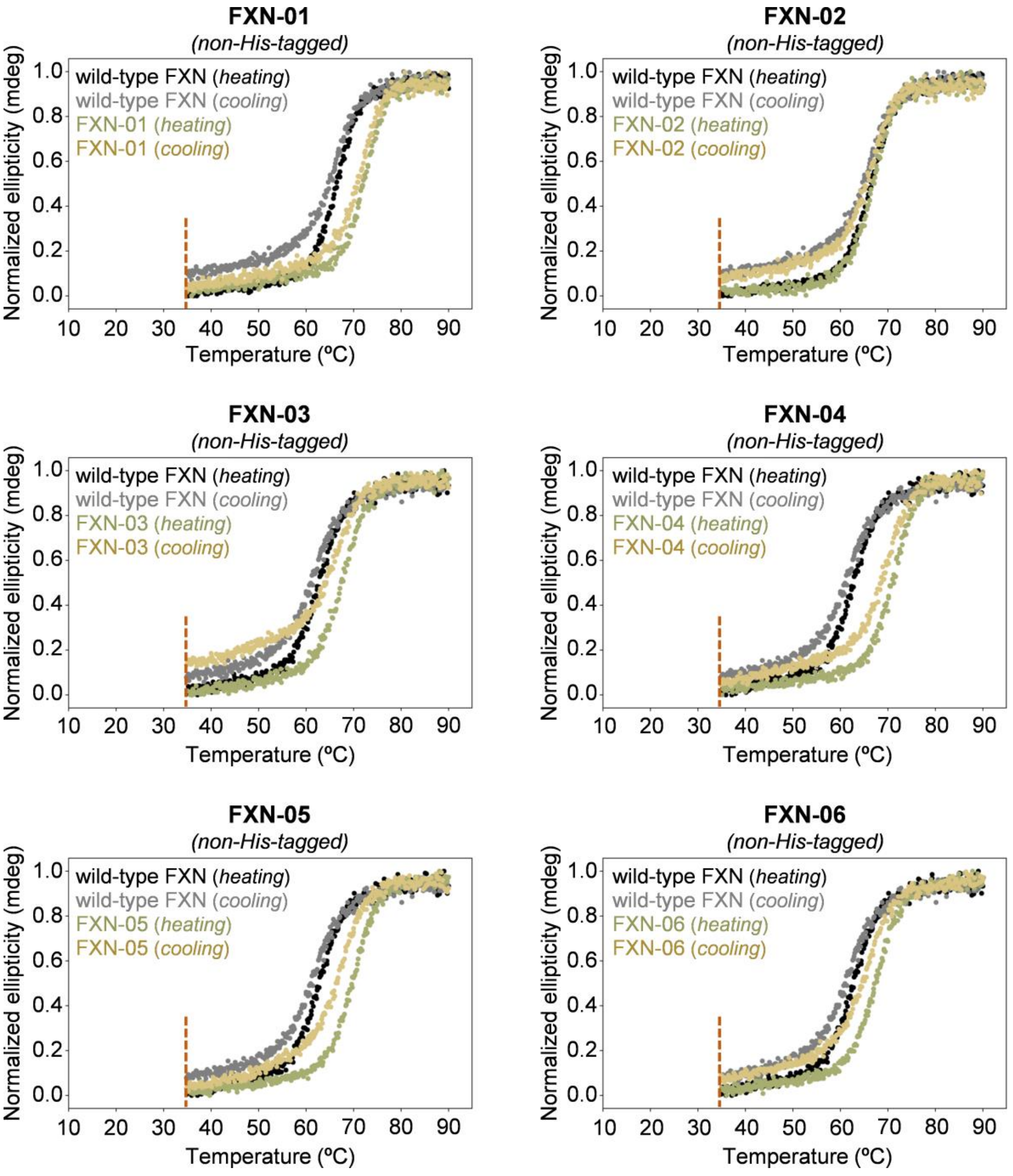


**Figure S9.** Thermal unfolding of frataxin (FXN) variants measured by circular dichroism (CD) spectroscopy. Reversibility is always measured at 35 ºC for fair comparison across different variants (dashed red line).

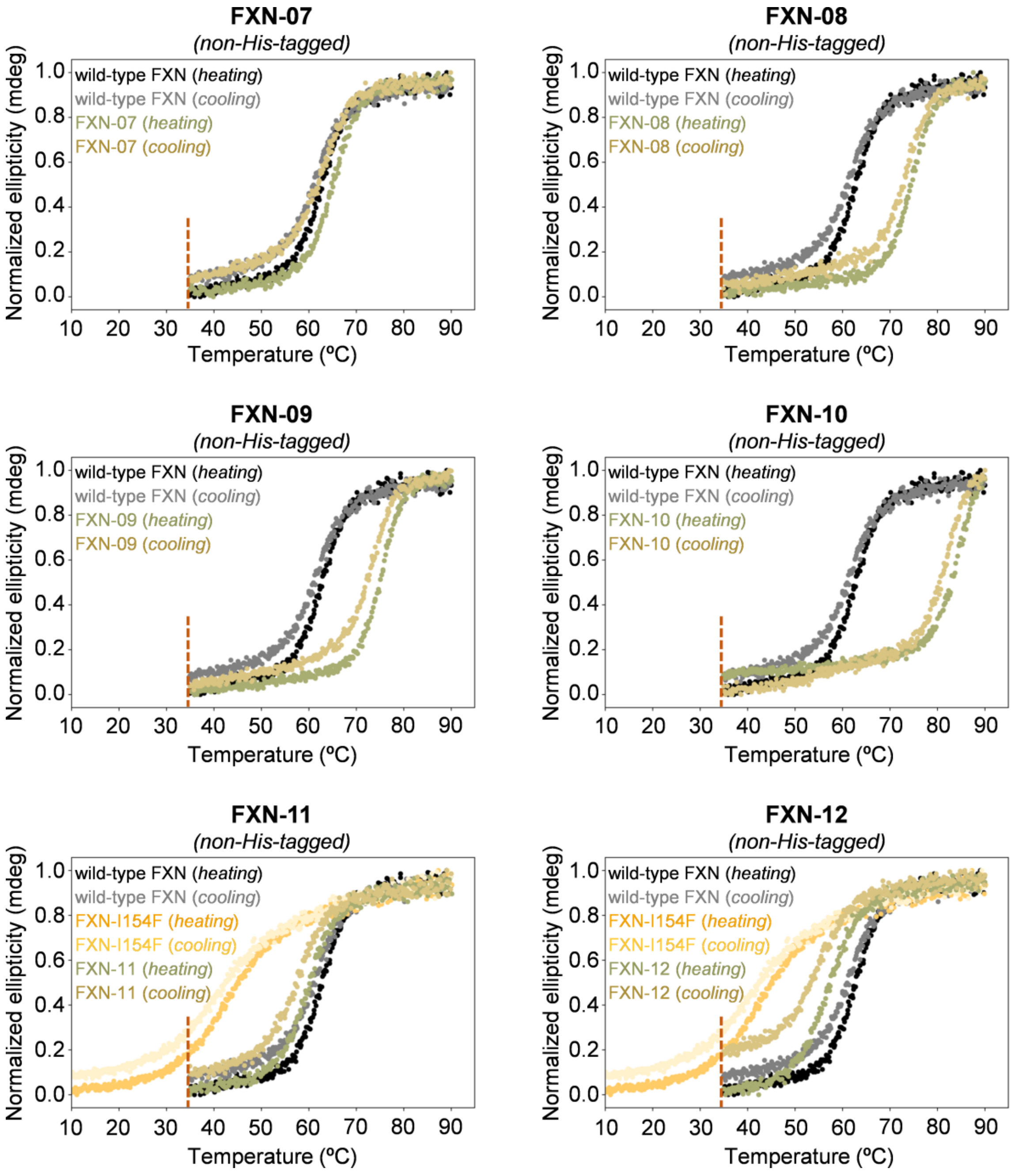


**Figure S9** (cont.). Thermal unfolding of frataxin (FXN) variants measured by circular dichroism (CD) spectroscopy. Reversibility is always measured at 35 ºC for fair comparison across different variants (dashed red line).

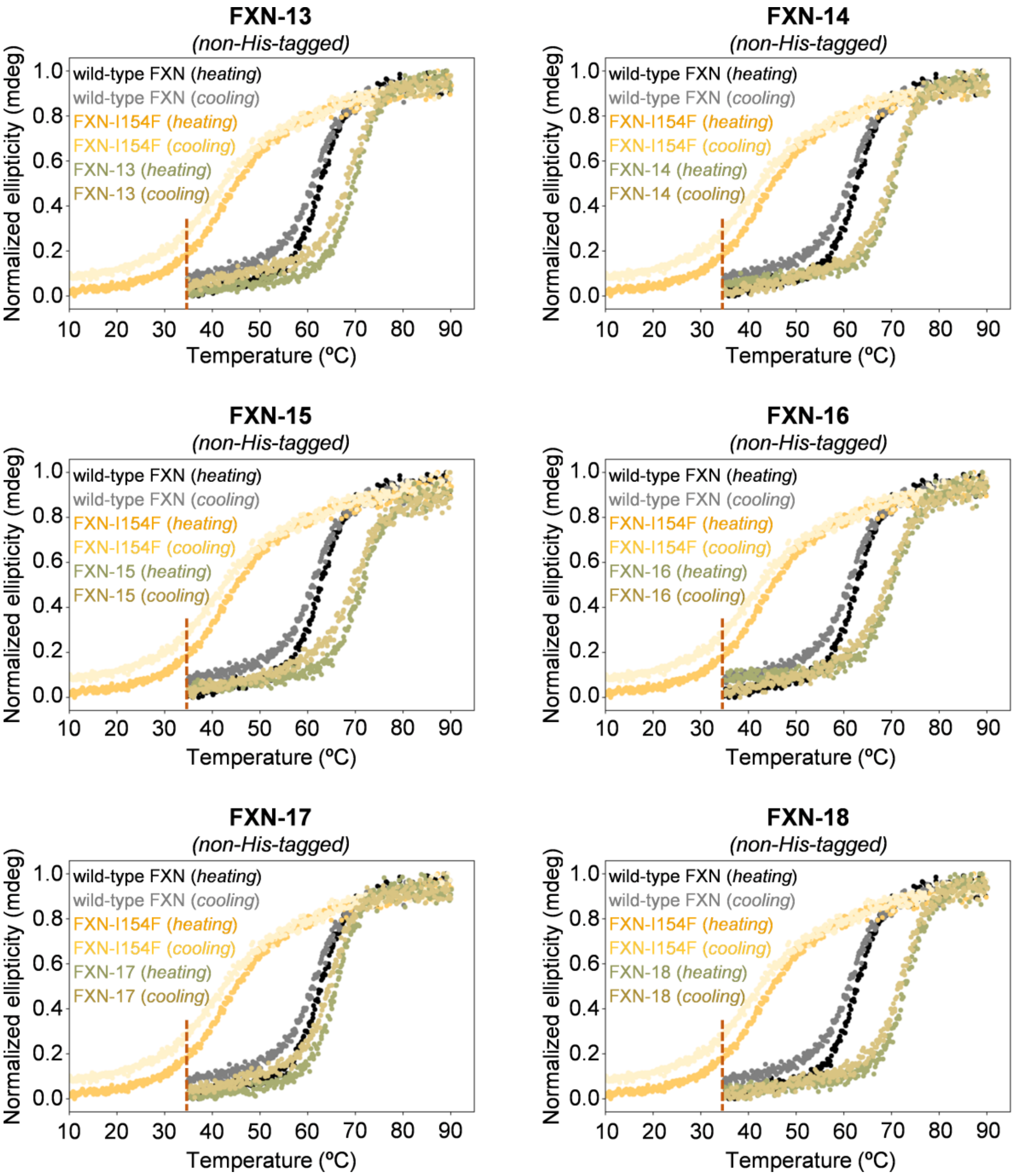


**Figure S9** (cont.). Thermal unfolding of frataxin (FXN) variants measured by circular dichroism (CD) spectroscopy. Reversibility is always measured at 35 ºC for fair comparison across different variants (dashed red line).

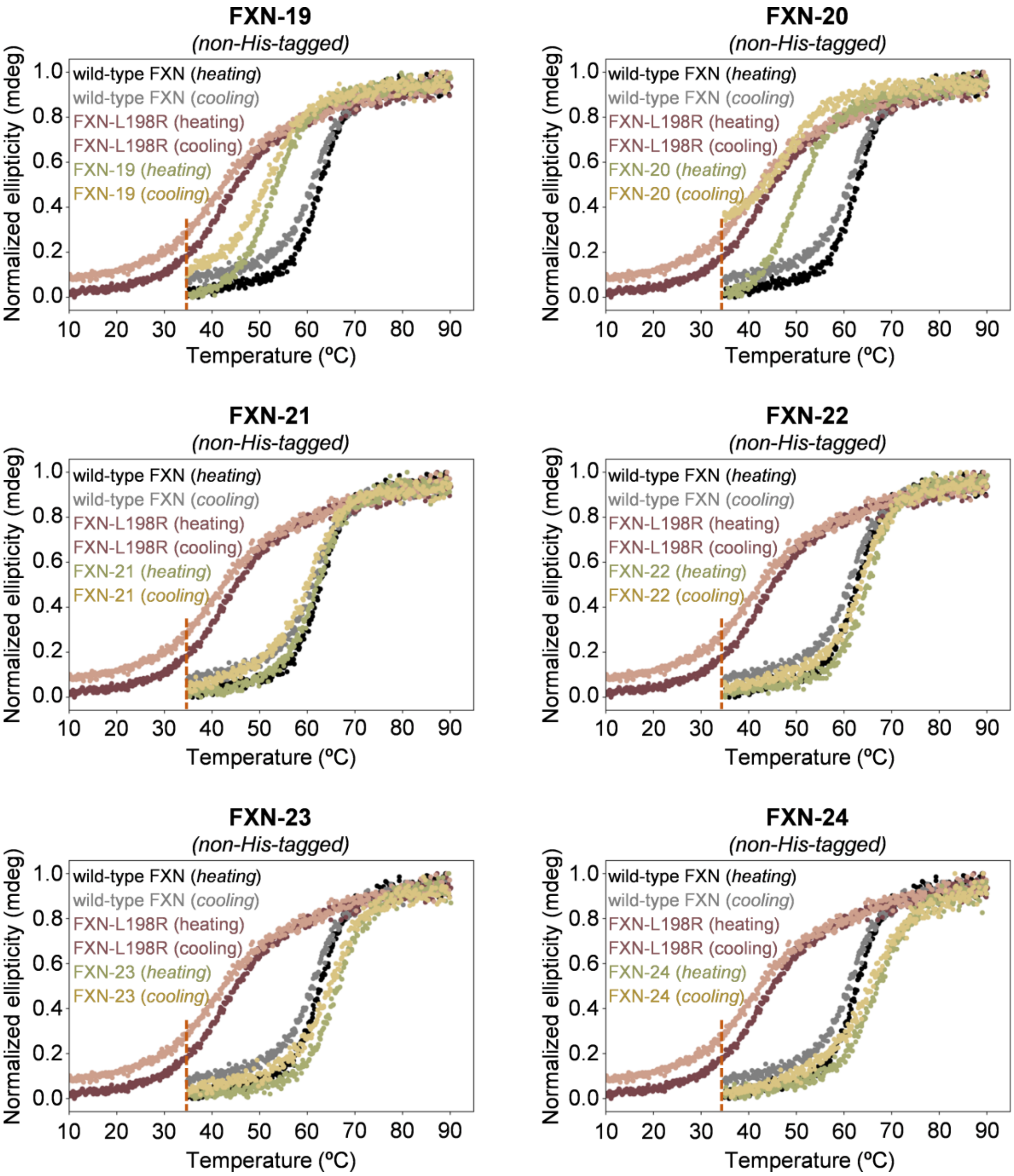


**Figure S9** (cont.). Thermal unfolding of frataxin (FXN) variants measured by circular dichroism (CD) spectroscopy. Reversibility is always measured at 35 ºC for fair comparison across different variants (dashed red line).

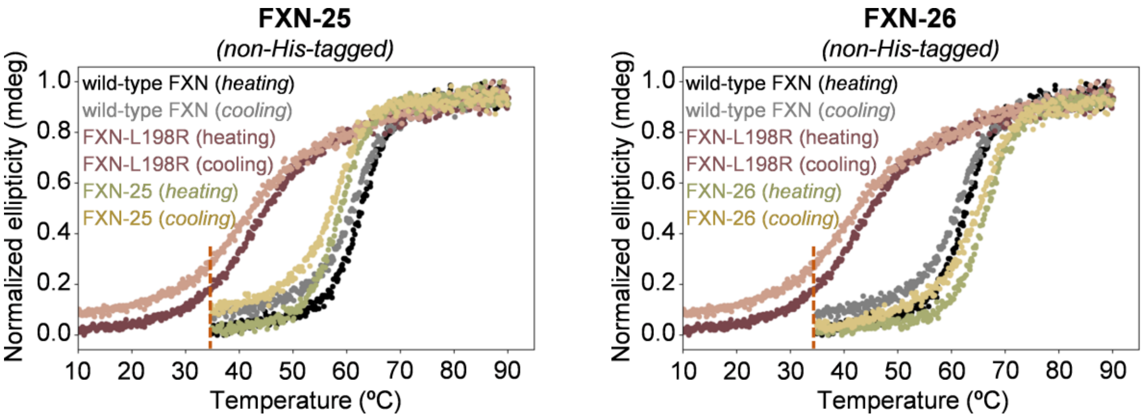


**Figure S9** (cont.). Thermal unfolding of frataxin (FXN) variants measured by circular dichroism (CD) spectroscopy. Reversibility is always measured at 35 ºC for fair comparison across different variants (dashed red line).

### 10. Stability curve determination

Chemical denaturation experiments were performed to obtain the Gibbs-Helmholtz curves ($\Delta G_u$ vs $T$) of wild-type frataxin and FXN-03, FXN-08, and FXN-10. Protein samples were prepared at 1 μM concentration in buffer solution (20 mM HEPES pH 7.4, 120 mM NaCl) in a 10 mm quartz cuvette. Guanidinium hydrochloride (GdnHCl) was used as the denaturant. The concentration of GdnHCl was gradually increased from 0 to 4 M while monitoring the change in ellipticity in the 220-230 nm range using a JASCO J-815 spectropolarimeter equipped with a Peltier temperature control unit. $\Delta G_u$ values derived from chemical denaturation experiments were considered at five temperatures for wild-type frataxin (10, 20, 30, 40 and 50 ºC), three for FXN-10 (10, 30, 50 ºC), and two for FXN-08 and FXN-03 (10 and 30 ºC). Ellipticity values ($E$) vs. denaturant concentration at each temperature were fitted using a least squares algorithm to a two-state model following eq. 12.

$$E = FF \cdot (m_1[D] + b_1) + (1 - FF) \cdot (m_2[D] + b_2) \qquad \text{eq. 12}$$

where $FF$ is the fraction of protein in the native (folded) state, $[D]$ the denaturant concentration, and parameters $m_1$, $b_1$, $m_2$, and $b_2$ the slopes and intercepts of the (linear) ellipticity in the native and denatured state, respectively. $FF$ can be expressed as a function of the free energy of unfolding in the presence of the denaturant $\Delta G_{uD}$, $FF = 1/(1 + e^{-\frac{\Delta G_{uD}}{RT}})$, which is itself a function of the denaturant concentration $\Delta G_{uD}([D]) = \Delta G_u - m[D]$, where $m$ is the denaturant slope and $\Delta G_u$ the free energy of unfolding at the given temperature in the absence of denaturant. Ellipticity data were simultaneously fitted optimizing i) $m_1$, $b_1$, $m_2$, and $b_2$, for each variant and temperature and ii) a common denaturant slope $m$ using an in house Python3 script, obtaining for each variant and temperature the extrapolated $\Delta G_u$ value.

Additional $\Delta G_u$ vs T datapoints were extracted from the CD spectra of thermal denaturation (i.e., $T_m$ measurement) in a symmetric 6 ºC range around the melting temperature (34 datapoints for wild-type frataxin, 31 for FXN-03, 32 for FXN-08, and 30 for FXN-10), where significant populations of the folded and unfolded states thus allowing the calculation of accurate $\Delta G_u$ values.

Finally, to obtain the stability curves the $\Delta G_u$ values were fitted to the Gibbs-Helmholtz equation (eq. 4) with a least squares algorithm optimizing the $\Delta H_m$, $\Delta C_p$ and $T_m$ parameters. $\Delta H_m$ is the enthalpy of unfolding at the melting temperature and $\Delta C_p$ the change in specific heat between denatured and native state.

$$\Delta G_u(T) = \Delta H_m - T\frac{\Delta H_m}{T_m} + \Delta C_p\left[T - T_m - T ln\left(\frac{T}{T_m}\right)\right] \qquad \text{eq. 13}$$

Under the assumption that the stability curves are parallel in the $T_m$ region, approximated changes in stabilization for related variants ($\Delta\Delta G^{appr}$) can be obtained from a simple linear relationship using the $\Delta T_m$ , $\Delta H_m$ and $T_m$ values derived from the stability curve of the wild-type. This approximation assumes a common first derivative of $\Delta G_u(T)$ at the melting temperatures ($\Delta H_m/T_m = \Delta S_m$).

$$\Delta\Delta G^{appr} = \frac{\Delta H_m}{T_m}\Delta T_m \qquad \text{eq. 14}$$

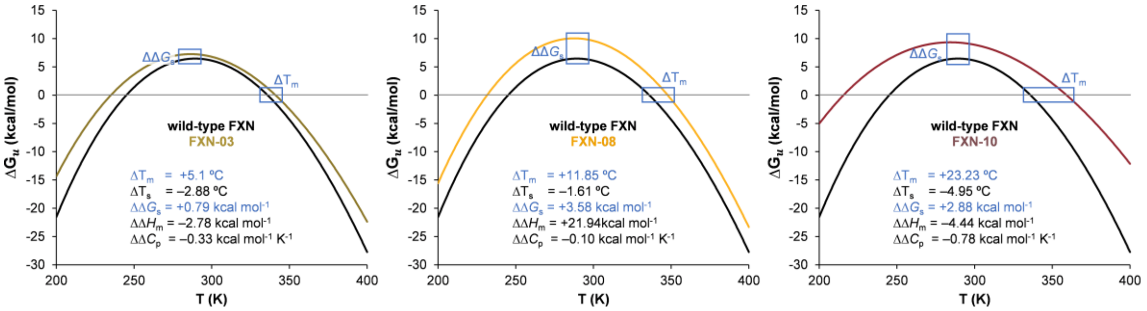


**Figure S10.** Fitted stability curves for wild-type frataxin (black), FXN-03 (green), FXN-08 (yellow) and FXN-10 (red), and changes in the thermodynamic parameters derived from them with respect to those of the wild-type. The differences between the *thermodynamic stabilization* ($\Delta\Delta G_s$) and *thermostabilization* ($\Delta T_m$) measured for each designed variant with respect to the wild-type are highlighted as blue boxes. Note that FXN-08 is the most *thermodynamically stable* variant, while FXN-10 is the most *thermostable* one.

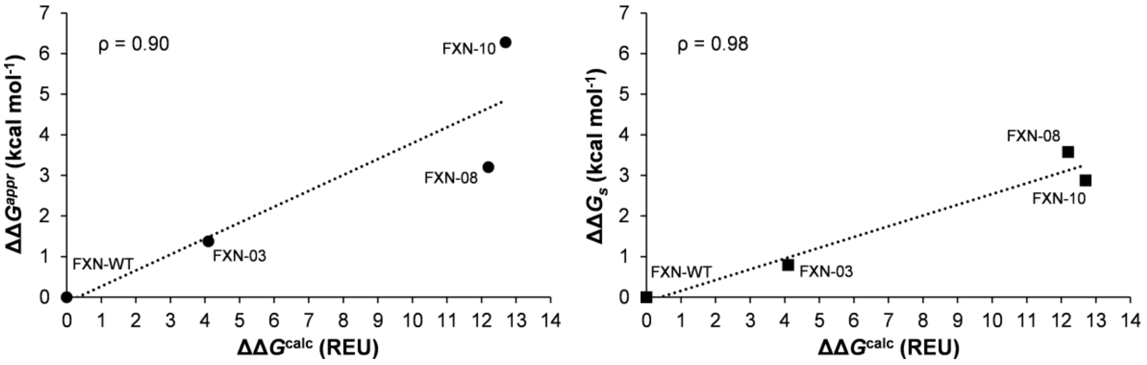


**Figure S11.** Correlation between theoretical $\Delta\Delta G^{calc}$ and experimental $\Delta\Delta G^{appr}$ (approximated using eq. 14; ●) and $\Delta\Delta G_s$ (derived from the stabilization curves; ■) for designed variants FXN-03, FXN-08 and FXN-10, using wild-type frataxin as a reference. Positive values indicate stabilization. ρ: Pearson correlation coefficient. The dashed line indicates a linear regression.

**Table S11.** Experimentally measured *unfolding*/*folding* temperature ($T_m$), differences in *unfolding* $T_m$ ($\Delta T_m$), reversibility, and approximated thermostability changes ($\Delta\Delta G^{appr}$) for wild-type, pathological mutants and designed frataxin variants. $\Delta\Delta G^{est}$ values were derived from $\Delta T_m$ values with respect to wild-type frataxin using eq. 5.

| Variant | $T_{m,u}$ (°C)[a] | $T_{m,f}$ (°C)[b] | rev. (%) | $\Delta T_m$ (°C)[a,c] | $\Delta T_m$ (°C)[a,d] | $\Delta T_m$ (°C)[a,e] | $\Delta\Delta G^{appr}$ (kcal mol$^{-1}$) |
|---|---|---|---|---|---|---|---|
| wild-type FXN | 62.8 | 61.4 | 92.9 | 0.0 | | | 0.00 |
| FXN-01 | 72.6 | 71.8 | 97.5 | +6.1[f] | | | 1.64 |
| FXN-02 | 66.9 | 66.5 | 93.1 | +0.4[f] | | | 0.11 |
| FXN-03 | 67.9 | 66.5 | 86.3 | +5.1 | | | 1.38 |
| FXN-04 | 71.3 | 69.9 | 96.6 | +8.6 | | | 2.32 |
| FXN-05 | 70.0 | 67.6 | 97.0 | +7.3 | | | 1.96 |
| FXN-06 | 67.6 | 65.5 | 94.7 | +4.8 | | | 1.30 |
| FXN-07 | 65.2 | 63.0 | 93.6 | +2.5 | | | 0.67 |
| FXN-08 | 74.6 | 73.5 | 96.4 | +11.9 | | | 3.20 |
| FXN-09 | 75.5 | 73.3 | 99.2 | +12.7 | | | 3.44 |
| FXN-10 | 86.0 | 82.7 | 111.8[g] | +23.3 | | | 6.29 |
| FXN-I154F | 49.9 | 47.3 | 95.1 | –12.9 | 0.0 | | –3.49 |
| FXN-11 | 59.7 | 57.6 | 93.0 | –3.1 | +9.9 | | –0.83 |
| FXN-12 | 56.6 | 54.6 | 82.8 | –6.2 | +6.7 | | –1.68 |
| FXN-13 | 70.2 | 69.2 | 94.3 | +7.4 | +20.3 | | 2.00 |
| FXN-14 | 71.0 | 70.5 | 102.4[g] | +8.2 | +21.1 | | 2.22 |
| FXN-15 | 71.0 | 69.9 | 99.3 | +8.2 | +21.1 | | 2.22 |
| FXN-16 | 70.4 | 69.5 | 106.9[g] | +7.6 | +20.5 | | 2.05 |
| FXN-17 | 66.3 | 65.1 | 98.1 | +3.5 | +16.4 | | 0.95 |
| FXN-18 | 73.3 | 72.4 | 99.7 | +10.5 | +23.4 | | 2.84 |
| FXN-L198R | 42.2 | 40.0 | 89.0 | –20.5 | | 0.0 | –5.56 |
| FXN-19 | 51.5 | 51.2 | 87.7 | –11.2 | | +9.3 | –3.03 |
| FXN-20 | 46.3 | 42.4 | 62.8 | –16.5 | | +4.1 | –4.46 |
| FXN-21 | 62.1 | 60.9 | 96.7 | –0.7 | | +19.8 | –0.19 |
| FXN-22 | 65.1 | 64.0 | 97.6 | +2.4 | | +22.9 | 0.64 |
| FXN-23 | 65.9 | 65.0 | 94.9 | +3.1 | | +23.7 | 0.85 |
| FXN-24 | 66.1 | 65.0 | 101.7[g] | +3.4 | | +23.9 | 0.91 |
| FXN-25 | 58.8 | 58.3 | 91.1 | –4.0 | | +16.6 | –1.08 |
| FXN-26 | 66.3 | 65.5 | 97.9 | +3.5 | | +24.1 | 0.95 |

[a] Measured during forward thermal *unfolding (u)* (i.e. heating). [b] Measured during reverse thermal *folding (f)* (i.e. cooling). [c] Calculated with respect to wild-type FXN. [d] Calculated with respect to FXN-I154F. [e] Calculated with respect to FXN-L198R. [f] The difference in melting temperature ($\Delta T_m$) for these variants is calculated with respect to non-6xHis-tagged wild-type frataxin ($T_{m,u}$ = 66.5; $T_{m,f}$ = 65.9). [g] Reversibility values higher than 100% are due to numerical errors in the calculation of fitted ellipticities, and complete reversibility is assumed.

### 11. Proteolytic resistance assay

#### Mass spectrometry

Wild-type frataxin and the superstable FXN-10 variant (three samples: R1, R2 and R3) were incubated with trypsin at 37 ºC, at an enzyme:protein ratio of 1:100 for 5, 10, 20 and 60 min. The resulting peptides were desalted and resuspended in 0.1% formic acid using C18 stage tips (Millipore). Samples were analyzed in a hybrid trapped ion mobility spectrometry – quadrupole time of flight mass spectrometer (timsTOF Pro with PASEF, Bruker Daltonics) coupled online to a nanoElute liquid chromatograph (Bruker). Each sample (100 ng approx.) was directly loaded in a 15 cm Bruker nanoelute FIFTEEN C18 analytical column (Bruker) and resolved at 400 nl/min with a 100 min gradient. The column was heated to 50 ºC using an oven.

#### Data analysis

Database searching was performed using MASCOT 2.2.07 (Matrixscience) through Proteome Discoverer 1.4 (Thermo) against a Uniprot/Swissprot database consisting of *Homo sapiens* entries, including the frataxin variants of interest. The following parameters were adopted for the searches: carbamidomethylation of cysteines (C) as fixed modification and oxidation of methionines (M) as variable modifications, 20 ppm of peptide mass tolerance, 0.05 Da fragment mass tolerance and up to 2 missed cleavages. Spectral counts, that is, the number of spectra matching to a certain protein in each sample, were used for the assessment of frataxin cleavage levels.

### 12. Mass spectrometry data

| WT_5min_R1 | | | | | | | | | | | | | | |
|---|---|---|---|---|---|---|---|---|---|---|---|---|---|---|
| Accession | Description | Score | Coverage | # Proteins | # Unique Peptides | # Peptides | # PSMs | # AAs | MW [kDa] | calc. pI | | | | |
| Q16595WT10 | FrataxinWT10 OS=Homo sapiens OX=9606 GN=FXN PE=1 SV=2 - [FRDAWT10_HUMAN] | 3284.26 | 88.98 | 2 | 16 | 16 | 91 | 127 | 14.23 | 5.67 | | | | |
| A3 | Sequence | # PSMs | # Proteins | # Protein Groups | Protein Group Accessions | Modifications | ΔCn | IonScore | Exp Value | Charge | MH+ [Da] | ΔM [ppm] | RT [min] | # Missed Cleavages |
| High | LAEETLDSLAEFFEDLADKPYTFEDYDVSFGSGVLTVK | 3 | 2 | 1 | Q16595WT10 | | 0 | 131.35 | 1.46E-12 | 3 | 4261.00 | -6.85 | 18.30 | 0 |
| High | LGGDLGTYVINKQTPNKQIWLSSPSSGPKR | 3 | 2 | 1 | Q16595WT10 | | 0 | 108.94 | 2.55E-10 | 4 | 3241.73 | -0.23 | 11.85 | 3 |
| High | NWVYSHDGVSLHELLAAELTK | 5 | 2 | 1 | Q16595WT10 | | 0 | 106.87 | 5.71E-10 | 3 | 2382.20 | -4.81 | 15.02 | 0 |
| High | LGGDLGTYVINK | 17 | 2 | 1 | Q16595WT10 | | 0 | 100.36 | 2.11E-09 | 2 | 1249.68 | -1.54 | 11.62 | 0 |
| High | TKLDLSSLAYSGKDA | 7 | 2 | 1 | Q16595WT10 | | 0 | 95.54 | 6.55E-09 | 2 | 1568.81 | -5.23 | 11.58 | 2 |
| High | TKLDLSSLAYSGK | 2 | 2 | 1 | Q16595WT10 | | 0 | 87.56 | 3.50E-08 | 2 | 1382.75 | -2.86 | 11.38 | 1 |
| High | LDLSSLAYSGKDA | 4 | 2 | 1 | Q16595WT10 | | 0 | 86.37 | 4.60E-08 | 2 | 1339.67 | -1.66 | 12.68 | 1 |
| High | LDLSSLAYSGK | 3 | 2 | 1 | Q16595WT10 | | 0 | 70.30 | 1.86E-06 | 2 | 1153.60 | -5.87 | 12.62 | 0 |
| High | QIWLSSPSSGPK | 7 | 2 | 1 | Q16595WT10 | | 0 | 62.73 | 1.35E-05 | 2 | 1286.67 | -2.64 | 10.98 | 0 |
| High | QTPNKQIWLSSPSSGPK | 8 | 2 | 1 | Q16595WT10 | | 0 | 59.33 | 2.57E-05 | 3 | 1854.95 | -11.18 | 9.88 | 1 |
| High | YDWTGKNWVYSHDGVSLHELLAAELTK | 6 | 2 | 1 | Q16595WT10 | | 0 | 58.72 | 3.25E-05 | 4 | 3132.53 | -3.49 | 15.38 | 1 |
| High | ALKTKLDLSSLAYSGKDA | 2 | 2 | 1 | Q16595WT10 | | 0 | 57.70 | 3.40E-05 | 3 | 1881.02 | -4.26 | 11.37 | 3 |
| High | RYDWTGKNWVYSHDGVSLHELLAAELTK | 14 | 2 | 1 | Q16595WT10 | | 0 | 57.62 | 4.13E-05 | 5 | 3288.62 | -6.90 | 15.10 | 2 |
| High | QTPNKQIWLSSPSSGPKR | 4 | 2 | 1 | Q16595WT10 | | 0 | 42.79 | 1.20E-03 | 3 | 2011.06 | -3.81 | 9.00 | 2 |
| High | LGGDLGTYVINKQTPNKQIWLSSPSSGPK | 3 | 2 | 1 | Q16595WT10 | | 0 | 39.47 | 2.25E-03 | 4 | 3085.61 | -7.76 | 12.47 | 2 |
| High | QIWLSSPSSGPKR | 3 | 2 | 1 | Q16595WT10 | | 0 | 39.40 | 2.57E-03 | 2 | 1442.77 | -5.59 | 9.78 | 1 |

| WT_5min_R2 | | | | | | | | | | | | | | |
|---|---|---|---|---|---|---|---|---|---|---|---|---|---|---|
| Accession | Description | Score | Coverage | # Proteins | # Unique Peptides | # Peptides | # PSMs | # AAs | MW [kDa] | calc. pI | | | | |
| Q16595WT10 | FrataxinWT10 OS=Homo sapiens OX=9606 GN=FXN PE=1 SV=2 - [FRDAWT10_HUMAN] | 3301.43 | 88.98 | 2 | 16 | 16 | 87 | 127 | 14.23 | 5.67 | | | | |
| A3 | Sequence | # PSMs | # Proteins | # Protein Groups | Protein Group Accessions | Modifications | ΔCn | IonScore | Exp Value | Charge | MH+ [Da] | ΔM [ppm] | RT [min] | # Missed Cleavages |
| High | TKLDLSSLAYSGKDA | 13 | 2 | 1 | Q16595WT10 | | 0 | 111.38 | 1.71E-10 | 2 | 1568.81 | -4.34 | 11.55 | 2 |
| High | LDLSSLAYSGKDA | 8 | 2 | 1 | Q16595WT10 | | 0 | 100.45 | 1.80E-09 | 2 | 1339.67 | -1.95 | 12.70 | 1 |
| High | LGGDLGTYVINK | 25 | 2 | 1 | Q16595WT10 | | 0 | 100.14 | 2.08E-09 | 2 | 1249.67 | -4.71 | 11.55 | 0 |
| High | TKLDLSSLAYSGK | 3 | 2 | 1 | Q16595WT10 | | 0 | 89.88 | 2.05E-08 | 2 | 1382.75 | -4.90 | 11.37 | 1 |
| High | LDLSSLAYSGK | 2 | 2 | 1 | Q16595WT10 | | 0 | 78.94 | 2.55E-07 | 2 | 1153.60 | -8.94 | 12.55 | 0 |
| High | QIWLSSPSSGPK | 10 | 2 | 1 | Q16595WT10 | | 0 | 68.69 | 3.30E-06 | 2 | 1286.67 | -4.89 | 10.92 | 0 |
| High | QTPNKQIWLSSPSSGPK | 6 | 2 | 1 | Q16595WT10 | | 0 | 56.82 | 4.70E-05 | 3 | 1854.97 | -2.47 | 9.83 | 1 |
| High | LAEETLDSLAEFFEDLADKPYTFEDYDVSFGSGVLTVK | 2 | 2 | 1 | Q16595WT10 | | 0 | 45.77 | 5.28E-04 | 3 | 4261.00 | -5.64 | 18.30 | 0 |
| High | RYDWTGKNWVYSHDGVSLHELLAAELTK | 4 | 2 | 1 | Q16595WT10 | | 0 | 45.32 | 7.17E-04 | 5 | 3288.64 | -1.99 | 14.87 | 2 |
| High | ALKTKLDLSSLAYSGKDA | 2 | 2 | 1 | Q16595WT10 | | 0 | 43.52 | 9.29E-04 | 3 | 1881.02 | -5.63 | 11.33 | 3 |
| High | QTPNKQIWLSSPSSGPKR | 2 | 2 | 1 | Q16595WT10 | | 0 | 39.39 | 2.62E-03 | 3 | 2011.05 | -9.22 | 8.90 | 2 |
| High | LGGDLGTYVINKQTPNK | 3 | 2 | 1 | Q16595WT10 | | 0 | 37.27 | 4.72E-03 | 2 | 1817.96 | -7.79 | 10.57 | 1 |
| High | QIWLSSPSSGPKR | 3 | 2 | 1 | Q16595WT10 | | 0 | 37.06 | 4.40E-03 | 2 | 1442.77 | -6.00 | 9.73 | 1 |
| High | YDWTGKNWVYSHDGVSLHELLAAELTK | 1 | 2 | 1 | Q16595WT10 | | 0 | 36.32 | 5.81E-03 | 4 | 3132.54 | -2.38 | 15.38 | 1 |
| High | NWVYSHDGVSLHELLAAELTK | 2 | 2 | 1 | Q16595WT10 | | 0 | 23.78 | 1.18E-01 | 3 | 2382.19 | -6.19 | 14.98 | 0 |
| High | LGGDLGTYVINKQTPNKQIWLSSPSSGPKR | 1 | 2 | 1 | Q16595WT10 | | 0 | 21.75 | 1.33E-01 | 4 | 3241.72 | -4.74 | 11.83 | 3 |

| WT_5min_R3 | | | | | | | | | | | | | | |
|---|---|---|---|---|---|---|---|---|---|---|---|---|---|---|
| Accession | Description | Score | Coverage | # Proteins | # Unique Peptides | # Peptides | # PSMs | # AAs | MW [kDa] | calc. pI | | | | |
| Q16595WT10 | FrataxinWT10 OS=Homo sapiens OX=9606 GN=FXN PE=1 SV=2 - [FRDAWT10_HUMAN] | 5166.11 | 59.06 | 2 | 14 | 14 | 150 | 127 | 14.23 | 5.67 | | | | |
| A3 | Sequence | # PSMs | # Proteins | # Protein Groups | Protein Group Accessions | Modifications | ΔCn | IonScore | Exp Value | Charge | MH+ [Da] | ΔM [ppm] | RT [min] | # Missed Cleavages |
| High | TKLDLSSLAYSGKDA | 35 | 2 | 1 | Q16595WT10 | | 0 | 126.90 | 4.64E-12 | 2 | 1568.81 | -5.74 | 11.55 | 2 |
| High | LGGDLGTYVINK | 47 | 2 | 1 | Q16595WT10 | | 0 | 93.28 | 1.01E-08 | 2 | 1249.67 | -4.65 | 11.57 | 0 |
| High | LDLSSLAYSGK | 3 | 2 | 1 | Q16595WT10 | | 0 | 91.65 | 1.36E-08 | 2 | 1153.61 | -2.56 | 12.55 | 0 |
| High | LGGDLGTYVINKQTPNK | 3 | 2 | 1 | Q16595WT10 | | 0 | 91.64 | 1.63E-08 | 2 | 1817.97 | -1.59 | 10.53 | 1 |
| High | LDLSSLAYSGKDA | 10 | 2 | 1 | Q16595WT10 | | 0 | 87.91 | 3.23E-08 | 2 | 1339.67 | -2.78 | 12.67 | 1 |
| High | TKLDLSSLAYSGK | 3 | 2 | 1 | Q16595WT10 | | 0 | 81.63 | 1.37E-07 | 2 | 1382.75 | -2.47 | 11.37 | 1 |
| High | QTPNKQIWLSSPSSGPK | 12 | 2 | 1 | Q16595WT10 | | 0 | 68.97 | 2.91E-06 | 3 | 1854.96 | -7.09 | 9.78 | 1 |
| High | QIWLSSPSSGPK | 10 | 2 | 1 | Q16595WT10 | | 0 | 63.98 | 1.02E-05 | 2 | 1286.67 | -0.74 | 10.97 | 0 |
| High | ALKTKLDLSSLAYSGKDA | 2 | 2 | 1 | Q16595WT10 | | 0 | 56.32 | 4.75E-05 | 3 | 1881.03 | -4.14 | 11.30 | 3 |
| High | QTPNKQIWLSSPSSGPKR | 19 | 2 | 1 | Q16595WT10 | | 0 | 51.29 | 1.67E-04 | 3 | 2011.06 | -5.57 | 8.95 | 2 |
| High | RYDWTGKNWVYSHDGVSLHELLAAELTK | 1 | 2 | 1 | Q16595WT10 | | 0 | 48.09 | 3.80E-04 | 5 | 3288.63 | -3.09 | 14.87 | 2 |
| High | YDWTGKNWVYSHDGVSLHELLAAELTK | 1 | 2 | 1 | Q16595WT10 | | 0 | 47.38 | 4.11E-04 | 4 | 3132.52 | -6.42 | 15.35 | 1 |
| High | QIWLSSPSSGPKR | 3 | 2 | 1 | Q16595WT10 | | 0 | 42.23 | 1.45E-03 | 2 | 1442.77 | -4.35 | 9.73 | 1 |
| High | LGGDLGTYVINKQTPNKQIWLSSPSSGPK | 1 | 2 | 1 | Q16595WT10 | | 0 | 26.70 | 4.27E-02 | 4 | 3085.62 | -2.71 | 12.43 | 2 |

| WT_1omin_R1 | | | | | | | | | | | | | | |
|---|---|---|---|---|---|---|---|---|---|---|---|---|---|---|
| Accession | Description | Score | Coverage | # Proteins | # Unique Peptides | # Peptides | # PSMs | # AAs | MW [kDa] | calc. pI | | | | |
| Q16595WT10 | FrataxinWT10 OS=Homo sapiens OX=9606 GN=FXN PE=1 SV=2 - [FRDAWT10_HUMAN] | 7373.43 | 59.06 | 2 | 16 | 16 | 223 | 127 | 14.23 | 5.67 | | | | |
| A3 | Sequence | # PSMs | # Proteins | # Protein Groups | Protein Group Accessions | Modifications | ΔCn | IonScore | Exp Value | Charge | MH+ [Da] | ΔM [ppm] | RT [min] | # Missed Cleavages |
| High | TKLDLSSLAYSGKDA | 22 | 2 | 1 | Q16595WT10 | | 0 | 131.60 | 1.61E-12 | 2 | 1568.80 | -8.38 | 11.53 | 2 |
| High | TKLDLSSLAYSGK | 7 | 2 | 1 | Q16595WT10 | | 0 | 110.94 | 1.66E-10 | 2 | 1382.74 | -9.41 | 11.32 | 1 |
| High | NWVYSHDGVSLHELLAAELTK | 10 | 2 | 1 | Q16595WT10 | | 0 | 92.98 | 1.35E-08 | 3 | 2382.18 | -11.18 | 14.95 | 0 |
| High | LGGDLGTYVINK | 69 | 2 | 1 | Q16595WT10 | | 0 | 92.29 | 1.35E-08 | 2 | 1249.67 | -8.22 | 11.48 | 0 |
| High | LDLSSLAYSGK | 6 | 2 | 1 | Q16595WT10 | | 0 | 90.40 | 1.66E-08 | 2 | 1153.60 | -11.54 | 12.50 | 0 |
| High | LGGDLGTYVINKQTPNK | 3 | 2 | 1 | Q16595WT10 | | 0 | 86.73 | 5.48E-08 | 2 | 1817.96 | -9.96 | 10.50 | 1 |
| High | LDLSSLAYSGKDA | 9 | 2 | 1 | Q16595WT10 | | 0 | 86.37 | 4.04E-08 | 2 | 1339.67 | -6.13 | 12.67 | 1 |
| High | QTPNKQIWLSSPSSGPK | 18 | 2 | 1 | Q16595WT10 | | 0 | 81.75 | 1.53E-07 | 3 | 1854.96 | -6.87 | 9.78 | 1 |
| High | QTPNKQIWLSSPSSGPKR | 15 | 2 | 1 | Q16595WT10 | | 0 | 75.56 | 6.21E-07 | 3 | 2011.05 | -10.31 | 8.95 | 2 |
| High | LGGDLGTYVINKQTPNKQIWLSSPSSGPKR | 2 | 2 | 1 | Q16595WT10 | | 0 | 74.50 | 6.21E-07 | 5 | 3241.71 | -7.01 | 11.78 | 3 |
| High | QIWLSSPSSGPK | 27 | 2 | 1 | Q16595WT10 | | 0 | 68.65 | 3.25E-06 | 2 | 1286.66 | -7.21 | 10.90 | 0 |
| High | QIWLSSPSSGPKR | 24 | 2 | 1 | Q16595WT10 | | 0 | 59.63 | 2.47E-05 | 2 | 1442.76 | -11.10 | 9.67 | 1 |
| High | YDWTGKNWVYSHDGVSLHELLAAELTK | 1 | 2 | 1 | Q16595WT10 | | 0 | 58.06 | 3.22E-05 | 4 | 3132.51 | -11.24 | 15.35 | 1 |
| High | ALKTKLDLSSLAYSGKDA | 3 | 2 | 1 | Q16595WT10 | | 0 | 57.93 | 3.66E-05 | 3 | 1881.02 | -9.45 | 11.30 | 3 |
| High | LGGDLGTYVINKQTPNKQIWLSSPSSGPK | 3 | 2 | 1 | Q16595WT10 | | 0 | 51.69 | 1.36E-04 | 4 | 3085.60 | -9.45 | 12.40 | 2 |
| High | RYDWTGK | 4 | 2 | 1 | Q16595WT10 | | 0 | 25.89 | 3.28E-02 | 2 | 925.45 | -7.40 | 7.35 | 1 |

| WT_1omin_R2 | | | | | | | | | | | | | | |
|---|---|---|---|---|---|---|---|---|---|---|---|---|---|---|
| Accession | Description | Score | Coverage | # Proteins | # Unique Peptides | # Peptides | # PSMs | # AAs | MW [kDa] | calc. pI | | | | |
| Q16595WT10 | FrataxinWT10 OS=Homo sapiens OX=9606 GN=FXN PE=1 SV=2 - [FRDAWT10_HUMAN] | 7768.90 | 59.06 | 2 | 3 | 17 | 235 | 127 | 14.23 | 5.67 | | | | |
| A3 | Sequence | # PSMs | # Proteins | # Protein Groups | Protein Group Accessions | Modifications | ΔCn | IonScore | Exp Value | Charge | MH+ [Da] | ΔM [ppm] | RT [min] | # Missed Cleavages |
| High | TKLDLSSLAYSGKDA | 34 | 3 | 2 | Q16595WT10; Q16595WT1 | | 0 | 127.03 | 4.51E-12 | 2 | 1568.81 | -6.09 | 11.48 | 2 |
| High | LDLSSLAYSGKDA | 15 | 3 | 2 | Q16595WT10; Q16595WT1 | | 0 | 100.66 | 1.53E-09 | 2 | 1339.66 | -7.06 | 12.65 | 1 |
| High | LGGDLGTYVINK | 86 | 3 | 2 | Q16595WT10; Q16595WT1 | | 0 | 100.58 | 1.90E-09 | 2 | 1249.67 | -6.06 | 11.60 | 0 |
| High | QTPNKQIWLSSPSSGPK | 18 | 3 | 2 | Q16595WT10; Q16595WT1 | | 0 | 94.85 | 7.50E-09 | 3 | 1854.96 | -6.46 | 9.78 | 1 |
| High | LDLSSLAYSGK | 4 | 3 | 2 | Q16595WT10; Q16595WT1 | | 0 | 78.85 | 2.51E-07 | 2 | 1153.60 | -6.20 | 12.53 | 0 |
| High | QTPNKQIWLSSPSSGPKR | 9 | 3 | 2 | Q16595WT10; Q16595WT1 | | 0 | 77.22 | 4.32E-07 | 3 | 2011.06 | -8.46 | 8.95 | 2 |
| High | YDWTGKNWVYSHDGVSLHELLAAELTK | 1 | 2 | 1 | Q16595WT10 | | 0 | 75.53 | 6.24E-07 | 4 | 3132.52 | -7.13 | 15.32 | 1 |
| High | QIWLSSPSSGPK | 22 | 3 | 2 | Q16595WT10; Q16595WT1 | | 0 | 75.49 | 6.58E-07 | 2 | 1286.66 | -9.22 | 11.08 | 0 |
| High | TKLDLSSLAYSGK | 8 | 3 | 2 | Q16595WT10; Q16595WT1 | | 0 | 74.81 | 6.90E-07 | 2 | 1382.74 | -8.23 | 11.33 | 1 |
| High | ALKTKLDLSSLAYSGKDA | 4 | 3 | 2 | Q16595WT10; Q16595WT1 | | 0 | 70.11 | 2.23E-06 | 3 | 1881.02 | -8.91 | 11.28 | 3 |
| High | LGGDLGTYVINKQTPNKQIWLSSPSSGPK | 2 | 3 | 2 | Q16595WT10; Q16595WT1 | | 0 | 56.48 | 5.05E-05 | 4 | 3085.59 | -13.62 | 12.40 | 2 |
| High | LGGDLGTYVINKQTPNK | 2 | 3 | 2 | Q16595WT10; Q16595WT1 | | 0 | 53.54 | 1.10E-04 | 2 | 1817.96 | -6.78 | 10.50 | 1 |
| High | NWVYSHDGVSLHELLAAELTK | 4 | 2 | 1 | Q16595WT10 | | 0 | 52.52 | 1.60E-04 | 3 | 2382.19 | -7.98 | 14.90 | 0 |
| High | QIWLSSPSSGPKR | 21 | 3 | 2 | Q16595WT10; Q16595WT1 | | 0 | 48.45 | 3.29E-04 | 2 | 1442.76 | -8.80 | 10.15 | 1 |
| High | LGGDLGTYVINKQTPNKQIWLSSPSSGPKR | 2 | 3 | 2 | Q16595WT10; Q16595WT1 | | 0 | 41.66 | 1.26E-03 | 4 | 3241.70 | -10.05 | 11.75 | 3 |
| High | RYDWTGKNWVYSHDGVSLHELLAAELTK | 1 | 2 | 1 | Q16595WT10 | | 0 | 31.77 | 1.52E-02 | 5 | 3288.62 | -8.41 | 14.77 | 2 |
| High | RYDWTGK | 2 | 3 | 2 | Q16595WT10; Q16595WT1 | | 0 | 22.56 | 8.03E-02 | 2 | 925.45 | -7.49 | 7.37 | 1 |

| WT_1omin_R3 | | | | | | | | | | | | | | |
|---|---|---|---|---|---|---|---|---|---|---|---|---|---|---|
| Accession | Description | Score | Coverage | # Proteins | # Unique Peptides | # Peptides | # PSMs | # AAs | MW [kDa] | calc. pI | | | | |
| Q16595WT10 | FrataxinWT10 OS=Homo sapiens OX=9606 GN=FXN PE=1 SV=2 - [FRDAWT10_HUMAN] | 5836.22 | 88.98 | 2 | 15 | 15 | 142 | 127 | 14.23 | 5.67 | | | | |
| A3 | Sequence | # PSMs | # Proteins | # Protein Groups | Protein Group Accessions | Modifications | ΔCn | IonScore | Exp Value | Charge | MH+ [Da] | ΔM [ppm] | RT [min] | # Missed Cleavages |
| High | TKLDLSSLAYSGKDA | 13 | 2 | 1 | Q16595WT10 | | 0 | 137.07 | 4.56E-13 | 2 | 1568.80 | -8.38 | 11.52 | 2 |
| High | LAEETLDSLAEFFEDLADKPYTFEDYDVSFGSGVLTVK | 2 | 2 | 1 | Q16595WT10 | | 0 | 126.55 | 4.03E-12 | 3 | 4260.98 | -10.41 | 18.30 | 0 |
| High | NWVYSHDGVSLHELLAAELTK | 6 | 2 | 1 | Q16595WT10 | | 0 | 100.38 | 2.55E-09 | 3 | 2382.18 | -10.15 | 14.95 | 0 |
| High | LGGDLGTYVINK | 36 | 2 | 1 | Q16595WT10 | | 0 | 93.31 | 1.09E-08 | 2 | 1249.67 | -10.11 | 11.47 | 0 |
| High | TKLDLSSLAYSGK | 4 | 2 | 1 | Q16595WT10 | | 0 | 85.64 | 5.83E-08 | 2 | 1382.74 | -8.46 | 11.33 | 1 |
| High | LDLSSLAYSGK | 5 | 2 | 1 | Q16595WT10 | | 0 | 84.98 | 5.78E-08 | 2 | 1153.60 | -7.48 | 12.58 | 0 |
| High | YDWTGKNWVYSHDGVSLHELLAAELTK | 3 | 2 | 1 | Q16595WT10 | | 0 | 78.11 | 3.31E-07 | 4 | 3132.52 | -8.40 | 15.30 | 1 |
| High | LDLSSLAYSGKDA | 13 | 2 | 1 | Q16595WT10 | | 0 | 73.90 | 7.41E-07 | 2 | 1339.66 | -11.78 | 13.52 | 1 |
| High | QIWLSSPSSGPK | 31 | 2 | 1 | Q16595WT10 | | 0 | 69.46 | 2.70E-06 | 2 | 1286.67 | -6.36 | 10.87 | 0 |
| High | QIWLSSPSSGPKR | 5 | 2 | 1 | Q16595WT10 | | 0 | 64.78 | 7.50E-06 | 2 | 1442.76 | -8.05 | 9.73 | 1 |
| High | QTPNKQIWLSSPSSGPKR | 3 | 2 | 1 | Q16595WT10 | | 0 | 63.32 | 1.06E-05 | 3 | 2011.05 | -9.07 | 8.90 | 2 |
| High | RYDWTGKNWVYSHDGVSLHELLAAELTK | 7 | 2 | 1 | Q16595WT10 | | 0 | 61.83 | 1.50E-05 | 4 | 3288.61 | -10.41 | 14.78 | 2 |
| High | ALKTKLDLSSLAYSGKDA | 3 | 2 | 1 | Q16595WT10 | | 0 | 60.35 | 1.90E-05 | 3 | 1881.02 | -6.07 | 11.30 | 3 |
| High | QTPNKQIWLSSPSSGPK | 9 | 2 | 1 | Q16595WT10 | | 0 | 59.07 | 2.89E-05 | 3 | 1854.96 | -7.58 | 9.80 | 1 |
| High | LGGDLGTYVINKQTPNK | 2 | 2 | 1 | Q16595WT10 | | 0 | 33.38 | 1.18E-02 | 2 | 1817.95 | -12.57 | 10.52 | 1 |

| WT_2omin_R1 | | | | | | | | | | | | | | |
|---|---|---|---|---|---|---|---|---|---|---|---|---|---|---|
| Accession | Description | Score | Coverage | # Proteins | # Unique Peptides | # Peptides | # PSMs | # AAs | MW [kDa] | calc. pI | | | | |
| Q16595WT10 | FrataxinWT10 OS=Homo sapiens OX=9606 GN=FXN PE=1 SV=2 - [FRDAWT10_HUMAN] | 6583.22 | 59.06 | 2 | 15 | 15 | 193 | 127 | 14.23 | 5.67 | | | | |
| A3 | Sequence | # PSMs | # Proteins | # Protein Groups | Protein Group Accessions | Modifications | ΔCn | IonScore | Exp Value | Charge | MH+ [Da] | ΔM [ppm] | RT [min] | # Missed Cleavages |
| High | TKLDLSSLAYSGKDA | 28 | 2 | 1 | Q16595WT10 | | 0 | 124.16 | 9.04E-12 | 2 | 1568.81 | -5.31 | 11.50 | 2 |
| High | LGGDLGTYVINK | 66 | 2 | 1 | Q16595WT10 | | 0 | 100.23 | 2.06E-09 | 2 | 1249.67 | -5.82 | 11.62 | 0 |
| High | TKLDLSSLAYSGK | 8 | 2 | 1 | Q16595WT10 | | 0 | 95.20 | 5.84E-09 | 2 | 1382.75 | -2.82 | 11.35 | 1 |
| High | LDLSSLAYSGKDA | 11 | 2 | 1 | Q16595WT10 | | 0 | 94.17 | 6.87E-09 | 2 | 1339.67 | -4.39 | 12.68 | 1 |
| High | LDLSSLAYSGK | 5 | 2 | 1 | Q16595WT10 | | 0 | 91.83 | 1.26E-08 | 2 | 1153.60 | -5.78 | 12.53 | 0 |
| High | QTPNKQIWLSSPSSGPKR | 5 | 2 | 1 | Q16595WT10 | | 0 | 84.24 | 8.65E-08 | 3 | 2011.06 | -7.48 | 8.90 | 2 |
| High | LGGDLGTYVINKQTPNK | 4 | 2 | 1 | Q16595WT10 | | 0 | 76.78 | 5.22E-07 | 2 | 1817.96 | -6.95 | 10.57 | 1 |
| High | QIWLSSPSSGPK | 26 | 2 | 1 | Q16595WT10 | | 0 | 75.59 | 6.58E-07 | 2 | 1286.66 | -7.10 | 10.88 | 0 |
| High | QIWLSSPSSGPKR | 17 | 2 | 1 | Q16595WT10 | | 0 | 73.18 | 1.08E-06 | 2 | 1442.77 | -6.21 | 10.20 | 1 |
| High | QTPNKQIWLSSPSSGPK | 14 | 2 | 1 | Q16595WT10 | | 0 | 65.88 | 5.89E-06 | 3 | 1854.96 | -6.49 | 9.82 | 1 |
| High | YDWTGKNWVYSHDGVSLHELLAAELTK | 1 | 2 | 1 | Q16595WT10 | | 0 | 52.49 | 1.27E-04 | 4 | 3132.52 | -6.66 | 15.32 | 1 |
| High | NWVYSHDGVSLHELLAAELTK | 3 | 2 | 1 | Q16595WT10 | | 0 | 51.42 | 2.04E-04 | 3 | 2382.19 | -9.41 | 14.93 | 0 |
| High | ALKTKLDLSSLAYSGKDA | 1 | 2 | 1 | Q16595WT10 | | 0 | 44.01 | 8.14E-04 | 3 | 1881.02 | -5.98 | 11.30 | 3 |
| High | RYDWTGKNWVYSHDGVSLHELLAAELTK | 2 | 2 | 1 | Q16595WT10 | | 0 | 38.02 | 3.66E-03 | 5 | 3288.61 | -9.28 | 14.73 | 2 |
| High | RYDWTGK | 2 | 2 | 1 | Q16595WT10 | | 0 | 24.78 | 4.36E-02 | 2 | 925.45 | -3.81 | 7.35 | 1 |

| WT_2omin_R2 | | | | | | | | | | | | | | |
|---|---|---|---|---|---|---|---|---|---|---|---|---|---|---|
| Accession | Description | Score | Coverage | # Proteins | # Unique Peptides | # Peptides | # PSMs | # AAs | MW [kDa] | calc. pI | | | | |
| Q16595WT10 | FrataxinWT10 OS=Homo sapiens OX=9606 GN=FXN PE=1 SV=2 - [FRDAWT10_HUMAN] | 7735.54 | 88.98 | 2 | 17 | 17 | 224 | 127 | 14.23 | 5.67 | | | | |
| A3 | Sequence | # PSMs | # Proteins | # Protein Groups | Protein Group Accessions | Modifications | ΔCn | IonScore | Exp Value | Charge | MH+ [Da] | ΔM [ppm] | RT [min] | # Missed Cleavages |
| High | LAEETLDSLAEFFEDLADKPYTFEDYDVSFGSGVLTVK | 3 | 2 | 1 | Q16595WT10 | | 0 | 135.62 | 5.15E-13 | 3 | 4260.99 | -8.08 | 18.30 | 0 |
| High | TKLDLSSLAYSGKDA | 22 | 2 | 1 | Q16595WT10 | | 0 | 124.16 | 9.19E-12 | 2 | 1568.81 | -3.06 | 11.50 | 2 |
| High | LDLSSLAYSGKDA | 16 | 2 | 1 | Q16595WT10 | | 0 | 100.71 | 1.60E-09 | 2 | 1339.67 | -4.67 | 12.73 | 1 |
| High | TKLDLSSLAYSGK | 11 | 2 | 1 | Q16595WT10 | | 0 | 100.53 | 1.82E-09 | 2 | 1382.75 | -3.44 | 11.32 | 1 |
| High | LGGDLGTYVINK | 77 | 2 | 1 | Q16595WT10 | | 0 | 96.95 | 4.35E-09 | 2 | 1249.67 | -4.38 | 11.52 | 0 |
| High | LDLSSLAYSGK | 6 | 2 | 1 | Q16595WT10 | | 0 | 79.82 | 2.01E-07 | 2 | 1153.60 | -5.99 | 12.52 | 0 |
| High | ALKTKLDLSSLAYSGKDA | 10 | 2 | 1 | Q16595WT10 | | 0 | 79.55 | 2.33E-07 | 3 | 1881.02 | -5.21 | 11.30 | 3 |
| High | NWVYSHDGVSLHELLAAELTK | 4 | 2 | 1 | Q16595WT10 | | 0 | 75.36 | 8.08E-07 | 3 | 2382.20 | -5.29 | 14.97 | 0 |
| High | QTPNKQIWLSSPSSGPK | 11 | 2 | 1 | Q16595WT10 | | 0 | 73.72 | 9.60E-07 | 3 | 1854.96 | -4.94 | 9.78 | 1 |
| High | QTPNKQIWLSSPSSGPKR | 10 | 2 | 1 | Q16595WT10 | | 0 | 68.36 | 3.35E-06 | 3 | 2011.05 | -8.77 | 8.88 | 2 |
| High | QIWLSSPSSGPK | 30 | 2 | 1 | Q16595WT10 | | 0 | 59.33 | 2.84E-05 | 2 | 1286.67 | -5.07 | 12.00 | 0 |
| High | LGGDLGTYVINKQTPNKQIWLSSPSSGPKR | 2 | 2 | 1 | Q16595WT10 | | 0 | 50.62 | 1.67E-04 | 4 | 3241.70 | -11.69 | 11.78 | 3 |
| High | QIWLSSPSSGPKR | 11 | 2 | 1 | Q16595WT10 | | 0 | 50.54 | 1.96E-04 | 2 | 1442.76 | -7.21 | 9.72 | 1 |
| High | LGGDLGTYVINKQTPNKQIWLSSPSSGPK | 3 | 2 | 1 | Q16595WT10 | | 0 | 47.95 | 3.01E-04 | 4 | 3085.62 | -2.84 | 12.43 | 2 |
| High | RYDWTGKNWVYSHDGVSLHELLAAELTK | 1 | 2 | 1 | Q16595WT10 | | 0 | 45.58 | 6.72E-04 | 5 | 3288.62 | -5.80 | 14.73 | 2 |
| High | RYDWTGK | 5 | 2 | 1 | Q16595WT10 | | 0 | 43.23 | 8.93E-04 | 2 | 925.45 | -7.66 | 7.35 | 1 |
| High | LGGDLGTYVINKQTPNK | 2 | 2 | 1 | Q16595WT10 | | 0 | 39.32 | 2.89E-03 | 2 | 1817.96 | -6.03 | 10.53 | 1 |

| WT_2omin_R3 | | | | | | | | | | | | | | |
|---|---|---|---|---|---|---|---|---|---|---|---|---|---|---|
| Accession | Description | Score | Coverage | # Proteins | # Unique Peptides | # Peptides | # PSMs | # AAs | MW [kDa] | calc. pI | | | | |
| Q16595WT10 | FrataxinWT10 OS=Homo sapiens OX=9606 GN=FXN PE=1 SV=2 - [FRDAWT10_HUMAN] | 8571.61 | 88.98 | 2 | 19 | 19 | 255 | 127 | 14.23 | 5.67 | | | | |
| A3 | Sequence | # PSMs | # Proteins | # Protein Groups | Protein Group Accessions | Modifications | ΔCn | IonScore | Exp Value | Charge | MH+ [Da] | ΔM [ppm] | RT [min] | # Missed Cleavages |
| High | NWVYSHDGVSLHELLAAELTK | 10 | 2 | 1 | Q16595WT10 | | 0 | 139.52 | 3.12E-13 | 3 | 2382.20 | -4.61 | 14.97 | 0 |
| High | TKLDLSSLAYSGKDA | 17 | 2 | 1 | Q16595WT10 | | 0 | 127.08 | 4.60E-12 | 2 | 1568.81 | -3.45 | 11.55 | 2 |
| High | LAEETLDSLAEFFEDLADKPYTFEDYDVSFGSGVLTVK | 2 | 2 | 1 | Q16595WT10 | | 0 | 101.72 | 8.51E-10 | 3 | 4261.00 | -6.87 | 18.30 | 0 |
| High | LGGDLGTYVINK | 32 | 2 | 1 | Q16595WT10 | | 0 | 99.63 | 2.47E-09 | 2 | 1249.67 | -7.93 | 11.55 | 0 |
| High | QTPNKQIWLSSPSSGPK | 21 | 2 | 1 | Q16595WT10 | | 0 | 94.33 | 8.17E-09 | 3 | 1854.97 | -2.76 | 9.78 | 1 |
| High | ALKTKLDLSSLAYSGKDA | 20 | 2 | 1 | Q16595WT10 | | 0 | 93.52 | 8.45E-09 | 3 | 1881.03 | -2.41 | 11.30 | 3 |
| High | LDLSSLAYSGKDA | 30 | 2 | 1 | Q16595WT10 | | 0 | 93.47 | 8.07E-09 | 2 | 1339.67 | -4.55 | 12.67 | 1 |
| High | LDLSSLAYSGK | 8 | 2 | 1 | Q16595WT10 | | 0 | 84.62 | 6.68E-08 | 2 | 1153.60 | -4.71 | 12.50 | 0 |
| High | QTPNKQIWLSSPSSGPKR | 17 | 2 | 1 | Q16595WT10 | | 0 | 84.07 | 8.81E-08 | 3 | 2011.06 | -4.97 | 8.88 | 2 |
| High | TKLDLSSLAYSGK | 6 | 2 | 1 | Q16595WT10 | | 0 | 83.26 | 9.09E-08 | 2 | 1382.75 | -1.82 | 11.33 | 1 |
| High | YDWTGKNWVYSHDGVSLHELLAAELTK | 1 | 2 | 1 | Q16595WT10 | | 0 | 81.04 | 1.77E-07 | 4 | 3132.52 | -6.52 | 15.32 | 1 |
| High | LGGDLGTYVINKQTPNK | 3 | 2 | 1 | Q16595WT10 | | 0 | 76.24 | 5.87E-07 | 2 | 1817.96 | -6.25 | 10.52 | 1 |
| High | LGGDLGTYVINKQTPNKQIWLSSPSSGPK | 3 | 2 | 1 | Q16595WT10 | | 0 | 73.92 | 7.93E-07 | 4 | 3085.61 | -7.36 | 12.42 | 2 |
| High | LGGDLGTYVINKQTPNKQIWLSSPSSGPKR | 2 | 2 | 1 | Q16595WT10 | | 0 | 71.35 | 1.13E-06 | 5 | 3241.73 | -1.84 | 11.78 | 3 |
| High | QIWLSSPSSGPK | 56 | 2 | 1 | Q16595WT10 | | 0 | 65.60 | 6.57E-06 | 2 | 1286.67 | -6.85 | 10.90 | 0 |
| High | ALKTKLDLSSLAYSGK | 5 | 2 | 1 | Q16595WT10 | | 0 | 59.22 | 1.50E-05 | 2 | 1694.96 | -4.52 | 11.18 | 2 |
| High | RYDWTGKNWVYSHDGVSLHELLAAELTK | 8 | 2 | 1 | Q16595WT10 | | 0 | 52.56 | 1.28E-04 | 4 | 3288.62 | -7.54 | 14.77 | 2 |
| High | QIWLSSPSSGPKR | 9 | 2 | 1 | Q16595WT10 | | 0 | 51.93 | 1.47E-04 | 2 | 1442.76 | -8.55 | 9.67 | 1 |
| High | RYDWTGK | 5 | 2 | 1 | Q16595WT10 | | 0 | 32.70 | 6.25E-03 | 2 | 925.44 | -11.03 | 7.33 | 1 |

| WT_1h_R1 | | | | | | | | | | | | | | |
|---|---|---|---|---|---|---|---|---|---|---|---|---|---|---|
| Accession | Description | Score | Coverage | # Proteins | # Unique Peptides | # Peptides | # PSMs | # AAs | MW [kDa] | calc. pI | | | | |
| Q16595WT10 | FrataxinWT10 OS=Homo sapiens OX=9606 GN=FXN PE=1 SV=2 - [FRDAWT10_HUMAN] | 8121.60 | 88.98 | 2 | 1 | 14 | 226 | 127 | 14.23 | 5.67 | | | | |
| A3 | Sequence | # PSMs | # Proteins | # Protein Groups | Protein Group Accessions | Modifications | ΔCn | IonScore | Exp Value | Charge | MH+ [Da] | ΔM [ppm] | RT [min] | # Missed Cleavages |
| High | TKLDLSSLAYSGKDA | 45 | 3 | 2 | Q16595WT10 Q16595WT1 | | 0 | 123.28 | 1.07E-11 | 2 | 1568.82 | 0.87 | 11.52 | 2 |
| High | TKLDLSSLAYSGK | 12 | 3 | 2 | Q16595WT10 Q16595WT1 | | 0 | 115.29 | 5.68E-11 | 2 | 1382.75 | -1.34 | 11.32 | 1 |
| High | LDLSSLAYSGKDA | 20 | 3 | 2 | Q16595WT10 Q16595WT1 | | 0 | 115.16 | 5.75E-11 | 2 | 1339.68 | 0.75 | 12.63 | 1 |
| High | LGGDLGTYVINK | 50 | 4 | 3 | Q16595WT10 Q16595WT1 Q16595D10 | | 0 | 102.76 | 1.22E-09 | 2 | 1249.68 | -0.85 | 11.50 | 0 |
| High | QTPNKQIWLSSPSSGPK | 14 | 3 | 2 | Q16595WT10 Q16595WT1 | | 0 | 94.18 | 8.61E-09 | 3 | 1854.96 | -3.50 | 9.78 | 1 |
| High | LDLSSLAYSGK | 16 | 3 | 2 | Q16595WT10 Q16595WT1 | | 0 | 91.86 | 1.25E-08 | 2 | 1153.61 | -2.42 | 12.57 | 0 |
| High | ALKTKLDLSSLAYSGKDA | 3 | 3 | 2 | Q16595WT10 Q16595WT1 | | 0 | 74.35 | 7.09E-07 | 3 | 1881.03 | -2.04 | 11.32 | 3 |
| High | QIWLSSPSSGPK | 45 | 3 | 2 | Q16595WT10 Q16595WT1 | | 0 | 71.94 | 1.65E-06 | 2 | 1286.67 | 0.22 | 10.87 | 0 |
| High | LAEETLDSLAEFFEDLADKPYTFEDYDVSFGSGVLTVK | 2 | 3 | 2 | Q16595WT10 Q16595WT1 | | 0 | 67.50 | 2.42E-06 | 3 | 4261.01 | -4.16 | 18.30 | 0 |
| High | LGGDLGTYVINKQTPNK | 3 | 4 | 3 | Q16595WT10 Q16595WT1Q16595D10 | | 0 | 67.48 | 4.31E-06 | 2 | 1817.97 | -2.60 | 10.55 | 1 |
| High | QTPNKQIWLSSPSSGPKR | 3 | 3 | 2 | Q16595WT10 Q16595WT1 | | 0 | 58.25 | 3.36E-05 | 3 | 2011.07 | -0.60 | 8.87 | 2 |
| High | NWVYSHDGVSLHELLAAELTK | 3 | 2 | 1 | Q16595WT10 | | 0 | 51.02 | 2.19E-04 | 3 | 2382.21 | -0.20 | 14.93 | 0 |
| High | QIWLSSPSSGPKR | 6 | 3 | 2 | Q16595WT10 Q16595WT1 | | 0 | 50.94 | 1.99E-04 | 2 | 1442.77 | -1.15 | 10.57 | 1 |
| High | RYDWTGK | 4 | 3 | 2 | Q16595WT10 Q16595WT1 | | 0 | 28.16 | 1.90E-02 | 2 | 925.45 | -0.53 | 7.33 | 1 |

| WT_1h_R2 | | | | | | | | | | | | | | |
|---|---|---|---|---|---|---|---|---|---|---|---|---|---|---|
| Accession | Description | Score | Coverage | # Proteins | # Unique Peptides | # Peptides | # PSMs | # AAs | MW [kDa] | calc. pI | | | | |
| Q16595WT10 | FrataxinWT10 OS=Homo sapiens OX=9606 GN=FXN PE=1 SV=2 - [FRDAWT10_HUMAN] | 11130.33 | 88.98 | 2 | 3 | 18 | 316 | 127 | 14.23 | 5.67 | | | | |
| A3 | Sequence | # PSMs | # Proteins | # Protein Groups | Protein Group Accessions | Modifications | ΔCn | IonScore | Exp Value | Charge | MH+ [Da] | ΔM [ppm] | RT [min] | # Missed Cleavages |
| High | NWVYSHDGVSLHELLAAELTK | 12 | 2 | 1 | Q16595WT10 | | 0 | 151.80 | 1.84E-14 | 3 | 2382.21 | -0.35 | 14.93 | 0 |
| High | TKLDLSSLAYSGKDA | 44 | 3 | 2 | Q16595WT10<br>Q16595WT1 | | 0 | 127.04 | 4.72E-12 | 2 | 1568.82 | -0.71 | 11.48 | 2 |
| High | LAEETLDSLAEFFEDLADKPYTFEDYDVSFGSGVLTVK | 2 | 3 | 2 | Q16595WT10<br>Q16595WT1 | | 0 | 115.94 | 3.85E-11 | 3 | 4261.03 | 0.01 | 18.30 | 0 |
| High | TKLDLSSLAYSGK | 21 | 3 | 2 | Q16595WT10<br>Q16595WT1 | | 0 | 101.04 | 1.55E-09 | 2 | 1382.75 | 1.43 | 11.37 | 1 |
| High | LGGDLGTYVINK | 69 | 4 | 3 | Q16595WT10<br>Q16595WT1<br>Q16595D10 | | 0 | 100.45 | 2.11E-09 | 2 | 1249.67 | -6.57 | 11.53 | 0 |
| High | LDLSSLAYSGK | 11 | 3 | 2 | Q16595WT10<br>Q16595WT1 | | 0 | 91.80 | 1.27E-08 | 2 | 1153.61 | -2.26 | 12.53 | 0 |
| High | LDLSSLAYSGKDA | 31 | 3 | 2 | Q16595WT10<br>Q16595WT1 | | 0 | 80.18 | 1.77E-07 | 2 | 1339.67 | -1.04 | 12.70 | 1 |
| High | ALKTKLDLSSLAYSGKDA | 6 | 3 | 2 | Q16595WT10<br>Q16595WT1 | | 0 | 80.00 | 1.95E-07 | 3 | 1881.03 | -1.19 | 11.33 | 3 |
| High | YDWTGKNWVYSHDGVSLHELLAAELTK | 2 | 2 | 1 | Q16595WT10 | | 0 | 71.19 | 1.90E-06 | 4 | 3132.54 | -2.41 | 15.32 | 1 |
| High | QIWLSSPSSGPK | 58 | 3 | 2 | Q16595WT10<br>Q16595WT1 | | 0 | 69.51 | 2.93E-06 | 2 | 1286.67 | -2.44 | 10.88 | 0 |
| High | LGGDLGTYVINKQTPNKQIWLSSPSSGPKR | 4 | 3 | 2 | Q16595WT10<br>Q16595WT1 | | 0 | 69.00 | 2.08E-06 | 5 | 3241.72 | -4.76 | 11.78 | 3 |
| High | QTPNKQIWLSSPSSGPK | 14 | 3 | 2 | Q16595WT10<br>Q16595WT1 | | 0 | 65.60 | 6.14E-06 | 3 | 1854.97 | -0.44 | 9.78 | 1 |
| High | QTPNKQIWLSSPSSGPKR | 12 | 3 | 2 | Q16595WT10<br>Q16595WT1 | | 0 | 60.75 | 1.84E-05 | 3 | 2011.07 | -2.45 | 8.87 | 2 |
| High | QIWLSSPSSGPKR | 22 | 3 | 2 | Q16595WT10<br>Q16595WT1 | | 0 | 59.44 | 2.71E-05 | 2 | 1442.78 | 1.82 | 10.52 | 1 |
| High | RYDWTGKNWVYSHDGVSLHELLAAELTK | 1 | 2 | 1 | Q16595WT10 | | 0 | 51.41 | 1.80E-04 | 5 | 3288.63 | -4.73 | 14.70 | 2 |
| High | LGGDLGTYVINKQTPNKQIWLSSPSSGPK | 3 | 3 | 2 | Q16595WT10<br>Q16595WT1 | | 0 | 32.56 | 9.96E-03 | 4 | 3085.62 | -2.94 | 12.40 | 2 |
| High | RYDWTGK | 2 | 3 | 2 | Q16595WT10<br>Q16595WT1 | | 0 | 29.82 | 1.33E-02 | 2 | 925.45 | -6.51 | 7.30 | 1 |
| High | LGGDLGTYVINKQTPNK | 2 | 4 | 3 | Q16595WT10<br>Q16595WT1<br>Q16595D10 | | 0 | 26.75 | 5.13E-02 | 3 | 1817.97 | -3.61 | 10.57 | 1 |

| WT_1h_R3 | | | | | | | | | | | | | | |
|---|---|---|---|---|---|---|---|---|---|---|---|---|---|---|
| Accession | Description | Score | Coverage | # Proteins | # Unique Peptides | # Peptides | # PSMs | # AAs | MW [kDa] | calc. pI | | | | |
| Q16595WT10 | FrataxinWT10 OS=Homo sapiens OX=9606 GN=FXN PE=1 SV=2 - [FRDAWT10_HUMAN] | 9459.54 | 88.98 | 2 | 3 | 18 | 272 | 127 | 14.23 | 5.67 | | | | |
| A3 | Sequence | # PSMs | # Proteins | # Protein Groups | Protein Group Accessions | Modifications | ΔCn | IonScore | Exp Value | Charge | MH+ [Da] | ΔM [ppm] | RT [min] | # Missed Cleavages |
| High | NWVYSHDGVSLHELLAAELTK | 12 | 2 | 1 | Q16595WT10 | | 0 | 141.73 | 1.87E-13 | 3 | 2382.21 | 0.22 | 14.92 | 0 |
| High | TKLDLSSLAYSGKDA | 38 | 3 | 2 | Q16595WT10 Q16595WT1 | | 0 | 140.98 | 1.82E-13 | 2 | 1568.82 | 1.24 | 11.50 | 2 |
| High | LGGDLGTYVINK | 68 | 4 | 3 | Q16595WT10 Q16595WT1 Q16595D10 | | 0 | 102.24 | 1.28E-09 | 2 | 1249.68 | -2.21 | 11.55 | 0 |
| High | TKLDLSSLAYSGK | 9 | 3 | 2 | Q16595WT10 Q16595WT1 | | 0 | 93.49 | 8.62E-09 | 2 | 1382.75 | -1.62 | 11.30 | 1 |
| High | LDLSSLAYSGK | 14 | 3 | 2 | Q16595WT10 Q16595WT1 | | 0 | 85.27 | 5.22E-08 | 2 | 1153.61 | -1.10 | 12.58 | 0 |
| High | LDLSSLAYSGKDA | 26 | 3 | 2 | Q16595WT10 Q16595WT1 | | 0 | 79.51 | 2.09E-07 | 2 | 1339.67 | 0.08 | 13.07 | 1 |
| High | ALKTKLDLSSLAYSGKDA | 6 | 3 | 2 | Q16595WT10 Q16595WT1 | | 0 | 76.64 | 4.04E-07 | 3 | 1881.03 | -0.33 | 11.30 | 3 |
| High | QTPNKQIWLSSPSSGPKR | 7 | 3 | 2 | Q16595WT10 Q16595WT1 | | 0 | 67.05 | 4.21E-06 | 3 | 2011.08 | 4.34 | 8.82 | 2 |
| High | QIWLSSPSSGPK | 49 | 3 | 2 | Q16595WT10 Q16595WT1 | | 0 | 63.87 | 1.05E-05 | 2 | 1286.67 | -1.44 | 11.67 | 0 |
| High | YDWTGKNWVYSHDGVSLHELLAAELTK | 2 | 2 | 1 | Q16595WT10 | | 0 | 61.94 | 1.50E-05 | 4 | 3132.53 | -4.27 | 15.30 | 1 |
| High | LGGDLGTYVINKQTPNKQIWLSSPSSGPKR | 2 | 3 | 2 | Q16595WT10 Q16595WT1 | | 0 | 60.42 | 1.39E-05 | 4 | 3241.73 | -0.61 | 11.77 | 3 |
| High | LAEETLDSLAEFFEDLADKPYTFEDYDVSFGSGVLTVK | 3 | 3 | 2 | Q16595WT10 Q16595WT1 | | 0 | 58.09 | 2.10E-05 | 4 | 4261.01 | -4.59 | 18.30 | 0 |
| High | LGGDLGTYVINKQTPNKQIWLSSPSSGPK | 3 | 3 | 2 | Q16595WT10 Q16595WT1 | | 0 | 55.63 | 5.39E-05 | 4 | 3085.60 | -8.75 | 12.42 | 2 |
| High | QTPNKQIWLSSPSSGPK | 12 | 3 | 2 | Q16595WT10 Q16595WT1 | | 0 | 53.79 | 9.30E-05 | 2 | 1854.97 | -3.12 | 9.77 | 1 |
| High | QIWLSSPSSGPKR | 9 | 3 | 2 | Q16595WT10 Q16595WT1 | | 0 | 52.72 | 1.32E-04 | 2 | 1442.77 | -0.73 | 9.68 | 1 |
| High | RYDWTGKNWVYSHDGVSLHELLAAELTK | 3 | 2 | 1 | Q16595WT10 | | 0 | 51.77 | 1.62E-04 | 5 | 3288.64 | 0.23 | 14.67 | 2 |
| High | LGGDLGTYVINKQTPNK | 3 | 4 | 3 | Q16595WT10 Q16595WT1 Q16595D10 | | 0 | 40.90 | 2.02E-03 | 2 | 1817.96 | -6.67 | 10.52 | 1 |
| High | RYDWTGK | 6 | 3 | 2 | Q16595WT10 Q16595WT1 | | 0 | 29.62 | 1.35E-02 | 2 | 925.45 | -3.32 | 7.32 | 1 |

| FXN-10_5min_R1 | | | | | | | | | | | | | | |
|---|---|---|---|---|---|---|---|---|---|---|---|---|---|---|
| Accession | Description | Score | Coverage | # Proteins | # Unique Peptides | # Peptides | # PSMs | # AAs | MW [kDa] | calc. pI | | | | |
| Q16595D10 | FrataxinD10 OS=Homo sapiens OX=9606 GN=FXN PE=1 SV=2 - [FRDAD10_HUMAN] | 1193.81 | 51.97 | 1 | 8 | 9 | 37 | 127 | 14.41 | 5.87 | | | | |
| A3 | Sequence | # PSMs | # Proteins | # Protein Groups | Protein Group Accessions | Modifications | ΔCn | IonScore | Exp Value | Charge | MH+ [Da] | ΔM [ppm] | RT [min] | # Missed Cleavages |
| High | LGGDLGTYVINK | 4 | 4 | 3 | Q16595WT10 Q16595WT1 Q16595D10 | | 0 | 75.76 | 5.81E-07 | 2 | 1249.66 | -11.71 | 11.50 | 0 |
| High | YDWTGTNWVYSHDGK | 3 | 1 | 1 | Q16595D10 | | 0 | 64.01 | 7.92E-06 | 2 | 1828.78 | -9.28 | 12.15 | 0 |
| High | QIWLSSPTSGPK | 9 | 1 | 1 | Q16595D10 | | 0 | 60.34 | 2.19E-05 | 2 | 1300.68 | -5.45 | 11.27 | 0 |
| High | TKLDLSHLK | 3 | 1 | 1 | Q16595D10 | | 0 | 58.75 | 2.66E-05 | 2 | 1054.62 | -6.73 | 8.77 | 1 |
| High | SLHELLSEELSK | 4 | 1 | 1 | Q16595D10 | | 0 | 53.82 | 8.38E-05 | 2 | 1384.73 | -4.43 | 12.13 | 0 |
| High | RYDWTGTNWVYSHDGK | 3 | 1 | 1 | Q16595D10 | | 0 | 53.14 | 9.68E-05 | 3 | 1984.88 | -5.67 | 11.03 | 1 |
| High | QTPNKQIWLSSPTSGPK | 5 | 1 | 1 | Q16595D10 | | 0 | 48.69 | 3.16E-04 | 3 | 1868.97 | -10.33 | 10.08 | 1 |
| High | QIWLSSPTSGPKR | 3 | 1 | 1 | Q16595D10 | | 0 | 37.59 | 3.99E-03 | 2 | 1456.78 | -5.84 | 9.98 | 1 |
| High | QTPNKQIWLSSPTSGPKR | 3 | 1 | 1 | Q16595D10 | | 0 | 35.65 | 6.28E-03 | 3 | 2025.09 | -0.39 | 9.13 | 2 |

| FXN-10_5min_R2 | | | | | | | | | | | | | | |
|---|---|---|---|---|---|---|---|---|---|---|---|---|---|---|
| Accession | Description | Score | Coverage | # Proteins | # Unique Peptides | # Peptides | # PSMs | # AAs | MW [kDa] | calc. pI | | | | |
| Q16595WT1 | FrataxinWT1 OS=Homo sapiens OX=9606 GN=FXN PE=1 SV=2 - [FRDAWT1_HUMAN] | 870.50 | 54.33 | 1 | 7 | 8 | 24 | 127 | 14.28 | 5.47 | | | | |
| A3 | Sequence | # PSMs | # Proteins | # Protein Groups | Protein Group Accessions | Modifications | ΔCn | IonScore | Exp Value | Charge | MH+ [Da] | ΔM [ppm] | RT [min] | # Missed Cleavages |
| High | LGGDLGTYVINK | 4 | 2 | 2 | Q16595WT1 Q16595D10 | | 0 | 100.11 | 2.29E-09 | 2 | 1249.67 | -6.70 | 11.53 | 0 |
| High | TKLDLSSLAYSGKDA | 2 | 1 | 1 | Q16595WT1 | | 0 | 74.91 | 7.34E-07 | 2 | 1568.81 | -6.19 | 11.52 | 2 |
| High | LDLSSLAYSGKDA | 3 | 1 | 1 | Q16595WT1 | | 0 | 73.53 | 8.85E-07 | 2 | 1339.67 | -1.61 | 12.68 | 1 |
| High | ALKTKLDLSSLAYSGKDA | 5 | 1 | 1 | Q16595WT1 | | 0 | 63.41 | 1.00E-05 | 3 | 1881.02 | -7.76 | 11.35 | 3 |
| High | QTPNKQIWLSSPSSGPKR | 4 | 1 | 1 | Q16595WT1 | | 0 | 56.49 | 4.95E-05 | 3 | 2011.06 | -3.51 | 8.88 | 2 |
| High | QIWLSSPSSGPK | 4 | 1 | 1 | Q16595WT1 | | 0 | 41.93 | 1.53E-03 | 2 | 1286.67 | -6.64 | 10.92 | 0 |
| High | NWVYSHDGVSLHELLADELTK | 1 | 1 | 1 | Q16595WT1 | | 0 | 26.32 | 5.13E-02 | 3 | 2426.18 | -6.30 | 14.77 | 0 |
| High | LGGDLGTYVINKQTPNKQIWLSSPSSGPKR | 1 | 1 | 1 | Q16595WT1 | | 0 | 20.60 | 1.74E-01 | 4 | 3241.69 | -14.19 | 11.82 | 3 |

| FXN-10_5min_R3 | | | | | | | | | | | | | | |
|---|---|---|---|---|---|---|---|---|---|---|---|---|---|---|
| Accession | Description | Score | Coverage | # Proteins | # Unique Peptides | # Peptides | # PSMs | # AAs | MW [kDa] | calc. pI | | | | |
| Q16595D10 | FrataxinD10 OS=Homo sapiens OX=9606 GN=FXN PE=1 SV=2 - [FRDAD10_HUMAN] | 1213.65 | 56.69 | 1 | 9 | 10 | 34 | 127 | 14.41 | 5.87 | | | | |
| A3 | Sequence | # PSMs | # Proteins | # Protein Groups | Protein Group Accessions | Modifications | ΔCn | IonScore | Exp Value | Charge | MH+ [Da] | ΔM [ppm] | RT [min] | # Missed Cleavages |
| High | LGGDLGTYVINK | 6 | 4 | 3 | Q16595WT10 Q16595WT1 Q16595D10 | | 0 | 93.28 | 1.09E-08 | 2 | 1249.67 | -7.77 | 11.50 | 0 |
| High | QIWLSSPTSGPK | 11 | 1 | 1 | Q16595D10 | | 0 | 75.89 | 6.02E-07 | 2 | 1300.68 | -4.60 | 11.23 | 0 |
| High | RYDWTGTNWVYSHDGK | 4 | 1 | 1 | Q16595D10 | | 0 | 70.95 | 1.05E-06 | 3 | 1984.88 | -7.64 | 11.00 | 1 |
| High | SLHELLSEELSK | 2 | 1 | 1 | Q16595D10 | | 0 | 62.86 | 1.06E-05 | 2 | 1384.72 | -7.37 | 12.10 | 0 |
| High | QTPNKQIWLSSPTSGPK | 3 | 1 | 1 | Q16595D10 | | 0 | 53.76 | 9.80E-05 | 3 | 1868.97 | -8.41 | 10.05 | 1 |
| High | YDWTGTNWVYSHDGK | 1 | 1 | 1 | Q16595D10 | | 0 | 46.69 | 2.81E-04 | 2 | 1828.78 | -5.53 | 12.12 | 0 |
| High | TKLDLSHLK | 1 | 1 | 1 | Q16595D10 | | 0 | 44.41 | 4.75E-04 | 2 | 1054.62 | -5.88 | 8.78 | 1 |
| High | QIWLSSPTSGPKR | 2 | 1 | 1 | Q16595D10 | | 0 | 38.40 | 3.33E-03 | 2 | 1456.78 | -9.22 | 9.97 | 1 |
| High | QTPNKQIWLSSPTSGPKR | 3 | 1 | 1 | Q16595D10 | | 0 | 37.27 | 4.40E-03 | 3 | 2025.07 | -7.96 | 9.12 | 2 |
| High | LDLSHLKYSGKDA | 1 | 1 | 1 | Q16595D10 | | 0 | 24.52 | 1.01E-01 | 2 | 1446.75 | -4.50 | 9.88 | 2 |

| FXN-10_10min_R1 | | | | | | | | | | | | | | |
|---|---|---|---|---|---|---|---|---|---|---|---|---|---|---|
| Accession | Description | Score | Coverage | # Proteins | # Unique Peptides | # Peptides | # PSMs | # AAs | MW [kDa] | calc. pI | | | | |
| Q16595D10 | FrataxinD10 OS=Homo sapiens OX=9606 GN=FXN PE=1 SV=2 - [FRDAD10_HUMAN] | 1346.57 | 51.97 | 1 | 8 | 9 | 37 | 127 | 14.41 | 5.87 | | | | |
| A3 | Sequence | # PSMs | # Proteins | # Protein Groups | Protein Group Accessions | Modifications | ΔCn | IonScore | Exp Value | Charge | MH+ [Da] | ΔM [ppm] | RT [min] | # Missed Cleavages |
| High | YDWTGTNWVYSHDGK | 2 | 1 | 1 | Q16595D10 | | 0 | 89.26 | 9.37E-09 | 2 | 1828.77 | -11.30 | 12.17 | 0 |
| High | LGGDLGTYVINK | 7 | 4 | 2 | Q16595WT1 Q16595D10 | | 0 | 82.88 | 1.15E-07 | 2 | 1249.67 | -8.38 | 11.57 | 0 |
| High | QIWLSSPTSGPK | 8 | 1 | 1 | Q16595D10 | | 0 | 69.60 | 2.52E-06 | 2 | 1300.68 | -10.28 | 11.22 | 0 |
| High | QTPNKQIWLSSPTSGPK | 5 | 1 | 1 | Q16595D10 | | 0 | 63.34 | 1.06E-05 | 3 | 1868.97 | -6.26 | 10.10 | 1 |
| High | RYDWTGTNWVYSHDGK | 4 | 1 | 1 | Q16595D10 | | 0 | 60.74 | 6.66E-06 | 3 | 1984.86 | -15.83 | 11.05 | 1 |
| High | TKLDLSHLK | 5 | 1 | 1 | Q16595D10 | | 0 | 58.99 | 1.34E-05 | 2 | 1054.62 | -6.05 | 8.77 | 1 |
| High | SLHELLSEELSK | 2 | 1 | 1 | Q16595D10 | | 0 | 49.82 | 2.14E-04 | 2 | 1384.72 | -10.79 | 12.15 | 0 |
| High | QIWLSSPTSGPKR | 3 | 1 | 1 | Q16595D10 | | 0 | 40.46 | 2.07E-03 | 2 | 1456.78 | -9.27 | 10.00 | 1 |
| High | QTPNKQIWLSSPTSGPKR | 1 | 1 | 1 | Q16595D10 | | 0 | 30.75 | 2.11E-02 | 3 | 2025.05 | -16.70 | 9.15 | 2 |

| FXN-10_10min_R2 | | | | | | | | | | | | | | |
|---|---|---|---|---|---|---|---|---|---|---|---|---|---|---|
| Accession | Description | Score | Coverage | # Proteins | # Unique Peptides | # Peptides | # PSMs | # AAs | MW [kDa] | calc. pI | | | | |
| Q16595D10 | FrataxinD10 OS=Homo sapiens OX=9606 GN=FXN PE=1 SV=2 - [FRDAD10_HUMAN] | 1469.35 | 56.69 | 1 | 9 | 10 | 42 | 127 | 14.41 | 5.87 | | | | |
| A3 | Sequence | # PSMs | # Proteins | # Protein Groups | Protein Group Accessions | Modifications | ΔCn | IonScore | Exp Value | Charge | MH+ [Da] | ΔM [ppm] | RT [min] | # Missed Cleavages |
| High | LGGDLGTYVINK | 6 | 2 | 2 | Q16595WT1 Q16595D10 | | 0 | 102.97 | 1.10E-09 | 2 | 1249.67 | -6.31 | 11.58 | 0 |
| High | RYDWTGTNWVYSHDGK | 9 | 1 | 1 | Q16595D10 | | 0 | 68.35 | 1.43E-06 | 3 | 1984.87 | -10.20 | 11.03 | 1 |
| High | YDWTGTNWVYSHDGK | 2 | 1 | 1 | Q16595D10 | | 0 | 61.77 | 6.51E-06 | 2 | 1828.78 | -8.89 | 12.12 | 0 |
| High | QIWLSSPTSGPK | 8 | 1 | 1 | Q16595D10 | | 0 | 60.79 | 1.92E-05 | 2 | 1300.68 | -11.09 | 11.25 | 0 |
| High | QTPNKQIWLSSPTSGPK | 5 | 1 | 1 | Q16595D10 | | 0 | 59.80 | 2.46E-05 | 3 | 1868.97 | -10.48 | 10.07 | 1 |
| High | SLHELLSEELSK | 3 | 1 | 1 | Q16595D10 | | 0 | 54.72 | 6.91E-05 | 2 | 1384.72 | -8.67 | 12.15 | 0 |
| High | LDLSHLKYSGKDA | 3 | 1 | 1 | Q16595D10 | | 0 | 51.95 | 1.75E-04 | 2 | 1446.75 | -8.30 | 9.88 | 2 |
| High | QIWLSSPTSGPKR | 3 | 1 | 1 | Q16595D10 | | 0 | 50.25 | 2.17E-04 | 2 | 1456.78 | -8.65 | 9.98 | 1 |
| High | QTPNKQIWLSSPTSGPKR | 2 | 1 | 1 | Q16595D10 | | 0 | 48.79 | 3.12E-04 | 3 | 2025.08 | -3.30 | 9.12 | 2 |
| High | TKLDLSHLK | 1 | 1 | 1 | Q16595D10 | | 0 | 45.24 | 2.93E-04 | 2 | 1054.62 | -9.40 | 8.78 | 1 |

| FXN-10_10min_R3 | | | | | | | | | | | | | | |
|---|---|---|---|---|---|---|---|---|---|---|---|---|---|---|
| Accession | Description | Score | Coverage | # Proteins | # Unique Peptides | # Peptides | # PSMs | # AAs | MW [kDa] | calc. pI | | | | |
| Q16595D10 | FrataxinD10 OS=Homo sapiens OX=9606 GN=FXN PE=1 SV=2 - [FRDAD10_HUMAN] | 1239.09 | 51.97 | 1 | 8 | 9 | 34 | 127 | 14.41 | 5.87 | | | | |
| A3 | Sequence | # PSMs | # Proteins | # Protein Groups | Protein Group Accessions | Modifications | ΔCn | IonScore | Exp Value | Charge | MH+ [Da] | ΔM [ppm] | RT [min] | # Missed Cleavages |
| High | LGGDLGTYVINK | 6 | 4 | 2 | Q16595WT1 Q16595D10 | | 0 | 87.83 | 3.58E-08 | 2 | 1249.67 | -5.74 | 11.55 | 0 |
| High | YDWTGTNWVYSHDGK | 3 | 1 | 1 | Q16595D10 | | 0 | 68.75 | 1.55E-06 | 2 | 1828.78 | -6.16 | 12.15 | 0 |
| High | QIWLSSPTSGPK | 6 | 1 | 1 | Q16595D10 | | 0 | 66.25 | 5.66E-06 | 2 | 1300.68 | -11.22 | 11.22 | 0 |
| High | RYDWTGTNWVYSHDGK | 3 | 1 | 1 | Q16595D10 | | 0 | 63.34 | 5.37E-06 | 3 | 1984.87 | -11.34 | 11.03 | 1 |
| High | SLHELLSEELSK | 3 | 1 | 1 | Q16595D10 | | 0 | 58.26 | 3.06E-05 | 2 | 1384.72 | -10.27 | 12.17 | 0 |
| High | QTPNKQIWLSSPTSGPK | 5 | 1 | 1 | Q16595D10 | | 0 | 57.08 | 4.56E-05 | 3 | 1868.97 | -8.42 | 10.07 | 1 |
| High | QIWLSSPTSGPKR | 3 | 1 | 1 | Q16595D10 | | 0 | 55.07 | 7.20E-05 | 2 | 1456.78 | -9.66 | 9.98 | 1 |
| High | QTPNKQIWLSSPTSGPKR | 1 | 1 | 1 | Q16595D10 | | 0 | 46.84 | 4.79E-04 | 3 | 2025.07 | -9.71 | 9.15 | 2 |
| High | TKLDLSHLK | 4 | 1 | 1 | Q16595D10 | | 0 | 41.76 | 7.73E-04 | 2 | 1054.62 | -9.67 | 8.82 | 1 |

| FXN-10_20min_R1 | | | | | | | | | | | | | | |
|---|---|---|---|---|---|---|---|---|---|---|---|---|---|---|
| Accession | Description | Score | Coverage | # Proteins | # Unique Peptides | # Peptides | # PSMs | # AAs | MW [kDa] | calc. pI | | | | |
| Q16595D10 | FrataxinD10 OS=Homo sapiens OX=9606 GN=FXN PE=1 SV=2 - [FRDAD10_HUMAN] | 1714.96 | 56.69 | 1 | 9 | 10 | 54 | 127 | 14.41 | 5.87 | | | | |
| A3 | Sequence | # PSMs | # Proteins | # Protein Groups | Protein Group Accessions | Modifications | ΔCn | IonScore | Exp Value | Charge | MH+ [Da] | ΔM [ppm] | RT [min] | # Missed Cleavages |
| High | LGGDLGTYVINK | 12 | 4 | 3 | Q16595WT10 Q16595WT1 Q16595D10 | | 0 | 91.23 | 1.73E-08 | 2 | 1249.68 | -0.34 | 11.55 | 0 |
| High | RYDWTGTNWVYSHDGK | 6 | 1 | 1 | Q16595D10 | | 0 | 84.57 | 3.51E-08 | 3 | 1984.89 | -3.58 | 11.03 | 1 |
| High | YDWTGTNWVYSHDGK | 3 | 1 | 1 | Q16595D10 | | 0 | 81.11 | 7.79E-08 | 2 | 1828.78 | -5.31 | 12.13 | 0 |
| High | LDLSHLKYSGKDA | 2 | 1 | 1 | Q16595D10 | | 0 | 75.28 | 8.26E-07 | 2 | 1446.75 | -4.58 | 9.90 | 2 |
| High | SLHELLSEELSK | 3 | 1 | 1 | Q16595D10 | | 0 | 71.67 | 1.49E-06 | 2 | 1384.73 | -2.66 | 12.15 | 0 |
| High | QIWLSSPTSGPK | 12 | 1 | 1 | Q16595D10 | | 0 | 69.74 | 2.48E-06 | 2 | 1300.68 | -5.17 | 11.18 | 0 |
| High | TKLDLSHLK | 3 | 1 | 1 | Q16595D10 | | 0 | 55.83 | 2.80E-05 | 2 | 1054.62 | -6.62 | 8.77 | 1 |
| High | QTPNKQIWLSSPTSGPKR | 2 | 1 | 1 | Q16595D10 | | 0 | 51.48 | 1.67E-04 | 3 | 2025.08 | -3.50 | 9.15 | 2 |
| High | QIWLSSPTSGPKR | 4 | 1 | 1 | Q16595D10 | | 0 | 40.59 | 2.06E-03 | 2 | 1456.79 | -0.79 | 10.00 | 1 |
| High | QTPNKQIWLSSPTSGPK | 7 | 1 | 1 | Q16595D10 | | 0 | 40.49 | 1.96E-03 | 3 | 1868.98 | -2.05 | 10.08 | 1 |

| FXN-10_20min_R2 | | | | | | | | | | | | | | |
|---|---|---|---|---|---|---|---|---|---|---|---|---|---|---|
| Accession | Description | Score | Coverage | # Proteins | # Unique Peptides | # Peptides | # PSMs | # AAs | MW [kDa] | calc. pI | | | | |
| Q16595D10 | FrataxinD10 OS=Homo sapiens OX=9606 GN=FXN PE=1 SV=2 - [FRDAD10_HUMAN] | 1653.96 | 56.69 | 1 | 9 | 10 | 50 | 127 | 14.41 | 5.87 | | | | |
| A3 | Sequence | # PSMs | # Proteins | # Protein Groups | Protein Group Accessions | Modifications | ΔCn | IonScore | Exp Value | Charge | MH+ [Da] | ΔM [ppm] | RT [min] | # Missed Cleavages |
| High | SLHELLSEELSK | 6 | 1 | 1 | Q16595D10 | | 0 | 91.48 | 1.56E-08 | 2 | 1384.73 | -2.81 | 12.18 | 0 |
| High | LGGDLGTYVINK | 5 | 4 | 3 | Q16595WT10 Q16595WT1 Q16595D10 | | 0 | 85.74 | 5.79E-08 | 2 | 1249.67 | -3.34 | 11.52 | 0 |
| High | YDWTGTNWVYSHDGK | 2 | 1 | 1 | Q16595D10 | | 0 | 84.32 | 4.45E-08 | 2 | 1828.78 | -5.16 | 12.12 | 0 |
| High | TKLDLSHLK | 3 | 1 | 1 | Q16595D10 | | 0 | 78.37 | 1.75E-07 | 2 | 1054.62 | -4.30 | 8.77 | 1 |
| High | QIWLSSPTSGPK | 9 | 1 | 1 | Q16595D10 | | 0 | 57.71 | 4.20E-05 | 2 | 1300.69 | -2.45 | 11.18 | 0 |
| High | LDLSHLKYSGKDA | 2 | 1 | 1 | Q16595D10 | | 0 | 54.01 | 1.09E-04 | 2 | 1446.75 | -5.56 | 9.85 | 2 |
| High | RYDWTGTNWVYSHDGK | 9 | 1 | 1 | Q16595D10 | | 0 | 53.86 | 4.95E-05 | 3 | 1984.89 | -1.80 | 10.98 | 1 |
| High | QTPNKQIWLSSPTSGPK | 8 | 1 | 1 | Q16595D10 | | 0 | 48.52 | 3.08E-04 | 3 | 1868.98 | -2.52 | 10.02 | 1 |
| High | QIWLSSPTSGPKR | 4 | 1 | 1 | Q16595D10 | | 0 | 44.11 | 9.30E-04 | 2 | 1456.79 | -0.47 | 9.95 | 1 |
| High | QTPNKQIWLSSPTSGPKR | 2 | 1 | 1 | Q16595D10 | | 0 | 43.39 | 1.07E-03 | 3 | 2025.08 | -2.87 | 9.10 | 2 |

| FXN-10_20min_R3 | | | | | | | | | | | | | | |
|---|---|---|---|---|---|---|---|---|---|---|---|---|---|---|
| Accession | Description | Score | Coverage | # Proteins | # Unique Peptides | # Peptides | # PSMs | # AAs | MW [kDa] | calc. pI | | | | |
| Q16595D10 | FrataxinD10 OS=Homo sapiens OX=9606 GN=FXN PE=1 SV=2 - [FRDAD10_HUMAN] | 1767.99 | 56.69 | 1 | 9 | 10 | 49 | 127 | 14.41 | 5.87 | | | | |
| A3 | Sequence | # PSMs | # Proteins | # Protein Groups | Protein Group Accessions | Modifications | ΔCn | IonScore | Exp Value | Charge | MH+ [Da] | ΔM [ppm] | RT [min] | # Missed Cleavages |
| High | SLHELLSEELSK | 5 | 1 | 1 | Q16595D10 | | 0 | 94.92 | 7.34E-09 | 2 | 1384.73 | -0.61 | 12.18 | 0 |
| High | LGGDLGTYVINK | 7 | 4 | 2 | Q16595WT1 Q16595D10 | | 0 | 88.32 | 3.08E-08 | 2 | 1249.68 | 0.01 | 11.53 | 0 |
| High | RYDWTGTNWVYSHDGK | 8 | 1 | 1 | Q16595D10 | | 0 | 73.04 | 5.88E-07 | 3 | 1984.89 | -4.01 | 11.03 | 1 |
| High | QTPNKQIWLSSPTSGPK | 7 | 1 | 1 | Q16595D10 | | 0 | 64.54 | 8.05E-06 | 3 | 1868.97 | -6.40 | 10.07 | 1 |
| High | YDWTGTNWVYSHDGK | 3 | 1 | 1 | Q16595D10 | | 0 | 57.40 | 2.15E-05 | 2 | 1828.79 | 0.73 | 12.13 | 0 |
| High | QIWLSSPTSGPK | 9 | 1 | 1 | Q16595D10 | | 0 | 57.05 | 4.79E-05 | 2 | 1300.69 | -3.44 | 11.22 | 0 |
| High | QTPNKQIWLSSPTSGPKR | 2 | 1 | 1 | Q16595D10 | | 0 | 56.23 | 5.77E-05 | 3 | 2025.08 | -3.89 | 9.10 | 2 |
| High | LDLSHLKYSGKDA | 2 | 1 | 1 | Q16595D10 | | 0 | 49.35 | 3.23E-04 | 2 | 1446.75 | -7.11 | 9.90 | 2 |
| High | TKLDLSHLK | 3 | 1 | 1 | Q16595D10 | | 0 | 45.62 | 3.24E-04 | 2 | 1054.62 | -3.64 | 8.75 | 1 |
| High | QIWLSSPTSGPKR | 3 | 1 | 1 | Q16595D10 | | 0 | 40.22 | 2.33E-03 | 2 | 1456.78 | -5.19 | 9.97 | 1 |

| FXN-10_1h_R1 | | | | | | | | | | | | | | |
|---|---|---|---|---|---|---|---|---|---|---|---|---|---|---|
| Accession | Description | Score | Coverage | # Proteins | # Unique Peptides | # Peptides | # PSMs | # AAs | MW [kDa] | calc. pI | | | | |
| Q16595D10 | FrataxinD10 OS=Homo sapiens OX=9606 GN=FXN PE=1 SV=2 - [FRDAD10_HUMAN] | 2647.30 | 77.95 | 1 | 6 | 7 | 90 | 127 | 14.41 | 5.87 | | | | |
| A3 | Sequence | # PSMs | # Proteins | # Protein Groups | Protein Group Accessions | Modifications | ΔCn | IonScore | Exp Value | Charge | MH+ [Da] | ΔM [ppm] | RT [min] | # Missed Cleavages |
| High | LGGDLGTYVINK | 68 | 4 | 3 | Q16595WT10 Q16595WT1 Q16595D10 | | 0.00 | 97.28 | 4.28E-09 | 2 | 1249.68 | -1.61 | 11.52 | 0 |
| High | QIWLSSPTSGPK | 10 | 1 | 1 | Q16595D10 | | 0.00 | 65.15 | 7.15E-06 | 2 | 1300.68 | -5.82 | 11.25 | 0 |
| High | SLHELLSEELSK | 4 | 1 | 1 | Q16595D10 | | 0.00 | 59.54 | 2.27E-05 | 2 | 1384.72 | -7.62 | 12.17 | 0 |
| High | TKLDLSHLK | 2 | 1 | 1 | Q16595D10 | | 0.00 | 54.54 | 7.01E-05 | 2 | 1054.62 | -0.78 | 8.75 | 1 |
| High | RYDWTGTNWVYSHDGK | 2 | 1 | 1 | Q16595D10 | | 0.00 | 45.36 | 5.81E-04 | 3 | 1984.88 | -7.42 | 11.02 | 1 |
| High | QIWLSSPTSGPKR | 3 | 1 | 1 | Q16595D10 | | 0.00 | 36.22 | 5.64E-03 | 2 | 1456.79 | -1.09 | 9.97 | 1 |
| High | LAEETLDSLAEFFEDLKDKPFTPEDYDVSFGDGVLTVK | 1 | 1 | 1 | Q16595D10 | | 0.00 | 31.67 | 1.36E-02 | 4 | 4280.05 | -4.87 | 18.27 | 1 |

| FXN-10_1h_R2 | | | | | | | | | | | | | | |
|---|---|---|---|---|---|---|---|---|---|---|---|---|---|---|
| Accession | Description | Score | Coverage | # Proteins | # Unique Peptides | # Peptides | # PSMs | # AAs | MW [kDa] | calc. pI | | | | |
| Q16595D10 | FrataxinD10 OS=Homo sapiens OX=9606 GN=FXN PE=1 SV=2 - [FRDAD10_HUMAN] | 2082.73 | 56.69 | 1 | 9 | 10 | 66 | 127 | 14.41 | 5.87 | | | | |
| A3 | Sequence | # PSMs | # Proteins | # Protein Groups | Protein Group Accessions | Modifications | ΔCn | IonScore | Exp Value | Charge | MH+ [Da] | ΔM [ppm] | RT [min] | # Missed Cleavages |
| High | LGGDLGTYVINK | 19 | 2 | 2 | Q16595WT1 Q16595D10 | | 0 | 96.54 | 5.10E-09 | 2 | 1249.68 | -0.92 | 11.52 | 0 |
| High | SLHELLSEELSK | 7 | 1 | 1 | Q16595D10 | | 0 | 91.76 | 1.52E-08 | 2 | 1384.73 | -0.58 | 12.17 | 0 |
| High | RYDWTGTNWVYSHDGK | 12 | 1 | 1 | Q16595D10 | | 0 | 87.85 | 1.16E-08 | 3 | 1984.89 | -3.66 | 11.03 | 1 |
| High | YDWTGTNWVYSHDGK | 2 | 1 | 1 | Q16595D10 | | 0 | 85.09 | 1.84E-08 | 2 | 1828.78 | -6.27 | 12.08 | 0 |
| High | QIWLSSPTSGPK | 11 | 1 | 1 | Q16595D10 | | 0 | 70.23 | 2.30E-06 | 2 | 1300.68 | -3.57 | 11.20 | 0 |
| High | TKLDLSHLK | 4 | 1 | 1 | Q16595D10 | | 0 | 59.54 | 1.23E-05 | 2 | 1054.62 | -4.49 | 8.75 | 1 |
| High | QTPNKQIWLSSPTSGPK | 4 | 1 | 1 | Q16595D10 | | 0 | 55.29 | 6.29E-05 | 3 | 1868.99 | 0.71 | 10.03 | 1 |
| High | QTPNKQIWLSSPTSGPKR | 2 | 1 | 1 | Q16595D10 | | 0 | 48.13 | 3.61E-04 | 3 | 2025.08 | -4.41 | 9.08 | 2 |
| High | LDLSHLKYSGKDA | 1 | 1 | 1 | Q16595D10 | | 0 | 42.04 | 1.72E-03 | 2 | 1446.75 | -5.96 | 9.85 | 2 |
| High | QIWLSSPTSGPKR | 4 | 1 | 1 | Q16595D10 | | 0 | 39.79 | 2.55E-03 | 2 | 1456.78 | -4.41 | 9.95 | 1 |

| FXN-10_1h_R3 | | | | | | | | | | | | | | |
|---|---|---|---|---|---|---|---|---|---|---|---|---|---|---|
| Accession | Description | Score | Coverage | # Proteins | # Unique Peptides | # Peptides | # PSMs | # AAs | MW [kDa] | calc. pI | | | | |
| Q16595D10 | FrataxinD10 OS=Homo sapiens OX=9606 GN=FXN PE=1 SV=2 - [FRDAD10_HUMAN] | 2251.10 | 56.69 | 1 | 9 | 10 | 67 | 127 | 14.41 | 5.87 | | | | |
| A3 | Sequence | # PSMs | # Proteins | # Protein Groups | Protein Group Accessions | Modifications | ΔCn | IonScore | Exp Value | Charge | MH+ [Da] | ΔM [ppm] | RT [min] | # Missed Cleavages |
| High | LGGDLGTYVINK | 13 | 2 | 2 | Q16595WT1 Q16595D10 | | 0 | 100.31 | 2.13E-09 | 2 | 1249.68 | -1.22 | 11.53 | 0 |
| High | SLHELLSEELSK | 11 | 1 | 1 | Q16595D10 | | 0 | 91.09 | 1.70E-08 | 2 | 1384.73 | -3.27 | 12.13 | 0 |
| High | YDWTGTNWVYSHDGK | 3 | 1 | 1 | Q16595D10 | | 0 | 79.50 | 8.84E-08 | 2 | 1828.79 | -0.34 | 12.10 | 0 |
| High | RYDWTGTNWVYSHDGK | 16 | 1 | 1 | Q16595D10 | | 0 | 76.01 | 1.98E-07 | 3 | 1984.88 | -4.60 | 11.00 | 1 |
| High | QIWLSSPTSGPK | 10 | 1 | 1 | Q16595D10 | | 0 | 74.86 | 7.61E-07 | 2 | 1300.68 | -4.43 | 11.27 | 0 |
| High | LDLSHLKYSGKDA | 2 | 1 | 1 | Q16595D10 | | 0 | 60.98 | 2.39E-05 | 2 | 1446.76 | -1.40 | 9.85 | 2 |
| High | TKLDLSHLK | 3 | 1 | 1 | Q16595D10 | | 0 | 52.34 | 6.48E-05 | 2 | 1054.62 | -5.10 | 8.72 | 1 |
| High | QIWLSSPTSGPKR | 3 | 1 | 1 | Q16595D10 | | 0 | 36.66 | 4.85E-03 | 2 | 1456.78 | -7.57 | 9.93 | 1 |
| High | QTPNKQIWLSSPTSGPK | 4 | 1 | 1 | Q16595D10 | | 0 | 34.89 | 7.43E-03 | 3 | 1868.97 | -6.34 | 10.03 | 1 |
| High | QTPNKQIWLSSPTSGPKR | 2 | 1 | 1 | Q16595D10 | | 0 | 34.23 | 8.55E-03 | 3 | 2025.09 | -1.01 | 9.08 | 2 |

### 13. Binding of frataxin variants to $Zn^{2+}$/ppIX and FeS assembly complex

#### Expression and purification of $^{15}N/^{13}C$ labelled proteins

The recombinant plasmid pG-S21a (purchased from GenScript Biotech) encoding between restriction sites *NdeI* and *XhoI* for residues 91-210 of FXN-10 bearing a N-terminal 6xHis tag, was transformed into BL21 (D3) *E. coli* competent cells, plated on Luria-Bertani (LB) broth-ampicillin agar plates and incubated overnight at 37 ºC. A single colony from each plate was picked and then resuspended in an aqueous solution of 10 mL of LB broth (Lennox); this process was repeated in triplicate. Subsequently, the suspensions were incubated at 37 ºC for 6-8 hours. Next, 200 µL of each of these preinocula were transferred to 200 mL of minimal M9 media (24 mM $Na_2HPO_4$, 11 mM $KH_2PO_4$, 4.3 mM NaCl, 2mM $MgSO_4$, 0.1 mM $CaCl_2$, 200 mg/L thiamine, 10 mg/L ampicillin, 10 mg/L biotine, and 50 mg/L ampicillin) supplemented with trace metal mix composition,[19] along with 1 g/L $^{15}NH_4Cl$ (≥98 atom % $^{15}N$, Sigma-Aldrich) for uniform $^{15}N$ labelling and 2 g/L D-glucose. These cultures were then incubated overnight at 37 ºC. Subsequently, these precultures were added to 1.5 L of minimal M9 media and incubated again at 37 ºC until the OD reached values between 0.4–0.6. Induction was carried out by adding 10 µL of 0.5 M isopropyl β-D-1-thiogalactopyranoside (IPTG) and growth continued overnight at 25 ºC. The subsequent cell harvest and purification steps followed the protocol described in section 6.

Uniform $^{15}N/^{13}C$ labelled human frataxin was prepared following the experimental procedure described previously by us[20] and used as a reference. Briefly, the recombinant plasmid pGS21a (purchased from Genescript) encoding for residues D91 to A210 of wild- type human frataxin fused via a trombin cleavage site to a N-terminal 6xHis-GST tag was transformed into *E. coli* BL21(DE3) competent cells, and expressed under control of the T7 promoter at 30 °C for about 18 h in minimal M9 media (24 mM $Na_2HPO_4$, 11 mM $KH_2PO_4$, 4.3 mM NaCl, 2mM $MgSO_4$, 0.1mM $CaCl_2$, 200 mg/L thiaminehydrochloride, 100 mg/L kanamycin) supplemented with trace metal mix composition,[19] along with 1 g/L $^{15}NH_4Cl$ (≥98 atom % $^{15}N$, Sigma-Aldrich) for uniform $^{15}N$ labelling and 3 g/L D-glucose or, optionally, for the production of samples bearing uniform $^{13}C$ labelling with 2 g/L D-glucose-$^{13}C_6$ (≥99 atom % $^{13}C$, Sigma-Aldrich) as the only nitrogen and carbon source, respectively.

For protein purification, the cell pellet was thawed and resuspended in lysis buffer (20 mM Tris pH 8, 120 mM NaCl, 2 mM imidazole, 10% glycerol, 0.1% triton-X100, and one tablet of cOmplete™ EDTA-free protease inhibitor cocktail), and incubated for about 30 min on ice prior completing cell lysis by ultrasonication on ice for a total time of 2.5 min (Vibracell VC505 sonicator, 14 mm diameter probe). The cell debris was removed by ultracentrifugation at 60k g. The soluble fraction was passed through a 0.22 µm filter and loaded onto 3 mL cobalt-charged NTA agarose column equilibrated with lysis buffer before extensive cleaning with washing buffer (20 mM Tris pH 8, 120 mM NaCl, 2 mM imidazole, and 10 % glycerol) for at least 20 column volumes before elution of the protein from the column in one step in presence of 200 mM imidazole in the washing buffer. For subsequent imidazole removal the pooled fractions were desalted using a Sephadex™ G-25 column equilibrated with 20 mM Tris pH 8, 120 mM NaCl, and 10% glycerol. Subsequently, the 6xHis-GST tag was cleaved by incubation with thrombin at room temperature for 6 h applying 2 units of the enzyme per mg of target protein. Traces for uncleaved protein were removed by a second affinity chromatography step before loading the sample onto a Sd75/16/600 gel filtration column for final polishing. The apparent elution volume was 76.5 mL. The purity of the obtained protein (13.8 kDa) was monitored using SDS-PAGE (5%-20% gradient gel) and the concentration was determined spectrophotometrically (using an extinction coefficient ε of 26930 $cm^{-1}M^{-1}$) and by a Bradford assay.

**Expression and purification of FeS assembly complex**

The FeS assembly complex prepared was composed of the iron-cluster assembly enzyme (ISCU, residues His36 to C-terminal Lys167, 15.0 kDa), human mitochondrial cysteine desulfurase (NFS1, residues Leu56 to C-terminal His457, 47.8 kDa, theoretical pI = 6.7), and human LYR motif-containing protein 4 (LYRM4, residue Arg6 to C-terminal Thr91, 10.6 kDa), all required for the *de novo* synthesis of iron-sulphur (Fe-S) clusters within mitochondria responsible for maturation of both, mitochondrial and cytoplasmic [2Fe-2S] and [4Fe-4S] proteins, respectively. The construct of ISCU was cloned into pET28a vector (purchased from GenScript Biotech), whereas LYRM4 and NFS1 were cloned into pCDFDuet$^{TM}$-1 plasmid (purchased from GenScript Biotech) containing two multiple cloning sites. The primary sequences for all constructs expressed are shown in Figure S15. Co-transformed BL21 *E. coli* cells for in vivo formation of the FeS assembly complex were obtained from sequential chemical transformation. Given the high affinity for the complex formation only the construct for cysteine desulfurase NFS1 was bearing a N-terminal His-tag (MGSSHHHHHH-, followed by a small linker and a TEV cleavage site SQDPNSSSG-ENLYFQ|G-). Thus, during the initial affinity step both other components were captured and co-purified further when bound to that bait protein (providing a secondary affinity support). Under applied buffer conditions (50 mM HEPES Na at pH 8.0, 120 mM NaCl), the hydrodynamic radii of the complex (about 73.4 kDa) observed by gel filtration resembled the size of a di-trimeric complex (as observed in the orthorhombic unit cell of the complex crystal structure (PDB ID 6NZU). The calculated extinction coefficient for the di-trimer complex (2 x 655 residues, 146.2 kDa) is 2 x 57.760 $M^{-1}$ $cm^{-1}$. Lysis of the cells were performed by a freeze and thaw cycle at -80 ºC followed by sonication in the absence of any detergent in the buffer. Due to the basic character of LYR motif-containing protein 4 (LYRM4, pI = 10.7) and the iron-sulphur cluster assembly enzyme (ISCU, pI = 8.8), a digestion step with benzonase at 25 ºC was mandatory to get rid of bound RNA/DNA contamination that would otherwise strongly interfere during the subsequent purification. After an initial affinity capture step using NTA resin charged with cobalt the concentrated elute was polished by gel filtration using a Sd200/16/600 column (preparative grade).

**Sample preparation**

Frataxin variants were transferred into 50 mM HEPES Na pH 8.0, 120 mM NaCl, and 100 μM TCEP using PD-10 desalting columns packed with Sephadex G-25 resin (GE Healthcare) followed by concentration to about 300 μM using a 12 ml Vivaspin device (cutoff 10 kDa). Samples for NMR supplemented with 7 % $D_2O$ were inserted into regular 5 mm tubes. The final protein concentrations were determined spectrophotometrically at 280 nm, using extinction coefficients of 26930 $M^{-1}$ $cm^{-1}$ and 25440 $M^{-1}$ $cm^{-1}$ for the wild-type protein and the FXN-10 mutant, respectively.

Protoporphyrin IX (ppIX) was purchased from Frontier Scientific (Product Number: P562-9; CAS Number: 553-12-8).

**NMR data acquisition**

For monitoring titration with increasing amounts of either $Zn(AcO)_2$/ppIX or FeS assembly complex, a series of 2D sofast HMQC experiments[21] with band-selective $^{H}N$ excitation for fast $T_1$ recovery were acquired at 298 K on a 600 MHz Bruker Avance III and a Bruker Avance III 800 MHz spectrometer equipped with a 5 mm TXI probe and a 5 mm TCI cryoprobe, respectively. The chemical shifts of the proton-bearing carbons and their attached protons of human wild type frataxin and the FXN-10 variant were derived from 2D $^{13}C$-HSQC spectra recorded with $^{1}J_{CH}$ matched adiabatic full passage (AFP) pulses and echo/anti-echo gradients for coherence selection. All experiments were acquired at 298 K. $^{1}H$ chemical shifts were directly referenced to added DSS (2,2-dimethyl-2-silapentane-5-sulphonic acid) and $^{15}N$ chemical shifts were referenced indirectly relative to $^{1}H$ using IUPAC ratios (https://bmrb.io/ref info/cshift.shtm). All NMR data were processed with NMRpipe[22] and analyzed with NMRFAM-Sparky.[23]

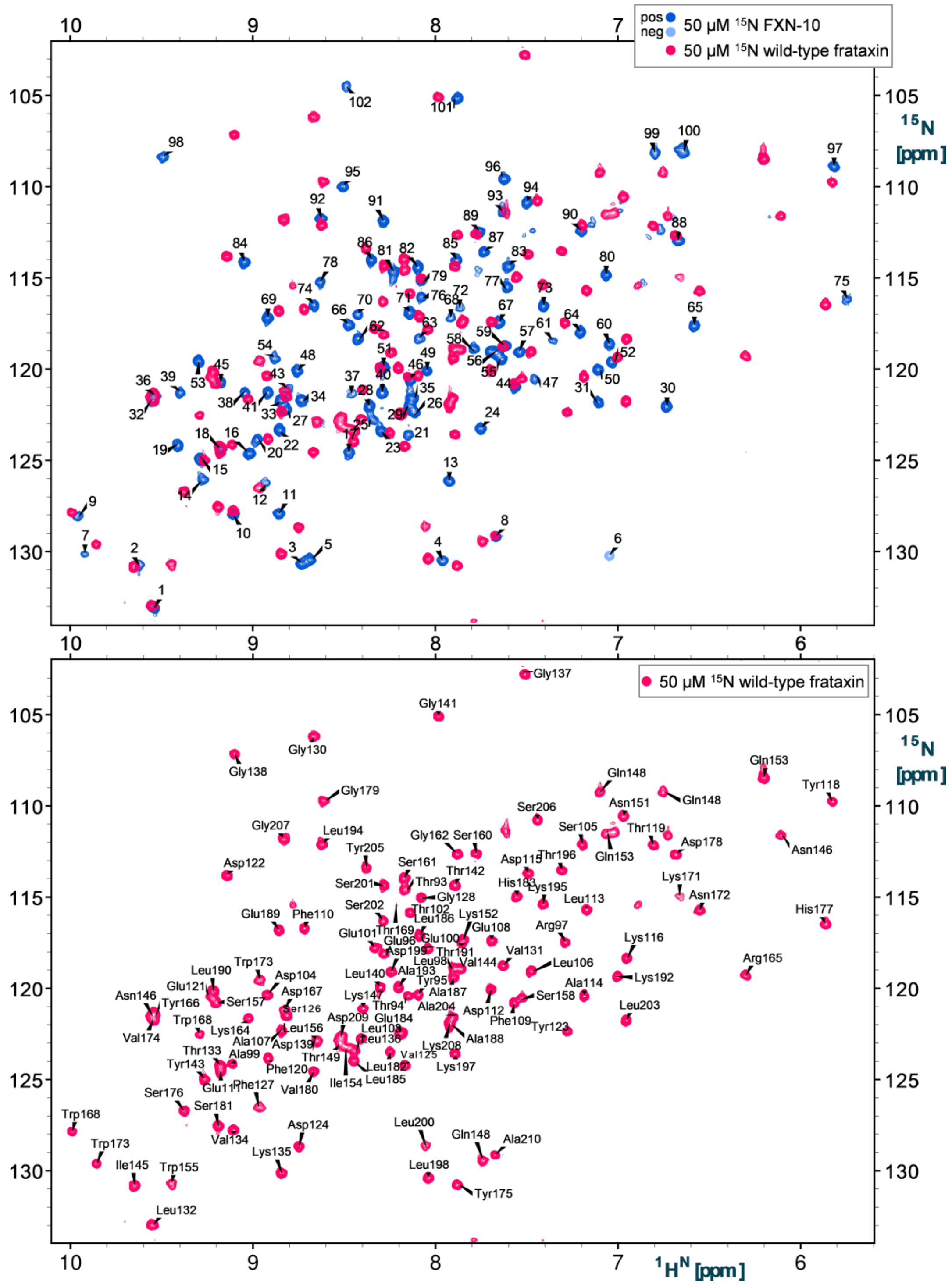


**Figure S12.** 600 MHz 2D $^{15}$N-sf-HMQC spectra of u-$^{15}$N,$^{13}$C-labelled FXN-10 and u-$^{15}$N labelled wild-type frataxin. Top: overall, the chemical shift coordinates of both proteins are quite different due to presence of 13 single-residue mutations in FXN-10. Therefore, $^{1}H^{N}/^{15}N$ resonances are not unambiguous assigned for FXN-10 and all peaks are labelled with arbitrary numbers. Bottom: 2D $^{15}$N-sf-HMQC spectra with backbone amide and sidechain amide assignments (for indol $H_{\varepsilon1}/N_{\varepsilon}$, Asn $H_{\gamma1,2}/N_{\gamma}$ and Gln $H_{\delta1,2}/N_{\delta}$, respectively) of wild-type frataxin.

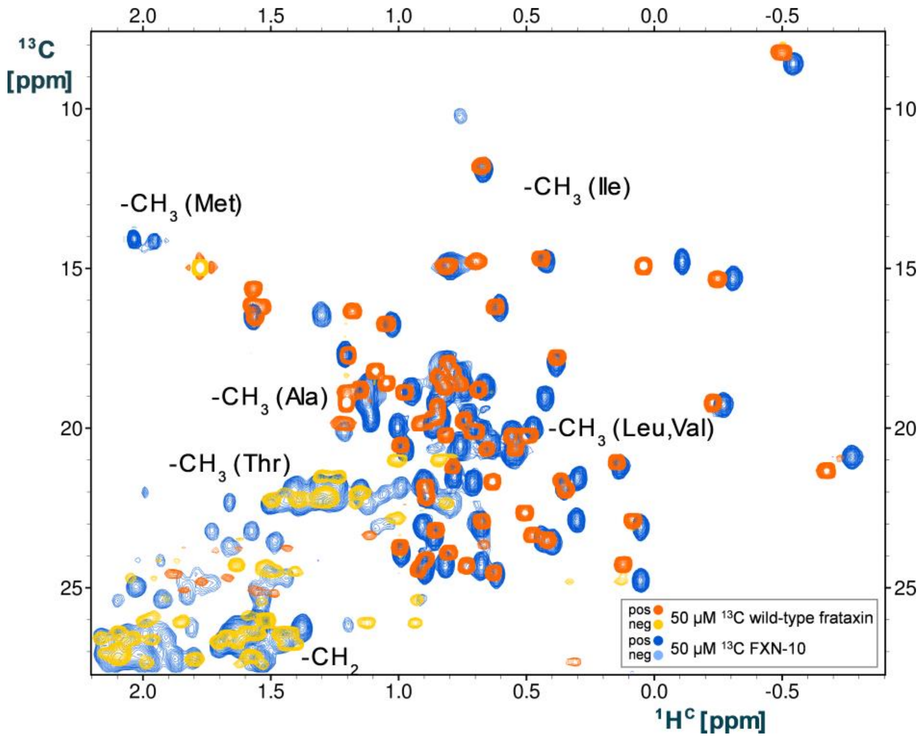


**Figure S13.** 800 MHz 2D $^{13}$C-HSQC spectra of FXN-10 and constant time (ct) multiplicity-edited 2D $^{13}$C-HSQC spectra of wild-type frataxin (methyl groups region). Despite the presence of 13 mutations in FXN-10, the resonances for the methyl groups superimpose reasonably well, suggesting that the packing of the protein core is very similar.

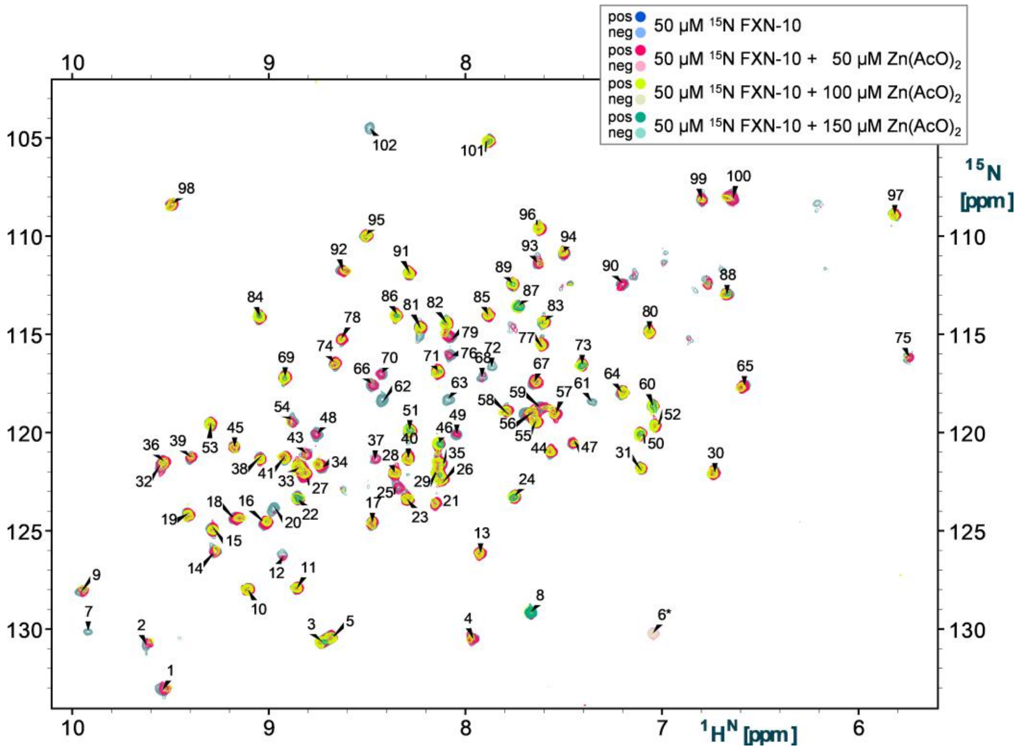


**Figure S14.** 600 MHz 2D $^{15}$N-sf-HMQC spectra of u-$^{15}$N,$^{13}$C-labelled FXN-10 (arbitrary residue numbers due to lack of assignment) in the presence of increasing $Zn^{2+}$ concentrations.

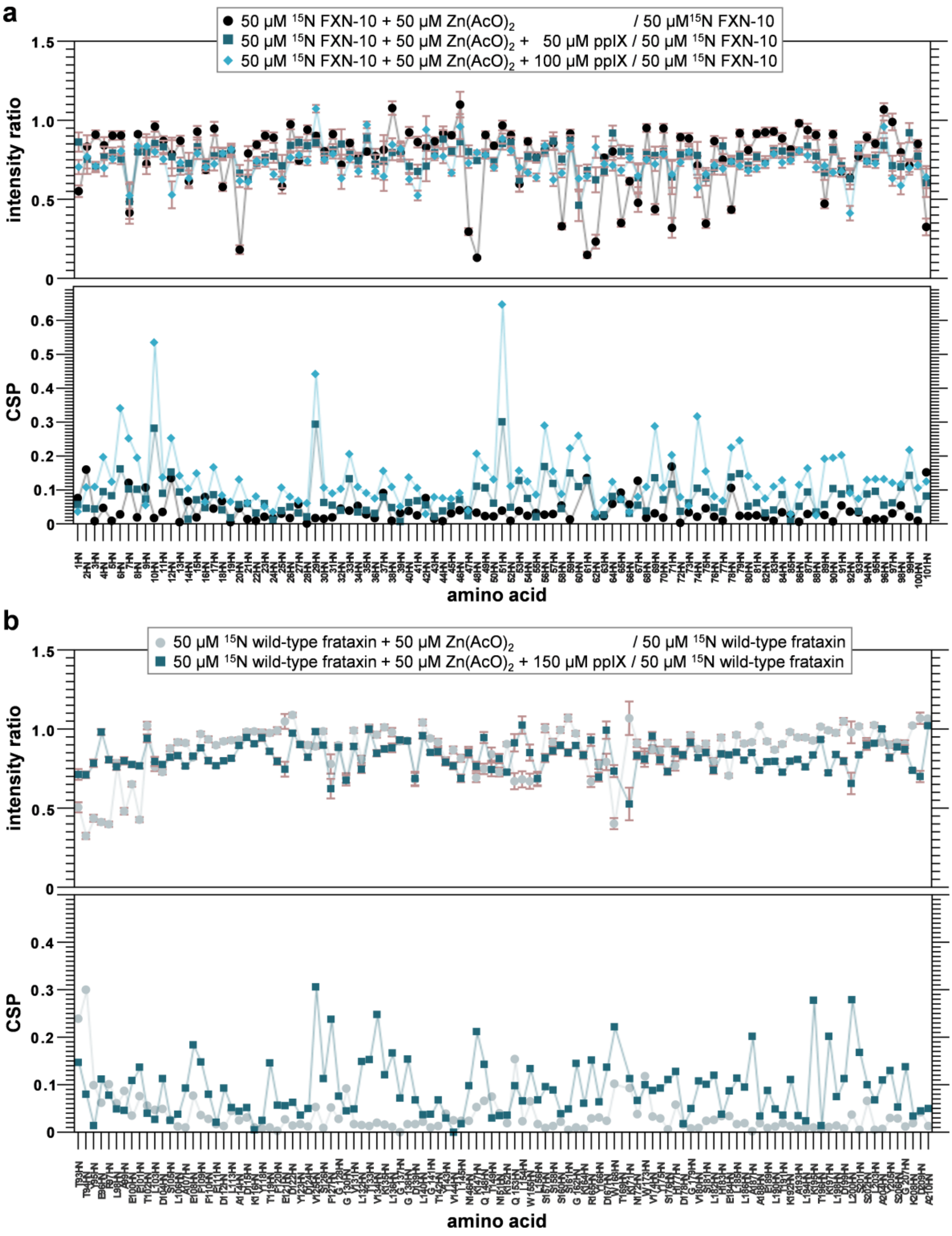


**Figure S15.** Signal intensity ratios and chemical shift perturbations (CSP) observed for a) u-$^{15}N$,$^{13}C$-labelled FXN-10 (arbitrary residue numbers due to lack of assignment), and b) u-$^{15}N$-labelled wild-type frataxin in the presence of $Zn^{2+}$ and protoporphyrin IX (ppIX). Top: signal intensity ratios extracted from 2D $^{15}N$-sf-HMQC spectra. Bottom: CSP plot shows the characteristic changes found at the C-terminus of helix α1 upon addition of a binder and indicates $Zn^{2+}$/ppIX binding.

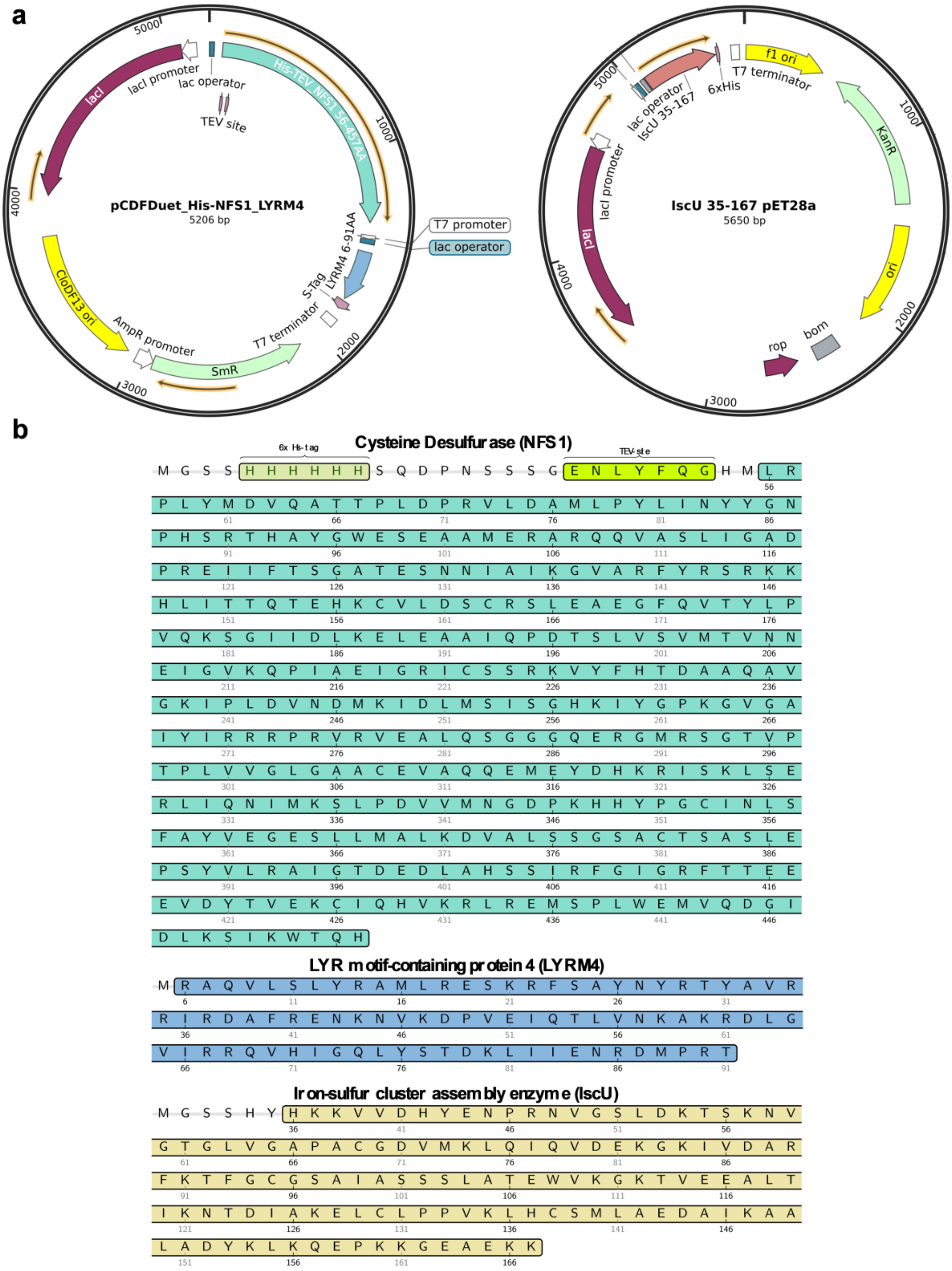


**Figure S16**. Constructs for the human mitochondrial iron-sulfur cluster assembly proteins expressed and purified. a) Schematic map of plasmids used. b) Primary sequences of expressed protein constructs.

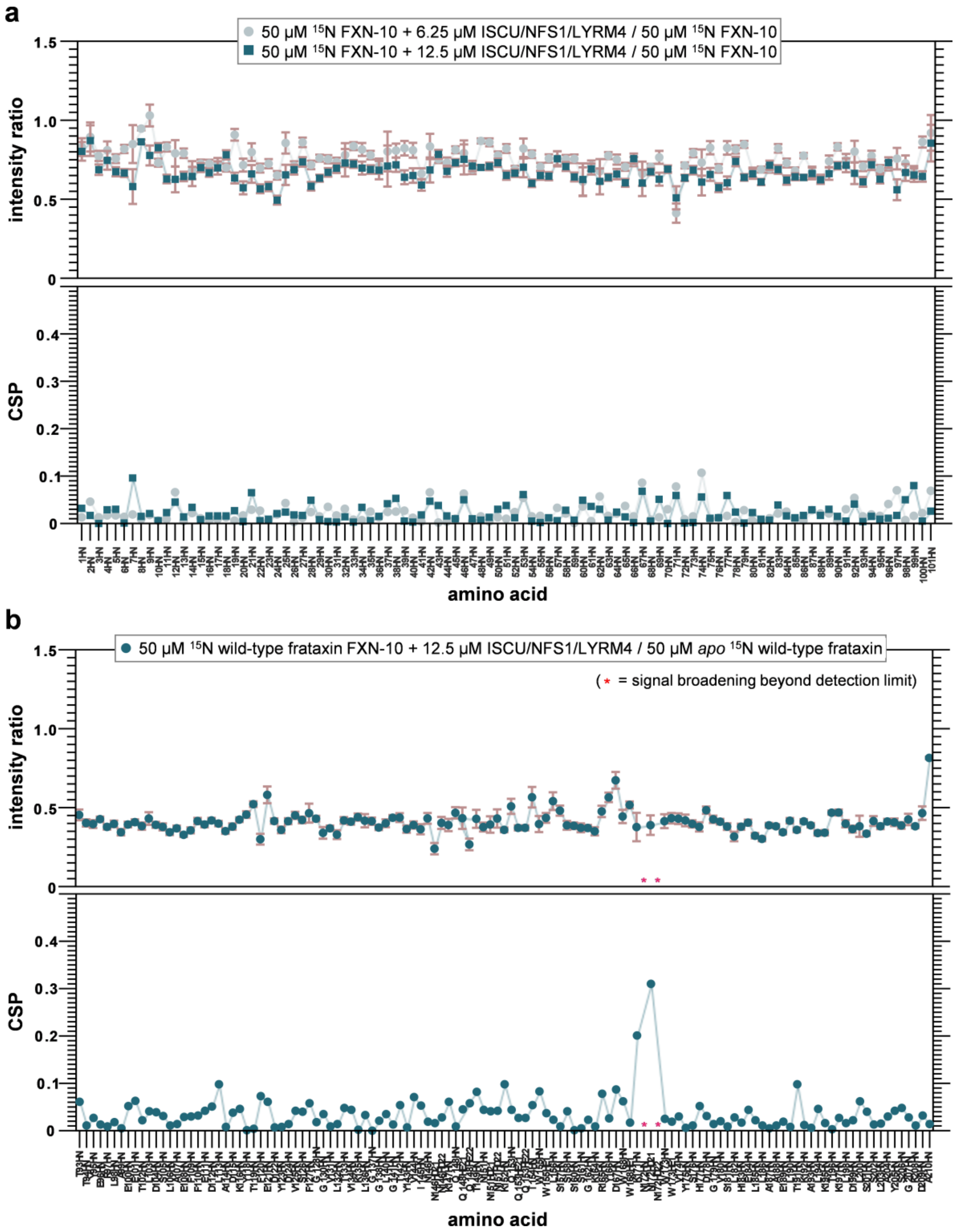


**Figure S17.** Signal intensity ratios and chemical shift perturbations (CSP) observed for a) u-$^{15}$N,$^{13}$C-labelled FXN-10 (arbitrary residue numbers due to lack of assignments), and b) u-$^{15}$N-labelled wild-type frataxin in the presence of FeS assembly complex. Top: signal intensity ratios extracted from 2D $^{15}$N-sf-HMQC spectra. Bottom: consistent with the slow exchange signature of the spectra, no changes of the chemical shifts are observed, except around residue K171 in the wild-type. This residue (substituted by Thr in FXN-10) stablishes transient interactions with N172 that might be differently populated in the presence of the FeS assembly complex.

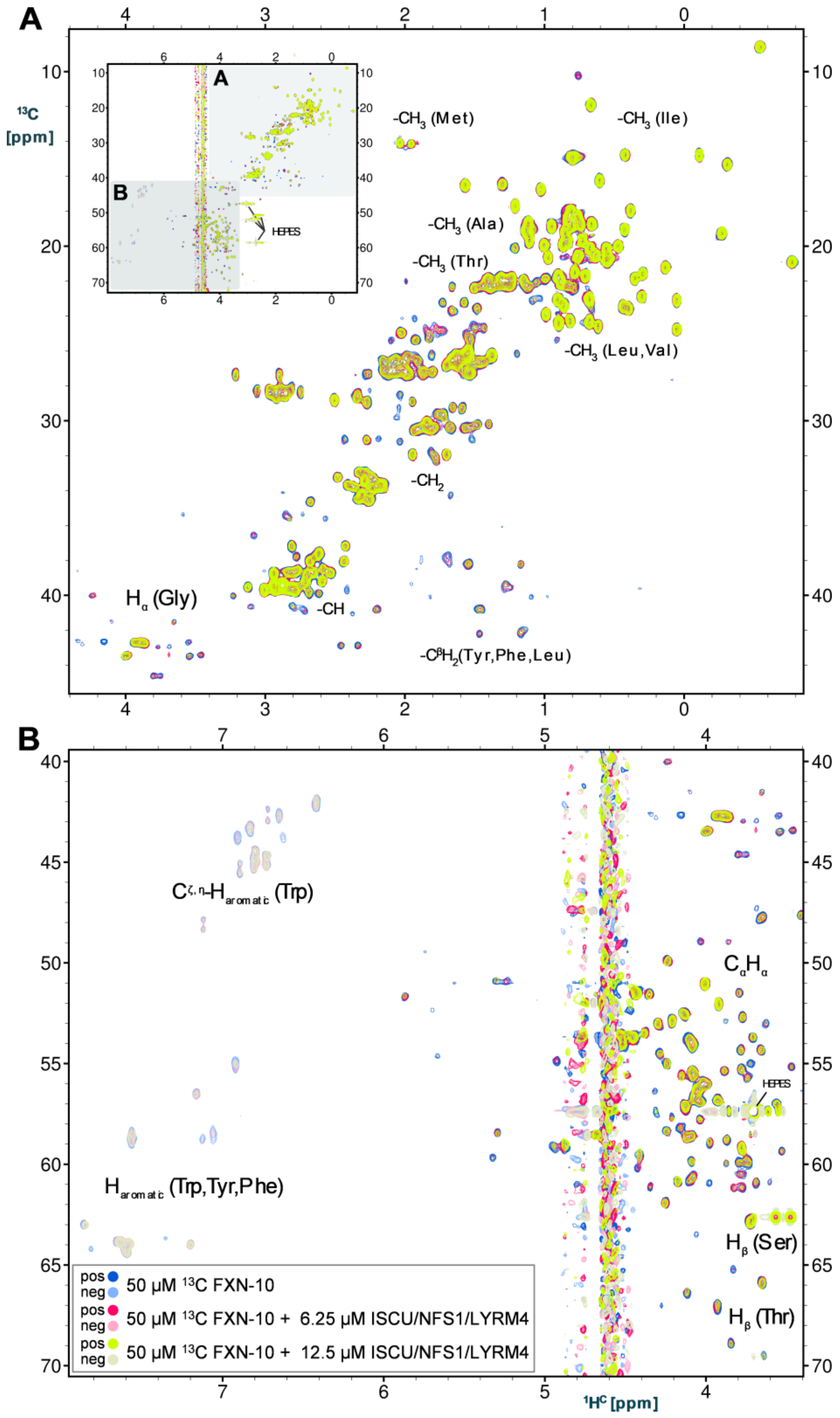


**Figure S18.** 800 MHz 2D non-constant time $^{13}$C-HSQC spectra of u-$^{15}$N,$^{13}$C-labelled FXN-10 in the presence of increasing concentrations of the FeS assembly complex. Regions of typical chemical shifts for A) sidechain methine, methylen and methyl moieties as well as B) Hα and aromatic proton-bearing carbons, are displayed. The complete spectra are shown in the top left insert. The intensities of the C–$H_{x=1\text{-}3}$ resonances clearly decrease upon ISCU/NFS1/LYRM4 addition without any alteration of their chemical shift coordinates, evidencing binding to the FeS assembly complex.

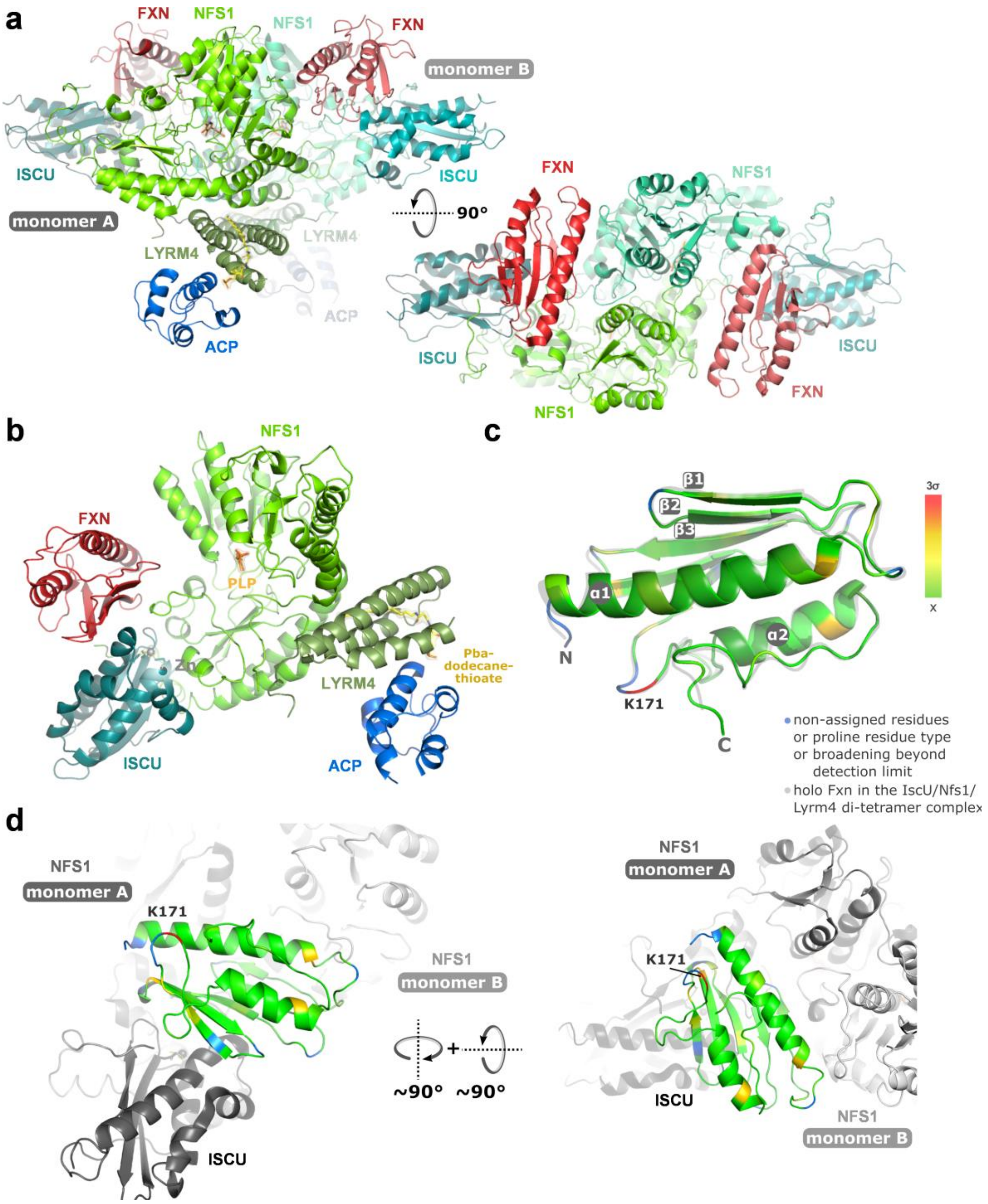


**Figure S19.** A) High resolution crystal structures of the homodimeric human FXN/NFS1/ISD11/ACP/ISCU-$Zn^{2+}$ complex (PDB ID 6NZU). Frataxin (FXN) is located between the ISCU and NFS1protomers. B) Each monomer contains a $Zn^{2+}$ ion bound to ISCU, a pyridoxalphosphate (PLP) cofactor for the cysteine desulfurase NFS1, and a long chain fatty acid (40-phosphopantetheine) of ACP inserting into the helical center of the LYRM4 subunit. Mapping of backbone chemical shift perturbations (CSP) onto the NMR solution model (C) and to bound frataxin in the FeS assembly complex (D) locates only effected site around residue K171 located in the flexible loop β5/α2 facing outwards into the solvent. The color code is proportional to the standard deviation scale from the average value, as indicated in the bar legend (x: average value, in green).
